# Transfer RNA acetylation regulates in vivo mammalian stress signaling

**DOI:** 10.1101/2024.07.25.605208

**Authors:** Supuni Thalalla Gamage, Roxane Khoogar, Shereen Howpay Manage, McKenna C. Crawford, Joe Georgeson, Bogdan V. Polevoda, Chelsea Sanders, Kendall A. Lee, Kellie D. Nance, Vinithra Iyer, Anatoly Kustanovich, Minervo Perez, Chu T. Thu, Sam R. Nance, Ruhul Amin, Christine N. Miller, Ronald J. Holewinski, Thomas Meyer, Vishal Koparde, Acong Yang, Parthav Jailwala, Joe T. Nguyen, Thorkell Andresson, Kent Hunter, Shuo Gu, Beverly A. Mock, Elijah F. Edmondson, Simone Difilippantonio, Raj Chari, Schraga Schwartz, Mitchell R. O’Connell, Colin Chih-Chien Wu, Jordan L. Meier

## Abstract

Transfer RNA (tRNA) modifications are crucial for protein synthesis, but their position-specific physiological roles remain poorly understood. Here we investigate the impact of N4-acetylcytidine (ac^4^C), a highly conserved tRNA modification, using a Thumpd1 knockout mouse model. We find that loss of Thumpd1-dependent tRNA acetylation leads to reduced levels of tRNA^Leu^, increased ribosome stalling, and activation of eIF2α phosphorylation. Thumpd1 knockout mice exhibit growth defects and sterility. Remarkably, concurrent knockout of Thumpd1 and the stress-sensing kinase Gcn2 causes penetrant postnatal lethality, indicating a critical genetic interaction. Our findings demonstrate that a modification restricted to a single position within type II cytosolic tRNAs can regulate ribosome-mediated stress signaling in mammalian organisms, with implications for our understanding of translation control as well as therapeutic interventions.

## INTRODUCTION

Transfer RNAs (tRNAs) are fundamental to protein synthesis, serving as the physical connection between amino acids and their corresponding nucleotide triplets as defined by the Genetic Code. A conspicuous trait of these molecules observed across all organisms is their decoration by >100 structurally diverse RNA modifications.^1–3^ Metazoan tRNAs contain on average ∼13 modified nucleotides per molecule. However, these modifications are not distributed evenly, with some reaching near ubiquity and others being quite rare, found on only a small subset of tRNA isoacceptors. Prominent among this latter more restricted class is RNA cytidine acetylation.

N4-acetylcytidine (ac^4^C) is a highly conserved RNA modification found across all domains of life (Fig. 1a).^4^ In humans and mice ac^4^C is introduced into RNA posttranscriptionally by the essential enzyme N-acetyltransferase 10 (NAT10).^5, 6^ One unique property of this enzyme is its ability to modify both tRNA and ribosomal RNA (rRNA). To address these distinct substrates Nat10 uses specific adapters, which for tRNA include THUMP domain-containing protein 1 (Thumpd1). The role of THUMP domain proteins in ac^4^C biogenesis was first discovered in *S. cerevisiae*, where genetic screens found deletion of a Thumpd1 homologue (Tan1) causes synthetic lethality in strains harboring a mutated allele of the single copy tRNA^Ser^_CGA_ gene.^7, 8^ Thumpd1 was found to be required for formation of ac^4^C at position C12 in the D-arm of the tRNA^Ser^ and tRNA^Leu^. Subsequent studies found loss of ac^4^C reduced the levels of these type II tRNAs in *S. cerevisiae* (tRNA^Ser^)^9^ and more recently *S. pombe* (tRNA^Leu^)^10^ and can cause growth defects at higher temperatures, consistent with its biophysically stabilizing role.^11^ Disruption of tRNA ac^4^C in yeast can also activate the general amino acid control pathway,^10^ suggestive of a connection to ribosome-mediated activation of Gcn2.^12–14^ In humans, rare bi-allelic mutations in *THUMPD1* are associated with loss of ac^4^C in tRNA as well as neurodevelopmental defects.^15^

**Figure 1.**
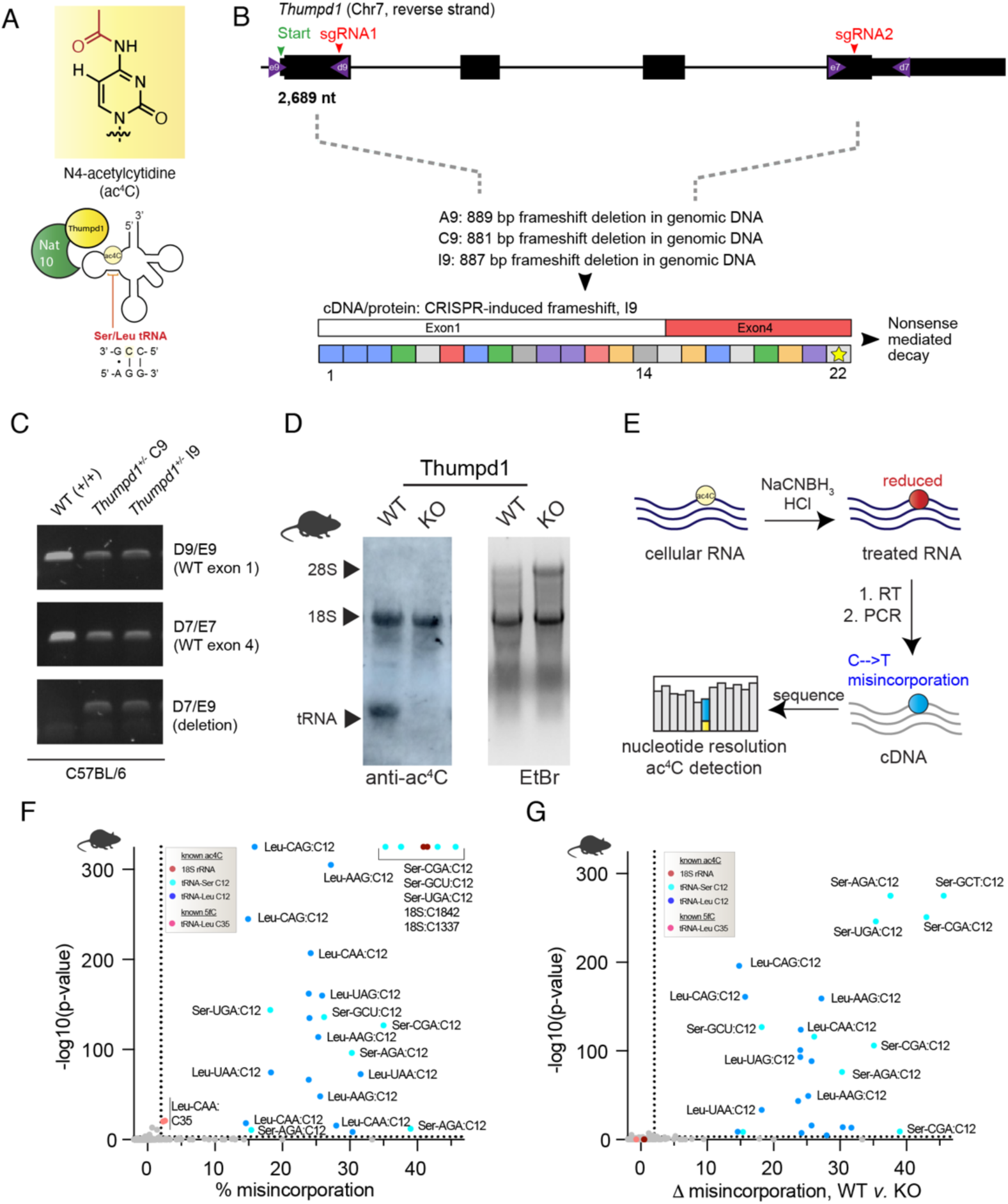
An *in vivo* model for studying mammalian transfer RNA acetylation. (a) Deposition of the RNA modification N4-acetylcytidine (ac^4^C) in eukaryotic transfer RNA requires Nat10 and Thumpd1. (b) Schematic of murine *Thumpd1* locus and CRISPR-Cas9 genome editing strategy. Multiple murine lines (A9, C9, and I9) were obtained and genotypes characterized by next-generation sequencing. (c) PCR-based genotyping confirms *Thumpd1* deletion. (d) Immuno-Northern blotting confirms loss of ac^4^C in tRNA upon Thumpd1 knockout. (e) Schematic for ac^4^C sequencing (ac^4^C-Seq). (f) Distribution of ac^4^C in unfractionated murine tRNA-enriched total RNA. Data represent mean values from three independent mouse liver samples (n=3 biological replicates). (g) Applying ac^4^C-seq to unfractionated total RNA isolated from wild-type (WT) or Thumpd1 KO mice highlights C12 in the D-arm of tRNA^Leu/Ser^ as the dominant position of Thumpd1-regulated RNA acetylation. Data represent mean values from three independent mouse liver samples (n=3 biological replicates per genotype).

A paradox in RNA modification biology is that even as the number of putative RNA modification sites known has rapidly increased, analyses of individual positions and their physiological impacts remain rare.^3^ In models like yeast RNA modification mutants often exhibit subtle phenotypes, motivating their dissection in multicellular organisms, where more pronounced and diverse phenotypes have the opportunity to present themselves.^16–18^ Towards this goal, here we investigate the physiological effects of transfer RNA ac^4^C in a mouse model of Thumpd1 loss. Using ac^4^C -seq, we find that across evolution, the dominant sites of tRNA acetylation are tightly confined to the D-arm of tRNA^Ser^ and tRNA^Leu^ isodecoders. Thumpd1 knockout mice, which lack tRNA acetylation, are viable but show sub-Mendelian birth ratios, exhibit growth defects, and fail to reproduce. Characterizing a human THUMPD1 knockout cell line yields evidence that loss of tRNA acetylation is associated with reduced levels of tRNA^Leu^, ribosome stalling, and ribosome collisions. This stimulates phosphorylation of the translation factor eIF2α, causing a general defect in protein synthesis. Similar evidence for reduced tRNA^Leu^ and eIF2α phosphorylation is observed in cerebella isolated from Thumpd1 KO mice. Finally, we show that dual knockout of Thumpd1 and Gcn2 causes penetrant post-natal lethality, linking tRNA acetylation to eIF2α kinase activity *in vivo*. Overall, our findings demonstrate the ability of a restricted and non-essential tRNA modification to regulate signaling through the ribosome in a multicellular organism, with implications for human physiology and disease.

## RESULTS

### In vivo manipulation of tRNA acetylation via Thumpd1

To develop an *in vivo* model for studying mammalian transfer RNA acetylation we constructed a heterozygous knockout mouse (*Thumpd1^+/-^*) by targeting sites in exons 1 and 4 of the *Thumpd1* gene located on chromosome 7 using CRISPR-Cas9. Next-generation sequencing of the *Thumpd1* locus revealed the formation of large (∼880 base pair) frameshift deletions in several lines, which could be readily differentiated from the wild-type allele by PCR (Fig. 1b-c). To measure the expression of Thumpd1, we analyzed Thumpd1 mRNA levels in liver isolated from wild-type (WT), heterozygous (*Thumpd1^+/−^*), and knockout (KO, *Thumpd1*^-/-^) variants. Thumpd1 transcript levels directly correlated with the presence of an intact allele, with WT mice expressing the highest and KO mice the lowest amount (Fig. S1a). Anti-ac^4^C immunoblotting confirmed loss of tRNA acetylation (lower band) but not rRNA acetylation (top band) in the KO line (Fig 1d).

Next, we sought to better understand the spectrum of tRNAs marked by Thumpd1-dependent cytidine acetylation. In our previous studies - as well as those of others – ac^4^C has been identified exclusively at C12 within the D-arm of eukaryotic tRNA^Leu/Ser^.^7, 19^ However, it is not known whether ac^4^C is a pervasive characteristic of tRNA^Leu/Ser^ or marks only a subset of isoacceptors and/or isodecoders. Furthermore, a few recent studies have suggested a broader role for ac^4^C within tRNAs.^20–22^ To characterize this, we characterized the presence of cytidine acetylation in eukaryotic tRNAs using ac^4^C-seq (Fig. 1e). This quantitative sequencing reaction uses acidic NaCNBH_3_ to alter the structure of ac^4^C, resulting in misincorporations at modification sites upon reverse transcription.^23^ These can then be identified as mutations by complementary DNA (cDNA) sequencing. The quantitative nature of ac^4^C-seq as well as its value in discovery applications has been validated across multiple settings.^19, 24^ Pooling three replicate ac^4^C-seq analyses of tRNA-enriched total RNA isolated from livers of wild-type and Thumpd1 KO mice, we sampled over 9,603 tRNA positions at depths of >100 reads, including over 75% of known tRNA^Ser^ and tRNA^Leu^ genes. NaCNBH_3_-dependent misincorporations were observed across all tRNA^Ser^ and tRNA^Leu^ genes sampled, which summed 26 in total and included all 9 isoacceptors (Fig. 1f, Supplementary Table 1a). In all cases modifications occurred at C12. Misincorporation signals were overall more penetrant for tRNA^Ser^ than tRNA^Leu^. However, careful inspection of the sequence coverage identified a higher propensity for stops adjacent to C12 of tRNA^Leu^ (Fig. S1b). This is an important illustration that while ac^4^C-seq is quantitative for individual sites it is not absolute; misincorporation signals across sites may vary based due both to reduced ac^4^C stoichiometry as well as each individual RNA’s propensity for stop versus readthrough. Strong misincorporation signals were also observed at C35 of tRNA^Leu^_CAA_, a known 5-formylcytidine (f^5^C) site. We have previously defined the cross-reactivity of f^5^C with the ac^4^C- seq chemistry.^25^ Other sites producing significant C > T signals upon acidic NaCNBH_3_ treatment were characterized by low (<2%) misincorporation ratios. In addition to their low stoichiometry, these sites did not occur at the 5’-CCG-3’ consensus sequence whose modification Nat10 is known to favor and were not sensitive to Thumpd1 KO (Fig. 1g, Supplementary Table S1b). To see if we could identify additional penetrant cytidine acetylation sites in tRNA we used ac^4^C-seq to analyze small RNA fractions isolated from additional yeast, mouse, and human models. Here again, we found modification sites conserved across models to be confined to tRNA^Leu/Ser^ C12 (Fig. S1c-e, Supplementary Table S1c). We further corroborated that C12 tRNA^Ser^_CGA_ acetylation is abolished in Thumpd1 KO mice using amplicon sequencing (Fig. S1f). Taken as a whole, our results indicate that the most penetrant sites of tRNA acetylation are confined to the D-arm of tRNA^Ser^ and tRNA^Leu^ across eukaryotes, can occur across all isoacceptors, and are highly dependent on the presence of Thumpd1.

### Thumpd1 loss in mice is associated with small size and sterility

Heterozygous *Thumpd1*^+/−^ mice are viable and fertile and do not present any overt phenotype. However, mice completely deficient in Thumpd1 *(Thumpd1^-/-^)* have significantly smaller body sizes, suggesting an important role for this gene in growth (Fig. 2a). Despite their small size, adult *Thumpd1^-/-^* mice are viable as assessed by maintenance of body weight over time (Fig. 2b). Breeding studies found *Thumpd1^-/-^* mice are born at sub-Mendelian ratios indicative of decreased pre- or perinatal fitness (Fig. 2c). Ovarian atrophy in KO mice is characterized histologically by decreased ovarian oocytes, developing follicles, and corpora lutea and replacement with interstitial cell hyperplasia (Fig. 2d). Testicular atrophy is characterized histologically by disorganized seminiferous tubules with loss or vacuolation of germ cells and multinucleated germ cells; exfoliated germ cells are also observed within the epididymis (Fig. 2e). Consistent with this observation, repeated matings of male *Thumpd1^-/-^* mice (6-21 weeks of age) with WT females, and female *Thumpd1^-/-^* mice (6-21 weeks of age) with WT males failed to produce pregnancies. In contrast to the neurodevelopmental defects observed in human patients,^15^ no obvious changes in brain weight or morphology were observed in KO mice. These findings define overt phenotypes that accompany loss of Thumpd1-regulated tRNA acetylation in an animal model.

**Figure 2.**
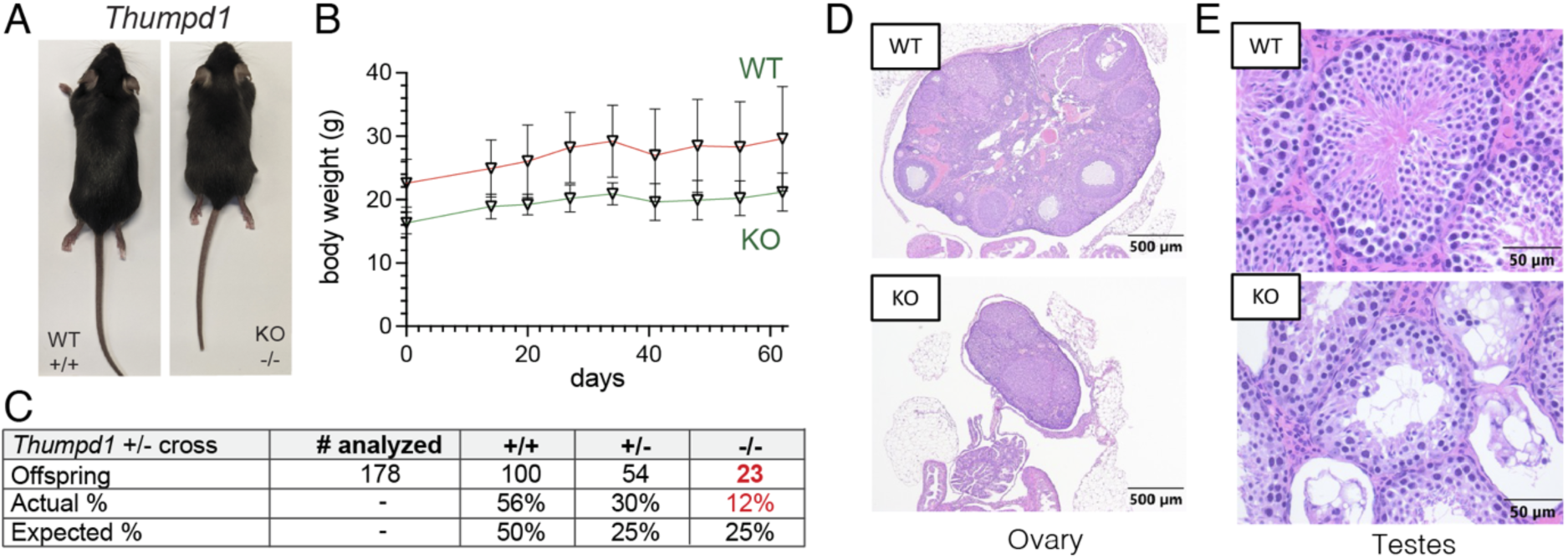
*In vivo* phenotypes associated with loss of Thumpd1 and tRNA acetylation. (a) *Thumpd1^-/-^*mice are runted. (b) *Thumpd1^-/-^* mice are viable for several weeks. (c) Offspring produced in *Thumpd1^+/-^* breeding studies. (d) Ovarian atrophy and (e) testicular seminiferous tubule degeneration in *Thumpd1^-/-^* mice. Immunohistochemistry results are representative of n=4 biological replicates per genotype.

### THUMPD1 regulates overall levels and ribosome occupancy of tRNA^Leu^

As a model system for mechanistic studies, we used CRISPR-Cas9 to delete THUMPD1 in an immortalized human HEK-293T cell line. Our rationale was that working in human cells might highlight commonalities shared across eukaryotes, while moreover providing a model technically tractable to analysis by a wide range of experimental techniques. Similar to observations in mice, targeting of the *THUMPD1* gene using CRISPR-Cas9 resulted in loss of THUMPD1 protein as well as specific abrogation of tRNA acetylation (Fig. 3a-b). Previous studies have found deletion of the *S. cerevisiae* Thumpd1 homologue decreased levels of ac^4^C-containing tRNAs.^9, 26^ Therefore, we performed unbiased profiling of tRNAs in our wild-type and KO models using modification-induced misincorporation tRNA sequencing (mim-tRNA-Seq).^27^ This revealed that THUMPD1 KO specifically affects the levels of ac^4^C-containing tRNAs in HEK-293T cells, causing an approximate 4-fold decrease in levels of tRNA^Leu^_UAA-2_ and smaller reductions in tRNA^Ser^_AGA-3_, tRNA^Ser^_UGA-4_, tRNA^Leu^_AAG-3_, and tRNA^Leu^_UAA-3_ (Fig. 3c, Supplementary Table S2) To define how these perturbations impacted translation we performed ribosome profiling in THUMPD1 WT and KO cells. Our data showed that loss of THUMPD1 results in increased A-site occupancy at UUG, CUA, CUU, and UUA codons (Fig. 3d, Supplementary Table S3). Invoking wobble rules, all four of these can be decoded by tRNA^Leu^_UAA_ and tRNA^Leu^_AAG_ (more often found as tRNA^Leu^_IAG_).^28^ Both of these species are detected as downregulated in our mim-tRNA-Seq data. Our data contrasts with that of Darnell and coworkers, who found ribosomes were recalcitrant to stalling at Leu codons even upon leucine deprivation^29^ and better matches the observation of Chou et al., who detected altered ribosome occupancy at Leu codons upon deletion of the *S. cerevisiae* Thumpd1 homologue.^30^ Global proteomics as well as immunoblotting indicated that genes with decreased translation efficiency (TE) in THUMPD1 KO cells were modestly downregulated at the protein level (Fig. 3e, Supplementary Table S4). Surveys of transcript level data for mRNAs such as PSMB4 (Fig. 3f) and CDK1 (Fig. S2) indicate that even in transcripts where strong pauses at tRNA^Leu^-decoded codons can be visualized, ribosome occupancy is still observed downstream. This suggests that even when THUMPD1-dependent pausing occurs at Leu codons, protein production may be unburdened.

**Figure 3.**
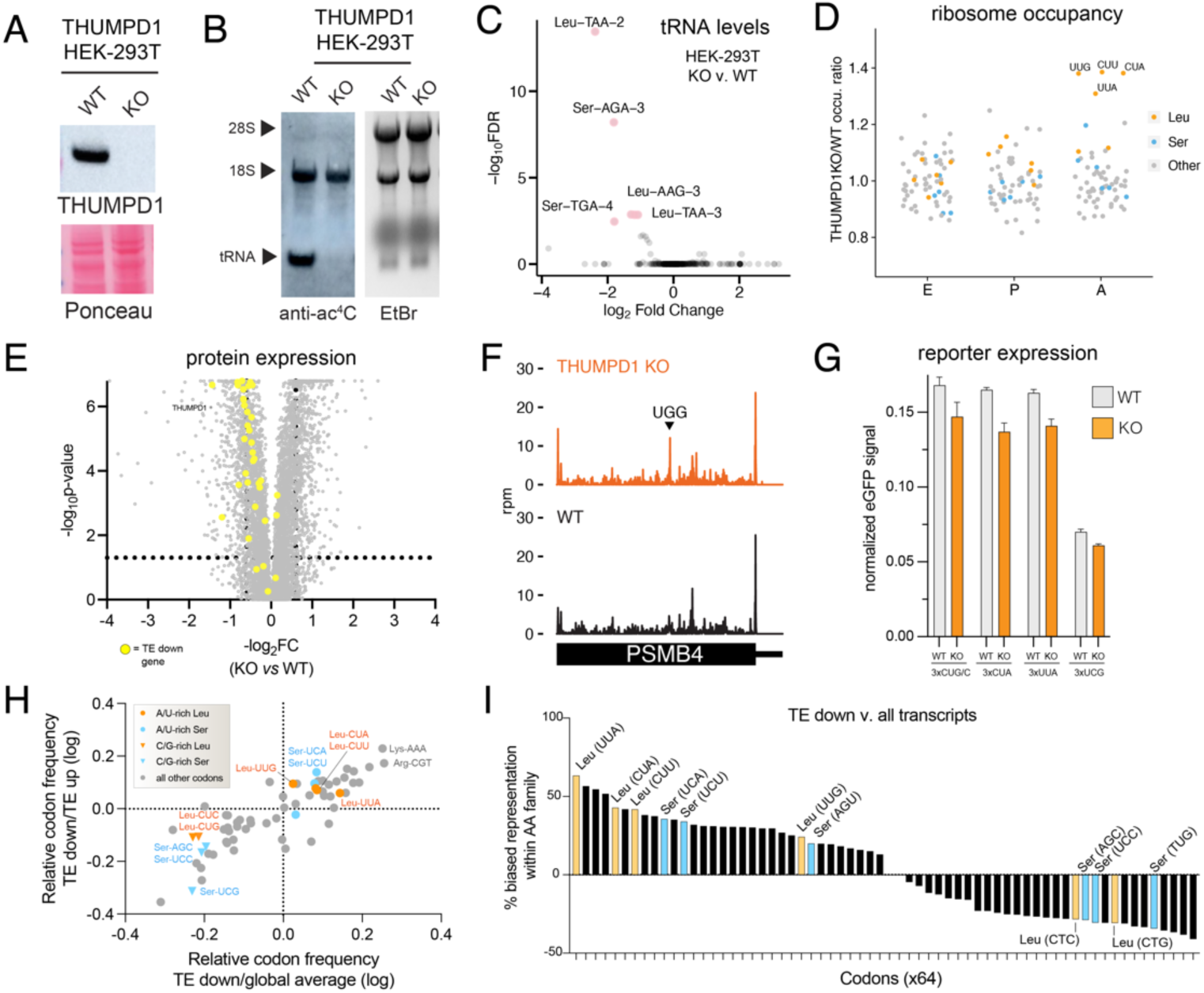
Molecular characterization of THUMPD1/tRNA acetylation loss in a mammalian cell line. (a) Western blot confirms THUMPD1 KO. Data are representative of n=3 biological replicates. (b) Immuno-Northern blotting confirms loss of ac4C in tRNA upon THUMPD1 knockout in HEK-293T cells. Data are representative of n=2 biological replicates. (c) mim-tRNA-Seq analysis of WT v. THUMPD1 KO cells indicates decreased levels of tRNA^Leu/Ser^ isodecoders. Data are derived from n=2 biological replicates. (d) Ribosome profiling of WT v. THUMPD1 KO cells at the E, P, and A sites indicate increased ribosome pausing at codons (UUG, CUU, CUA, and UUA) decoded by tRNA^Leu^ in the ribosomal A site. Codons encoding Leu, Ser, and other amino acids are shown in blue, orange, and grey, respectively. Data are derived from n=4 biological replicates. (e) Proteomic profiling of WT v. THUMPD1 KO cells. Genes showing decreased translation efficiency (TE down) are highlighted in yellow. Values are derived from n=3 biological replicates. (f) Ribosome footprints in PSMB4 gene indicates that stalling at Leu codons in KO cells does not limit downstream ribosome occupancy. (g) eGFP reporters with multiple (3x) copies of Leu or Ser codons do not show significantly decreased protein production in THUMPD1 KO cells. Data are derived from n=3 technical replicates. (h) Scatterplot of codon frequency changes mRNAs differential translated in Thumpd1 KO cells. Glbal average refers to the average codon usage of all CCDS-defined consensus coding sequences. Values shifted up and to the right indicate codons more frequent in TE down transcripts. Leu codons = orange, Ser codons = blue. (i) Analysis of amino acid family-specific codon bias in TE down transcripts. The representation of each individual codon (e.g. Leu-UUA) relative to its amino acid family (e.g. all Leu) was calculated. Average values for TE down sequences and all CCDS-defined consensus coding sequences were compared to assess representation. Leu codons = orange, Ser codons = blue, U/A-rich Leu codons are labeled on the x-axis in red.

One way tRNA modifications can shape the proteome is by influencing codon-biased translation.^31–36^ To test whether a similar phenomenon acted in THUMPD1 KO cells, we performed comparisons using a reporter containing multiple copies of Leu codons that exhibited stalling (UUA and CUA) or which didn’t exhibit stalling (CUG/C and UCG) upstream of an eGFP gene. No significant difference in protein production was observed relative to WT cells (Fig. 3g). To explore this phenomenon further we analyzed codon composition in genes showing altered TE in THUMPD1 KO cells (Supplementary Table S5). Comparing canonical transcripts found in the TE up and TE down datasets (Supplementary Table S5), we found the latter contained a significantly greater percentage of the U/A-rich Leu codons that showed stalling in our ribosome profiling studies (UUG, CUA, CUU, and UUA) relative to C/G-rich Leu codons that did not show stalling (CUC and CUG; Fig. 3h y-axis, Fig. S3a). Compared to codon composition of all human genes U/A-rich Leu codons were also modestly enriched (Fig. 3h, x-axis, Fig. S3b-c). Analysis of amino acid pairs within TE down transcripts also found modest enrichment of di-amino acids containing U/A-rich Leu (Supplementary Table S5).^37^ Evaluating the skew between representation of individual codons (e.g. Leu-UUA) relative to the entirety of their amino acid family (e.g. all Leu) in TE down coding sequences revealed a clear bifurcation between U/A- and C/G-rich Leu/Ser (Fig. 3i, Fig. 3d). The former are more enriched TE down mRNAs, offering additional evidence of U/A-rich Leu/Ser bias. Interestingly, while TE down sequences exhibit a shift in Leu codon composition favoring U/A-rich variants, other codons such as Lys-AAA and Arg-CU exhibit the highest absolute frequencies of enrichment (Fig. 3h). This suggests that context, rather than frequency, determines the effect of Thumpd1 KO on protein translation. As one example, association of positively charged amino acids with the ribosomal exit tunnel can decrease translational velocity^38^ and cause premature termination,^39, 40^ raising the possibility that THUMPD1 may collaborate with this or other transcript features. Our results paint a picture that requires holding two opposing ideas in mind: first, sequence biases exist that imply - but do not demonstrate - a subtle, context-dependent impact of THUMPD1 on codon-specific translation. Second, the magnitude of this effect appears insufficient to disrupt reporter gene expression or obviously account for the extensive remodeling of HEK-293T proteomes caused by loss of tRNA ac^4^C.

### THUMPD1 loss stimulates phosphorylation of eIF2 alpha

As an alternative to codon-specific effects, we considered what signaling pathways may help cells maintain homeostasis in response to reduced tRNA^Leu/Ser^ levels. The integrated stress response (ISR) is major cellular mechanism used to adapt to changing environmental conditions including nutrient deprivation, pathogens, heme deficiency, and endoplasmic reticulum (ER) stress.^41^ Uncharged tRNAs and/or ribosomal collisions activate the sensor kinase Gcn2, which in turn phosphorylates the translation factor eIF2α at Ser51 (Fig. 4a). This phosphorylation event promotes formation of a tight eIF2α/eIF2β complex incapable of the nucleotide recycling necessary for canonical cap-dependent translation. Instead, proteins are produced primarily from mRNAs whose translation is recalcitrant to this event such as the stress-responsive transcription factor ATF4. As previous studies have found uncharged tRNAs to be more rapidly degraded than their charged counterparts,^42^ we hypothesized that the lower levels of tRNA^Ser^ and tRNA^Leu^ in THUMPD1 KO cells may reflect reduced aminoacylation. However, applying both qRT-PCR and hybridization-based assays we did not observe significant evidence for decreased charging of tRNA^Leu^ in THUMPD1 KO cells (Fig. S4a-b). This is consistent with a recent study which found deletion of the yeast Thumpd1 homologue does not affect tRNA charging.^43^ To probe for evidence of ribosome collisions, we applied disome profiling to profile mRNA fragments differentially protected by endogenous stacked ribosomes in the wild-type and THUMPD1 mutant cell lines. High resolution footprinting revealed increased A-site occupancy across several codons in THUMPD1 KO cells (Fig. 4b-d, Supplementary Table S6). While amino acid-selective occupancy changes were not as pronounced as with monosome analyses, the greatest differences were observed at Ser (UCA) and Leu (UUG, UUA, and CUC; Fig. 4b, 4d) codons. Each of these are decoded by an ac4C-containing tRNA.

**Figure 4.**
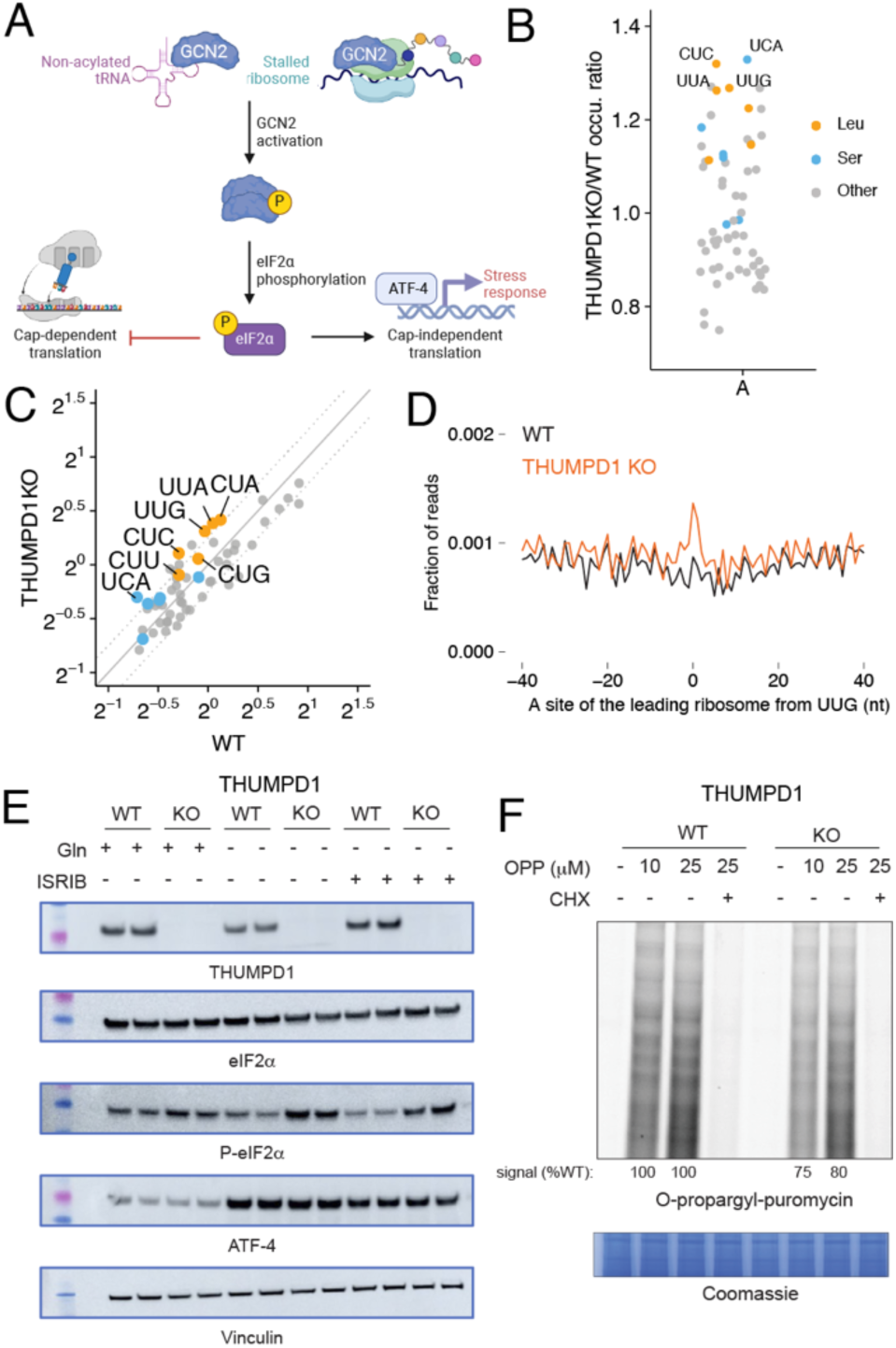
THUMPD1-mediated tRNA acetylation regulates ribosome collisions. (a) Defective tRNA function can activate the sensor kinase GCN2 and trigger the integrated stress response (ISR). (b) Disome occupancies of WT v. THUMPD1 KO cells at the ribosomal A site of the leading ribosome indicate increased disome formation at codons (UCA, CUC, UUG, and UUA) decoded by tRNA^Leu/Ser^. Codons encoding Ser, Leu, and other amino acids are shown in orange, blue, and grey, respectively. Data are representative of n=2 biological replicates. (c) Scatter plot of disome occupancies at 61 sense codons in THUMPD1 KO v. WT cells indicates increased ribosome collisions at multiple codons decoded by tRNA^Leu/Ser^. (d) Meta-codon plot centered at UUG codons shows increased ribosome collisions in THUMPD1 KO (orange) compared to WT cells (black). (e) Analysis of eIF2a phosphorylation in THUMPD1 KO cells. Amino acid deprivation (-Gln) and ISRIB (1 μM) were used to induce translational stress. Biological replicates are loaded in adjacent lanes. (f) Analysis of global translation in THUMPD1 KO cells. O-propargyl puromycin (OPP) was used to label nascent transcripts, which were then ligated to a fluorophore-azide and visualized via SDS-PAGE. Densitometry analysis was calculated using ImageJ software and is graphed in Fig. S4g.

To determine whether ribosome collisions were increase ISR kinase activation, we analyzed lysates derived from THUMPD1 KO cells with a site-specific Ser51 P-eIF2α antibody. A subtle but reproducible increase in P-eIF2α was observed in THUMPD1 KO cells (Fig. 4e), which became more apparent upon glutamine (Gln) deprivation or ISRIB treatment. Interestingly, THUMPD1 KO by itself was not sufficient to activate ATF4 (Fig. 4e). This is in line with our proteomic and transcriptomic data, which did not find upregulation of canonical ATF4 targets such as asparagine synthetase (ASNS) in THUMPD1 KO cells (Fig. S4c). A genome-wide screen for novel regulators of ATF4 activation also did not identify THUMPD1.^44^ The threshold at which Gln deprivation activates ATF4 expression was also not altered in THUMPD1 KO cells (Fig. S4d). This suggests either THUMPD1 is not a sufficiently strong stimuli to activate ATF4 or that our cell lines harbor compensatory mechanisms that blunt ISR activation in response to permanent THUMPD1 loss.

As increased levels of P-eIF2α would be expected to impede translation, we assessed nascent protein synthesis by subjecting wild-type and KO cells to labeling by O-propargyl puromycin (OPP). Loss of THUMPD1 was associated with lower levels of OPP incorporation, consistent with a translation defect (Fig. 4f, S4g). In addition to reduced translation caused by eIF2α phosphorylation, our proteomic data also observed a significant decrease in ribosomal proteins in THUMPD1 KO cells (Fig. S4d) This is in line with the notion that serial passage of THUMPD1 KO cells may select for clones that adapt to increased ribosome collisions by downregulating the translational machinery. Consistent with this view, genome-wide Perturb-seq recently found the single-cell RNA-seq signature of THUMPD1KO was most similar to those elicited by disrupting of genes involved in ribosome biosynthesis, including POLR1A, POLR1B, POLR1C, and POLR1E (Fig. S4f).^45^ Examination of THUMPD1KO cells did not reveal changes in mTOR activation, suggesting the effects of THUMPD1on ribosomal protein levels are not mediated by this pathway. (Fig. S4e). Due to the aforementioned potential for compensatory changes to be selected for in KO cell lines care must be taken not to overinterpret these findings. However, the identification of a link between THUMPD1, eIF2α phosphorylation, and translational downregulation is consistent with recent studies in model organisms.^10, 46^ These results indicate that loss of THUMPD1-dependent ac^4^C can cause ribosome collisions associated with eIF2α phosphorylation and defective translation, spurring us to explore the significance of this mechanism *in vivo*.

### Thumpd1 genetically interacts with Gcn2 in vivo

Next, we sought to understand if our observations from HEK-293T cells were relevant *in vivo*. As an initial test, we isolated tissues from age-matched wild type and Thumpd1 KO mice and assessed levels of P- eIF2α by immunohistochemical staining (Fig. 5a). While eIF2α levels were statistically identical across all samples, the percentage of P-eIF2α positive cells was significantly increased in cryosections isolated from the brain (hippocampus and cerebral cortex; Fig. 5b). This change was organ-specific, as P-eIF2α levels in the kidney and liver were not affected (Fig. 5b, Fig. S5a). Applying mim-tRNA-Seq to small RNA fractions isolated from cerebellum and liver, we again observed tRNA^Leu^ and tRNA^Ser^ to be the most downregulated (Fig. 5c-d, Supplementary Table S7). Consistent with the increased eIF2α phosphorylation observed in brain tissues, more isodecoders were impacted in cerebellum (7) than liver (2), with the caveat that the data was overall noisier in the former tissue. To further investigate the consequences of Thumpd1 KO in cerebellum we carried out RNA-Seq analysis. A recent study found exit from pluripotency is associated with eIF2α phosphorylation and used WT and eIF2α S51A mouse embryonic stem cells to define a set of transcripts dependent on eIF2α phosphorylation upon withdrawal of “2i” (two chemical inhibitors of MEK1/2 and GSK3α/β).^47^ Overlaying these genes onto our dataset, we found the majority of them were upregulated upon Thumpd1 KO, consistent with eIF2α phosphorylation (Fig. 5e cyan, Supplementary Table S8). To assess activation of the ISR, we further compared our differentially expressed gene set to Atf4 target genes previously identified in mouse embryonic fibroblasts (Fig. 5e, red).^48,12^ Here, we found a statistically significant upregulation of several ISR genes including Atf4 itself (FC_KO v. WT_ = 1.25x, adjusted *p*-value = 8.7E-4). However, the majority of the eIF2α and Atf4-dependent gene sets were unaffected, consistent with a modest response. Transcriptomic analysis of liver tissue isolated did not show this signature (Supplementary Table S9). Notably, among many affected pathways Gene Set Enrichment Analysis indicated upregulation of inflammatory signaling in cerebellum but downregulation of the mTOR signaling in liver, possibly indicating a selective compensatory response in the latter tissue (Fig. S5b, Supplementary Tables 8-9). These findings indicate a degree of concordance between our *in vitro* and *in vivo* results and prompted us to explore links between Thumpd1 and the stress response in further detail.

**Figure 5.**
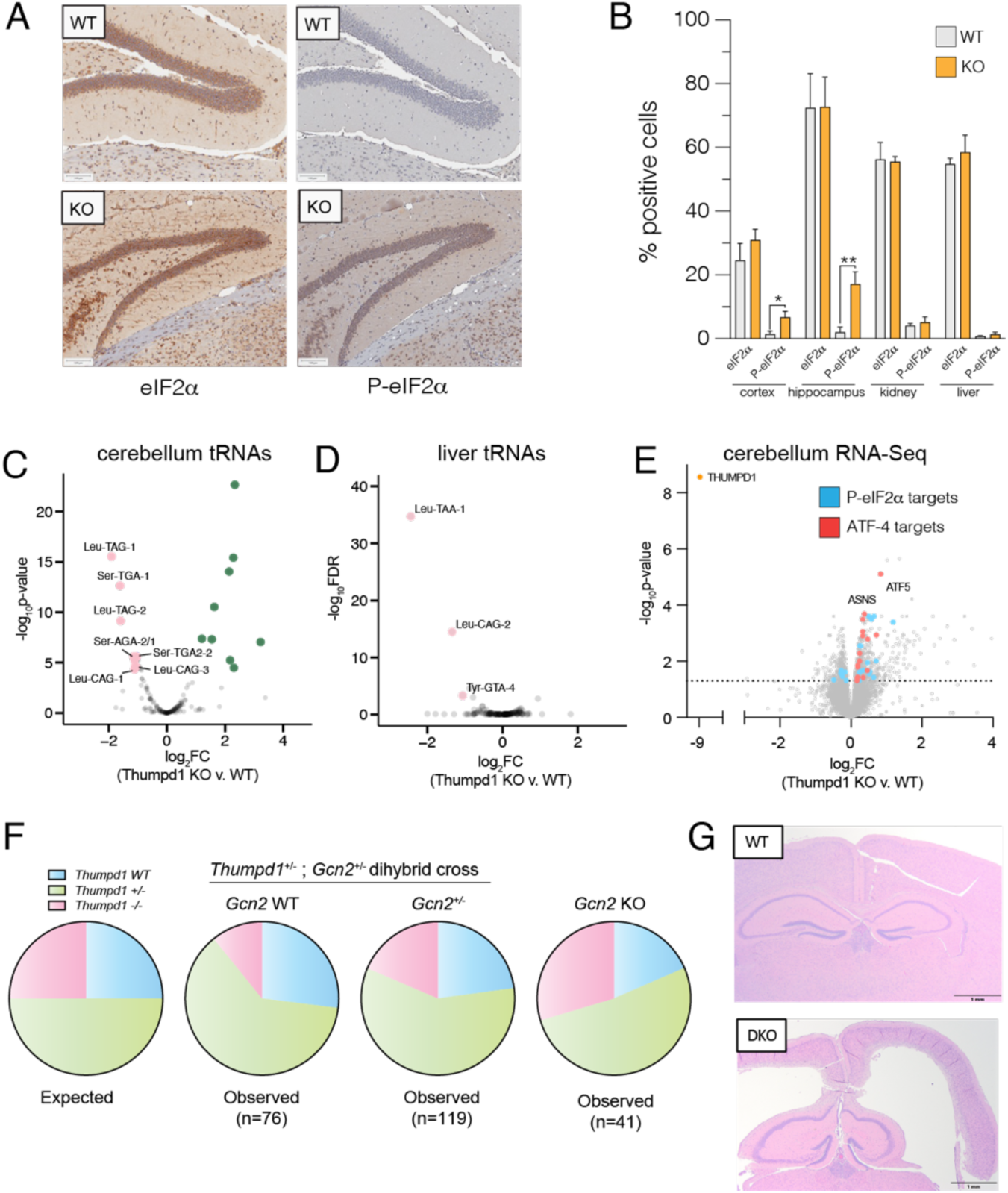
Thumpd1 and Gcn2 interact *in vivo*. (a) Immunohistochemical (IHC) staining of total eIF2a (left) and (Ser^51^) P-eIF2a (right) in brain tissue isolated from age-matched WT and Thumpd1 KO mice. Data are representative of n=4 biological replicates. (b) Quantification of percent positive cells in cerebral cortex, hippocampus, kidney cortex, and liver. (c-d) mim-tRNA-Seq analysis of tRNA levels in cerebellum (c) and liver (d) of WT and Thumpd1 KO mice show reduced levels of tRNA^Leu/Ser^ isodecoders. Data are derived from n=2 biological replicates for each tissue. (e) Gene expression profiling of cerebella isolated from WT and Thumpd1 KO mice. Literature annotated targets of Atf4 (red) and P-eIF2a (blue) exhibit a skew towards greater expression upon Thumpd1 KO. Values represent the average of n=4 biological replicates. (f) Analysis of off-spring produced by *Thumpd1^+/-^*,*Gcn2^+/-^*dihybrid cross. Pie charts are organized by *Gcn2* genotype, with WT on the left, heterozygotes in middle, and KO’s on left. The proportion of mice born born on a given background are color coded as follows: blue = *Thumpd1^+/+^*, green = *Thumpd1^-+-^*, pink = *Thumpd1^-/-^* Full numbers for dihybrid cross are provided in Fig. S6a. (g) Pathology analysis of WT and *Thumpd1^-/-^*, *Gcn^-/-^* double knockout (DKO) mouse. In the DKO animal (bottom), lateral ventricles are moderately expanded by increased clear space, resulting in loss of the neuroparenchyma in the cerebral cortex.

Gcn2 is the most well-characterized sensor kinase involved in the tRNA-dependent ISR. In budding yeast (*S. cerevisiae)* deletion of a Gcn2 homologue protects from growth defects caused by the loss of tRNA body modifications such as ac^4^C, while in fission yeast (*S. pombe)* the opposite occurs.^9, 26, 46, 49^ The Janus-like ability of Gcn2 to either propagate or mitigate tRNA-dependent phenotypes has also been observed in mammalian systems. Evidence indicates an overactive integrated stress response helps drive pathology in Charcot-Marie-Tooth (CMT) disease,^50^ a hereditary disorder that can be driven by dominant mutations in several tRNA synthetase genes. This has led to Gcn2 inhibition being explored as a therapeutic approach.^51^ Alternatively, the group of Ackerman has shown that activation of Gcn2 plays a protective role in animal models of cerebellar and retinal degeneration caused by dual mutations in the ribosome rescue factor GTPBP2 and CNS-specific tRNA^Arg^^12, 52^ However, the *in vivo* genetic interaction of Gcn2 with loss of a non-essential tRNA body modification has never been explored.

To test for a potential genetic interaction between *Thumpd1* and *Gcn2*, we generated and bred *Thumpd1*^+/-^, *Gcn2*^+/-^ males and females. *Gcn2*^-/-^ (*Thumpd1*^+/+^) mice fed a normal diet possessed no overt phenotype, consistent with previous studies.^53^ Genotyping of 262 offspring mice at day 14 (P14) after birth indicated that on the *Gcn2^WT^* and *Gcn2^+/-^* backgrounds, ∼13% of mice born were *Thumpd1^-/-^*(Fig. 5f, Fig. S6a). The sub-Mendelian production of *Thumpd1^-/-^* mice is consistent with our initial breeding studies. However, of mice with a full knockout (*Gcn2^-/-^*) background, ∼22% of mice born were *Thumpd1^-/-^*. The increased frequency of *Thumpd1^-/-^* mice born on the *Gcn2^-/-^* background is statistically significant and suggests suppression of Gcn2 activation in *Thumpd1^-/-^* KO mice beneficially impacts pre-natal fitness. In contrast, in the post-natal setting Gcn2 loss appeared to be extremely deleterious. *Gcn2*^-/−^, *Thumpd1*^-/−^ mice were extremely runted (Fig. S6b-c) and uniformly perished before P30 even when extra care measures such as delayed weaning and heat pads were provided. The early lethality of *Gcn2*^-/−^, *Thumpd1*^-/−^ dual knockout made it challenging to identify and prepare these mice for histopathological analysis prior to decomposition. However, analysis of a limited number of animals (n=4) found evidence for exacerbated phenotypes across multiple tissues, including one instance of cataract formation, increased germ cell degeneration, and a higher incidence of hydrocephalus, albeit with incomplete penetrance (Fig. 5g, Fig. S6d-e). Taken together, these results demonstrate the existence of a severe synthetic lethal genetic interaction between *Thumpd1* and *Gcn2* in mice.

## DISCUSSION

To date, the *in vivo* role of tRNA modification machinery has been most thoroughly characterized via enzymes targeting the anticodon and variable loops as well as mitochondrial tRNAs.^35, 54–66^ Here we harness *in vivo* disruption of a non-essential tRNA D-arm modification that specifically marks two cytosolic type II tRNAs to reveal a modification-dependent sensing mechanism that extends from yeast to vertebrates. Our research showcases how interrogating non-essential enzymes *in vivo* can uncover new modification-driven phenotypes and proposes a model whereby Thumpd1-dependent tRNA acetylation influences cell fate based on the severity of translational stress its loss elicits (Fig. 6).

**Figure 6.**
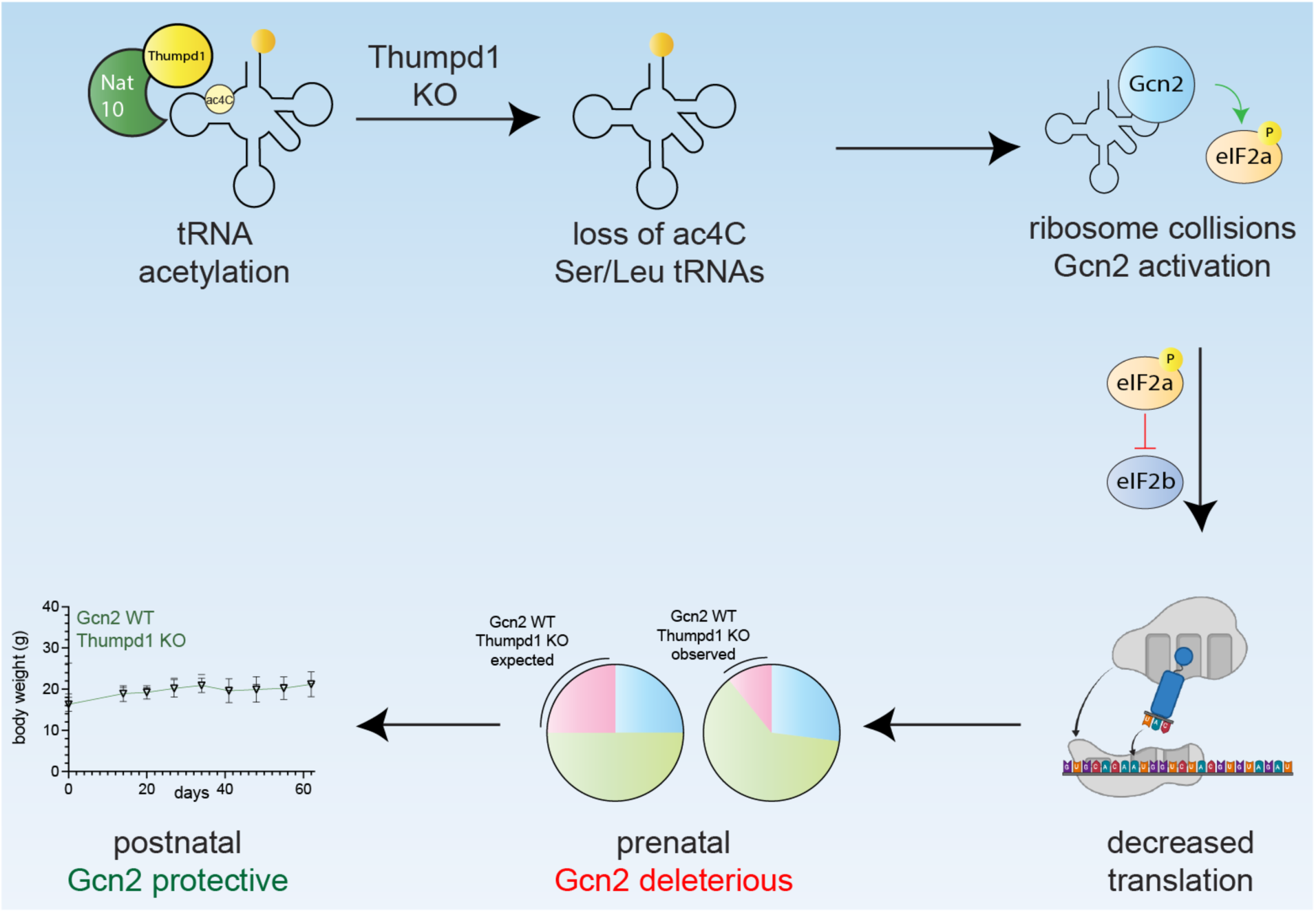
Model for Thumpd1-mediated tRNA ac^4^C in mammalian ribosome function. Nat10/Thumpd1 is responsible for C12 modification of tRNA^Leu^ and tRNA^Ser^. Genetic mutations and potentially other stimuli can cause a loss of ac^4^C from Ser/Leu tRNAs. This reduces the levels of tRNA^Leu/Ser^ isodecoders and possibly slows translocation, resulting in increased A-site occupancy at Leu/Ser codons and ribosome collisions. This activate the sensor kinase Gcn2 which causes tissue- specific phosphorylation of eIF2α. Cellular studies suggest differential activation of Gcn2 may reflect the ability of tissues to access compensatory mechanisms upon ribosome collisions for example downregulating ribosomal proteins. The sub-Mendelian production of *Thumpd1^-/-^*KO offspring is rescued by concurrent KO of *Gcn2*, suggesting Thumpd1-dependent Gcn2 activation is deleterious during prenatal development. However, dual KO animals perish shortly after birth, suggesting in the postnatal setting Gcn2 plays a protective role.

Our data suggest tRNA acetylation can impact physiology by three distinct scenarios: homeostasis, adaptive response, and cellular dysfunction. In the homeostasis scenario, normal levels of tRNA acetylation maintain optimal tRNA levels and translational efficiency. The adaptive response regime is triggered when THUMPD1 loss leads to reduced tRNA acetylation, causing a decrease in tRNA^Leu^ and tRNA^Ser^ levels and increased ribosomal occupancy at their respective codons. Ribosome collisions activate the Gcn2-eIF2α axis, promoting a modest stress response that allows cells to adapt to the translational perturbation. In the cellular dysfunction regime, the lack of Gcn2 - or tissue-specific compensatory factors which remain to be defined - enables translational stress to surpass a critical threshold, leading to apoptosis and organismal lethality.

Breeding studies revealed a complex interplay between Thumpd1 and Gcn2. Gcn2 loss appeared to improve prenatal fitness in Thumpd1-deficient mice, suggesting excessive Gcn2 activation is detrimental to embryonic development. However, Thumpd1/Gcn2 DKO mice exhibited severe postnatal lethality, indicating a crucial protective role for Gcn2 in the postnatal setting. This finding also hints at other contexts where Thumpd1-mediated tRNA acetylation may be required. For instance, the parallels between embryonic development and cancer metastasis led us to carry out a pilot study in which heterozygous *Thumpd1^+/-^* mice - which harbor quantitatively lower levels of tRNA ac^4^C – were crossed into a model of breast cancer metastasis (Fig. S7a-b). Primary tumor burden was not affected in *Thumpd1^+/-^* mice; however, metastases were significantly repressed (Fig. S7). While we defer further characterization to future work, this experiment provides another example of the biological relevance of tRNA ac^4^C and the enabling nature of the models developed here.

Our work raises several questions for future investigation. First, the mechanism by which loss of ac^4^C triggered tRNA degradation remains undefined. While the rapid decay of hypomodified tRNAs is well-known in yeast,^9, 67^ in mammals these pathways are less extensively characterized. SLFN11 and SLFN12 are the most well-known regulators of type II tRNAs in mammals^68, 69^ but are not present in HEK-293T cells^70, 71^ raising the possibility of novel degradation mechanisms. Furthermore, we did not determine the precise cause of ribosome stalling in the absence of tRNA acetylation. In theory, A-site stalling could reflect rate-limited tRNA^Leu/Ser^ levels, altered activity of these tRNAs during translation, or some combination thereof. Previous studies have noted the proximity of the D-stem to helix 69 of rRNA during translocation,^43, 72^ suggesting the potential for ac^4^C -dependent conformational dynamics to directly influence this process. In addition, the adaptations and tissue-specific responses to tRNA acetylation raise the question: are there sensing mechanisms that directly link ac^4^C or tRNA^Leu/Ser^ to translational control? Our results provide a starting point for pursuing these questions, whose answers could afford critical insights into the role of tRNA modifications in development and disease.

Finally, our studies suggest new avenues that may be explored for treatment of THUMPD1-dependent neurodegenerative effects. Given GCN2 KO potentiates the deleterious effects of THUMPD1 loss, we speculate that THUMPD1-deficient patients may benefit from strategies that increase eIF2α phosphorylation and reduce ribosome collisions. Genetic ablation of CHOP^73^ and small molecule inhibition of the eIF2α phosphatase GADD34^74^ have shown promise in models of tRNA-related neurological disorders.^75^ In contrast, molecules such as ISRIB that activate eIF2α phosphorylation by amplifying ribosome collisions may be less desirable. A recent report described the ability of the serine/threonine kinase ZAKα (ZAK) to sense ribosome collisions and establish regimes of either tolerance or cell death in response to translational dysfunction.^76^ While the role of this apoptotic mechanism in THUMPD1-dependent phenotypes is yet undetermined, clinically approved kinase inhibitors including nilotinib (BCR-ABL) and vemurafenib (BRAF) show crossover inhibition of ZAK, providing a potential approach to probe this mechanism. Another strategy to combat ribosome collisions would be to downregulate the translational machinery, for instance by mTOR inhibition. This strategy is supported by studies in yeast showing that ribosomal protein mutations can suppress growth defects caused by tRNA acetylation loss^10^ and by our observations of compensatory ribosomal protein downregulation in THUMPD1-deficient human cells. The *in vivo* models and phenotypes described in this study provide a robust foundation for testing these hypotheses. Collectively, our studies define the ability of tRNA acetylation to impact translational signaling in mammalian lineages, providing a conceptual foundation for understanding the role of this modification in biology and disease.

### Limitations of the study

Our work characterizes the occurrence of ac^4^C at C12 across a variety of tRNA^Ser^ and tRNA^Leu^ isodecoders. However, we cannot rule out that additional sites of tRNA acetylation may occur in different cell types or tissues. Similarly, our *in vitro* mechanistic studies were confined to a human cell line and do dismiss the possibility that more profound codon-biased translation may be regulated by THUMPD1-dependent ac^4^C in other settings. Delineating these and other potential mechanistic inputs such as tRNA fragments^77^ will be important for a holistic understanding of how tRNA acetylation influences translation.

Since the only characterized function of THUMPD1is to assist deposition of ac^4^C in eukaryotic tRNA, a parsimonious model is that the phenotypes observed in our study reflect changes in ac^4^C-dependent tRNA function. This notion is supported by the fact that ectopic expression of tRNA^Ser^ can rescue *S. cerevisiae* growth defects caused by mutation of a Thumpd1 homologue.^9, 26^ However, we cannot rule out the possibility that THUMPD1-dependent phenotypes may stem from an uncharacterized chaperone role of this protein.^78^ Structural and biochemical analyses of the eukaryotic NAT10/ THUMPD1complex will be important to constructing mutants that can more precisely modulate THUMPD1’sactivity.

Our strategy for modulating ac^4^C contrasts with studies using NAT10 deletion,^79^ a more severe perturbation that is not known to cause eIF2α phosphorylation or activation of the ISR. We hypothesize that differences in these models may arise due to the centrality of NAT10 to additional processes besides tRNA modification such as ribosome biogenesis.^6^ The development of strategies for temporal modulation of NAT10 and THUMPD1activity such as small molecule inhibition and degron-tagged models will likely be useful in differentiating first-order effects from compensatory mechanisms.

Finally, while our studies provide an overview of the organismal consequences of THUMPD1 deletion but do not specify i*n vivo* mechanism or report behavioral phenotypes. Defining these attributes will undoubtedly be helpful in applying these models to study the role of THUMPD1/tRNA acetylation in neurodevelopment.

## ACKNOWLEDGEMENTS

We thank D. Tripu, D. Gallimore, and R. Hitzman (Chemical Biology Laboratory, National Cancer Institute) for assistance optimizing cell growth and tissue extraction, D. McVicar, E. Palmieri, and J. Weiss (Cancer Innovation Laboratory, National Cancer Institute) and N. Snyder (Temple University) for preliminary characterization of Thumpd1 KO cells, P. Awasthi (Laboratory Animal Sciences Program, Frederick National Laboratory for Cancer Research) for mouse model assistance and J. Labrador and E. Kosmeder (Laboratory of Cancer Biology and Genetics, National Cancer Institute) and D. Butcher (Laboratory Animal Sciences Program, Frederick National Laboratory for Cancer Research) for technical assistance. J.L.M. is supported by the Intramural Research Program of the National Institutes of Health (NIH), the National Cancer Institute, The Center for Cancer Research (ZIA BC011488). C.C.-C.W. is supported by the Intramural Research Program of the National Institutes of Health (NIH), the National Cancer Institute, The Center for Cancer Research (ZIA BC012037). M.R.O. is supported by the National Institute of General Medical Sciences of the NIH (R35 GM133462). S. S. is funded by the Israel Science Foundation (grant no. 913/21), the European Research Council (ERC) under the European Union’s Horizon 2020 research and innovation programme (grant no. 101000970). In addition, this project has been funded in part with federal funds from the National Cancer Institute, National Institutes of Health, under contract number HHSN261200800001E and HHSN261201500003I.

## AUTHOR CONTRIBUTIONS

Conceptualization, M.R.O., C.C.C.W., and J.L.M.; Methodology, S.T.G., R.K., S.H.M., M.C.C., J.G., B.V.P., C.S., K.A.L., K.D.N., V.I., A.K., M.P., C.T.T., S.R.N., C.N.M., R.J.H., T.M., V.K., A.Y., P.J., J.T.N., T.A., B.A.M., S.G., E.F.E., S.D., R.C., S.S., M.R.O., C.C.C.W., and J.L.M.; Investigation, S.T.G., R.K., S.H.M., M.C.C., J.G., B.V.P., C.S., K.A.L., K.D.N., V.I., A.K., M.P., C.T.T., S.R.N., R.J.H., T.M., P.J., J.T.N., T.A., B.A.M., E.F.E., S.D., R.C., S.S., M.R.O., C.C.C.W., and J.L.M.; Resources, V.K. and R.C. Visualization, S.T.G., R.K., S.H.M., M.C.C., J.G., C.C.C.W., and J.L.M.; Data Curation, S.H.M., J.G., R.J.H., T.M., V.K., and R.C.; Writing, S.T.G., R.K., S.S., M.R.O., C.C.C.W., and J.L.M.; Supervision, C.S., P.J., T.A., B.A.M., S.G., S.D., R.C., S.S., M.R.O., C.C.C.W., and J.L.M.;

## DECLARATION OF INTERESTS

The authors have no positions or financial interests to declare.

## SUPPLEMENTARY INFORMATION

Full experimental details, additional figures, tables, experimental protocols, full gels, and nucleotide sequences are available as supplementary information.

## SUPPLEMENTARY FIGURES

**Figure S1.**
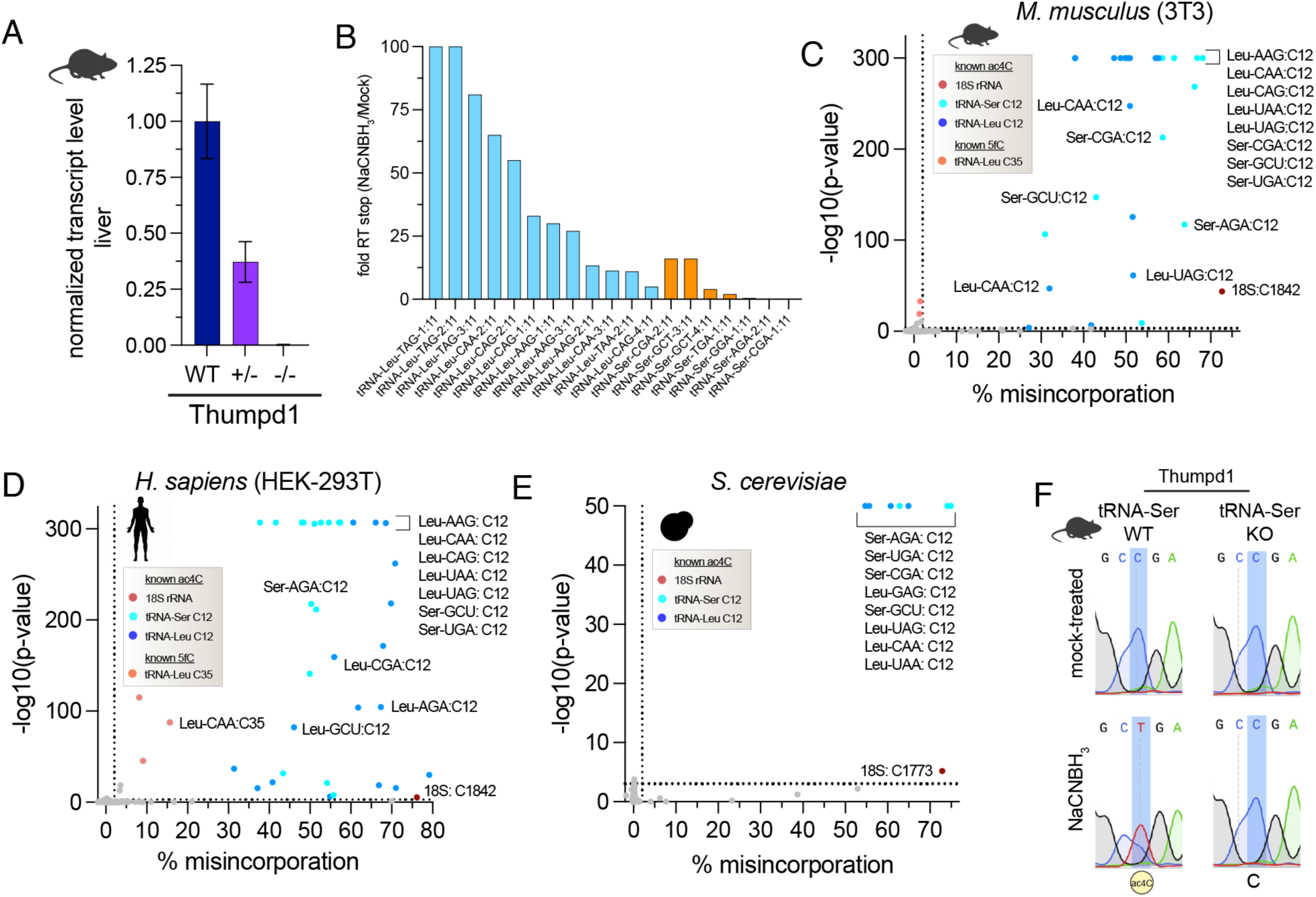
(a) Quantitative real-time PCR-based analysis of *Thumpd1* gene expression in RNA isolated from liver of *Thumpd1^WT^*, *Thumpd^+/-^*, or *Thumpd1^-/-^* KO lines. Data represent the average of n=3 biological replicates. (b) Analysis of RT stops in tRNA^Leu^ (blue) and tRNA^Ser^ (orange). Fold RT stop was estimated by comparing the number of reads starting at C11 in NaCNBH_3_-treated (‘starts.sample’) versus control (‘starts.control’). At positions where no stops were observed in the control the ratio was arbitrarily set to 100. (c) Distribution of ac^4^C in murine (3T3) small RNA fraction. (d) Distribution of ac^4^C in *H. sapiens* (HEK-293T) small RNA fraction. (e) Distribution of ac^4^C in *S. cerevisiae* small RNA fraction. Values for c-e were calculated from the ‘C2T.MRD’ and ‘pval.CT2’ columns which correspond to the C -> T misincorporation rate and C->T p-value, respectively in Table S1. Nucleotide with a ‘pval.CT2’ of 0 were graphed on the y-axis at the value corresponding to the lowest calculatable p-value. (f) Sanger sequencing based ac^4^C sequencing confirms loss of ac^4^C in murine tRNA upon Thumpd1 knockout.

**Figure S2.**
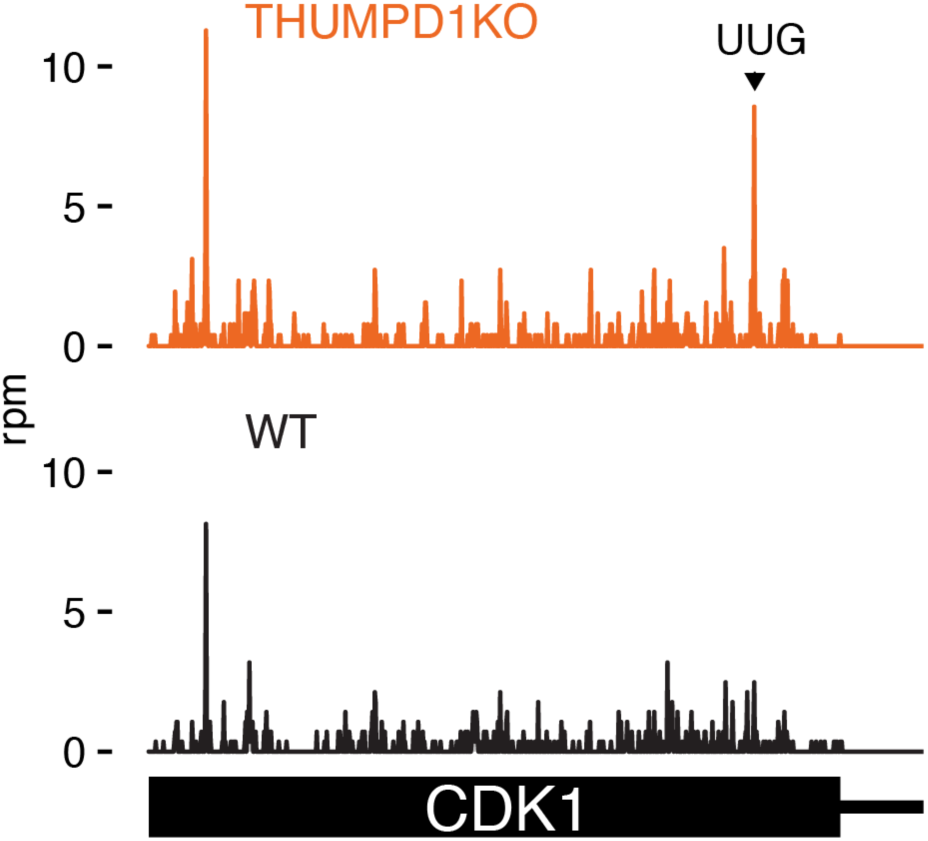
Analysis of codon occupancy across the *CDK1* transcript, with a stalling site at a Leu UUG codon highlighted.

**Figure S3.**
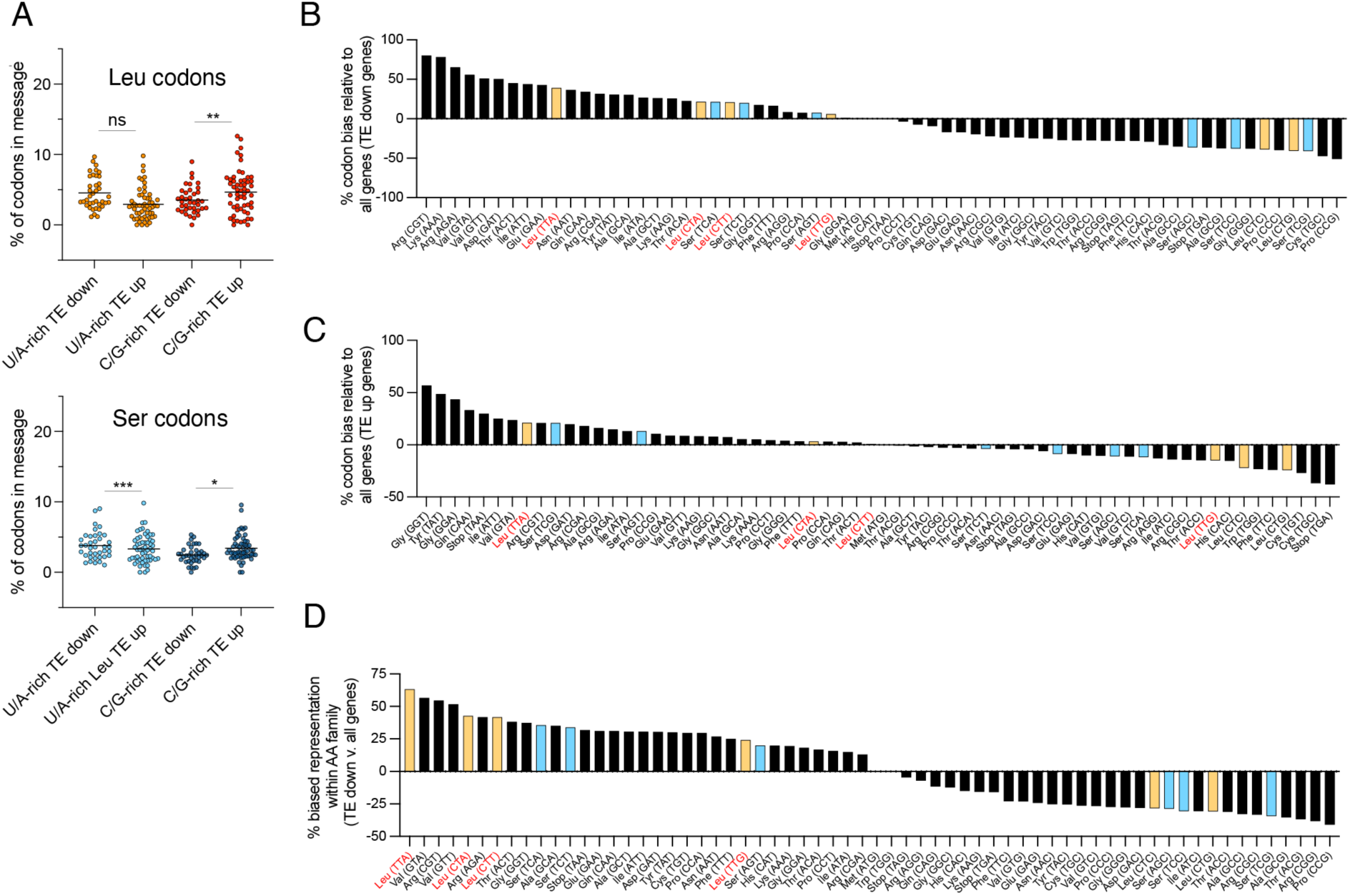
(a) Top: Comparative analysis of U/A-rich Leu codons (UUA, CUA, CUU, UUG) and C/G-rich Leu codons (CUG, CUC) in genes with increased (TE up) or decreased (TE down) translational efficiency in THUMPD1 KO HEK-293T cells. Bottom: Comparative analysis of U/A-rich Ser codons (UCA, UCU, AGU) and C/G-rich Ser codons (AGC, UCC, UCG) in genes with increased (TE up) or decreased (TE down) translational efficiency in THUMPD1 KO HEK-293T cells. Each point represents the combined codon content of a single TE up or TE down transcript. Significance was analyzed by two-tailed Student’s *t* test (ns = not significant, * = *P* < 0.05, ** = *P* < 0.01, and *** = *P* < 0.001). (b) Analysis of codon bias in TE down transcripts relative to all CCDS-defined consensus coding sequences. A positive value indicates an increased proportion of the codon is present in TE down sequences. Leu codons = orange, Ser codons = blue, U/A-rich Leu codons are labeled on the x-axis in red. (c) Analysis of codon bias in TE up transcripts relative to all CCDS-defined consensus coding sequences. A positive value indicates an increased proportion of the codon is present in TE up sequences. Leu codons = orange, Ser codons = blue, U/A-rich Leu codons are labeled on the x-axis in red. (d) Analysis of amino acid family-specific codon bias in TE down transcripts relative to all CCDS-defined consensus coding sequences. For this analysis, the representation of each individual codon (e.g. Leu-UUA) relative to its amino acid family (e.g. all Leu) was calculated, and then the average values for TE down sequences were compared to the average values for all CCDS-defined consensus coding sequences. Leu codons = orange, Ser codons = blue, U/A-rich Leu codons are labeled on the x-axis in red.

**Figure S4.**
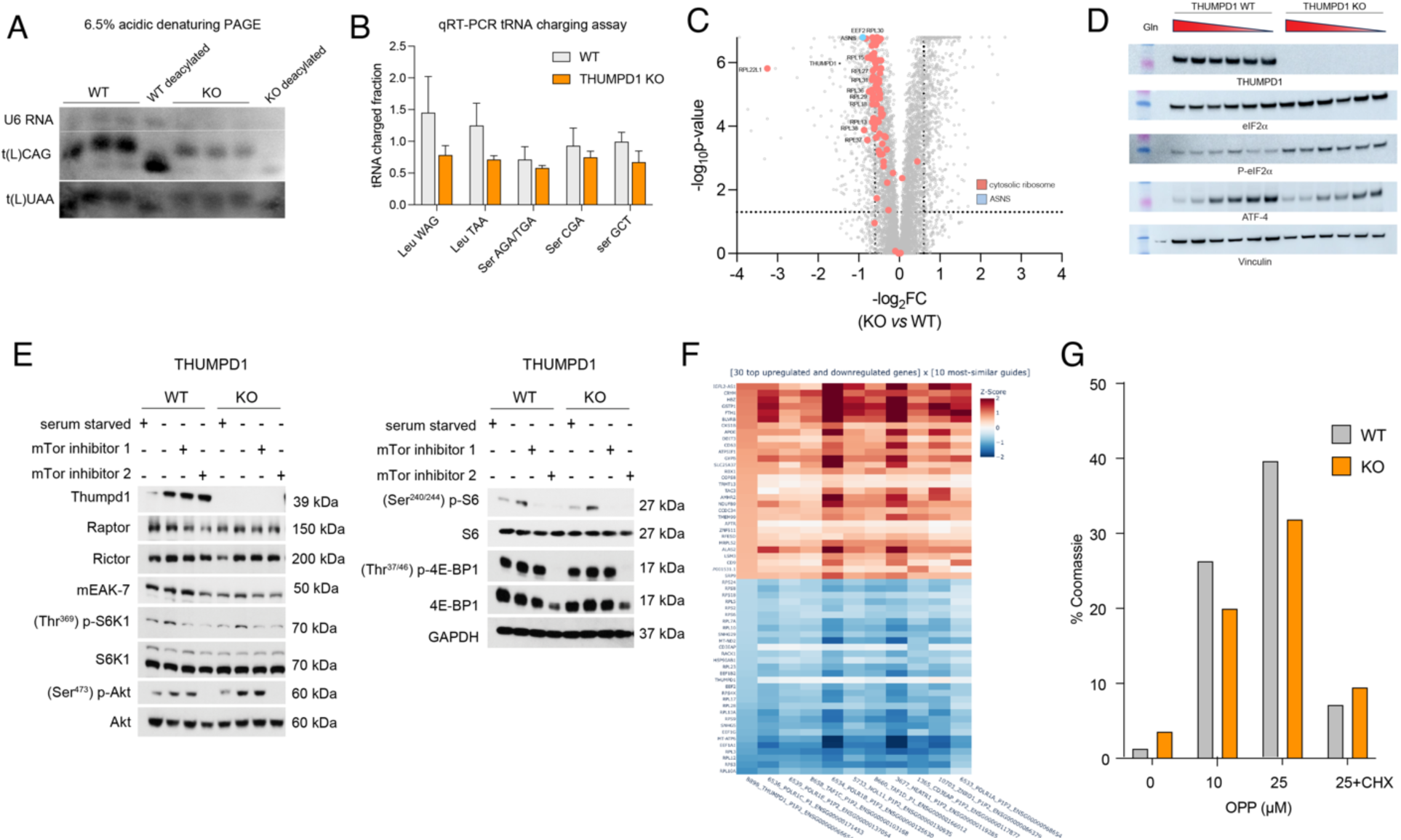
(a) Acidic denaturing PAGE analysis of tRNA charging in WT or THUMPD1 KO HEK-293T cells. Data are representative of n=2 biological replicates. (b) qRT-PCR analysis of tRNA charging in WT or THUMPD1 KO HEK-293T cells. Values for WT and KO tRNA paires were not significant as analyzed by two-tailed Student’s *t* test (*P* > 0.05). Data are representative of n=2 biological replicates. (c) Ribosomal proteins (red) are downregulated in THUMPD1 KO HEK-293T cells. The ATF-4 target ASNS (blue) is also downregulated, suggesting the ISR is not activated by THUMPD1 KO. Values are derived from n=3 biological replicates. (d) THUMPD1 KO does not alter the threshold for glutamine-dependent activation of ATF-4. Glutamine concentrations (left to right): 2 mM, 0.2 mM, 0.02 mM, 0.002 mM, and no glutamine. (e) THUMPD1 KO cells do not show altered ability to activate mTOR signaling. Treatment conditions as follows: serum starved = removal of all amino acids and serum from medium (1 h), mTor inhibitor 1 = 1 µM AZD2014 (1 h), mTor inhibitor 2 = 10 nM rapamycin (1 h). Data are representative of n=2 biological replicates. (f) Single-cell RNA-Seq signature of THUMPD1 KO cells observed by Perturb-seq analysis of Replogle *et al*.^31^ (g) Gel densitometry analysis of fluorescence signal from treatment of THUMPD1 WT and KO HEK-293T with *O*-propargyl puromycin (OPP) followed by click chemistry to a fluorescent azide. The percent of the fluorescent signal relative to the Coomassie signal was calculated, and used to produce the relative values given in Fig. 4g. Data are representative of n=2 biological replicates.

**Figure S5.**
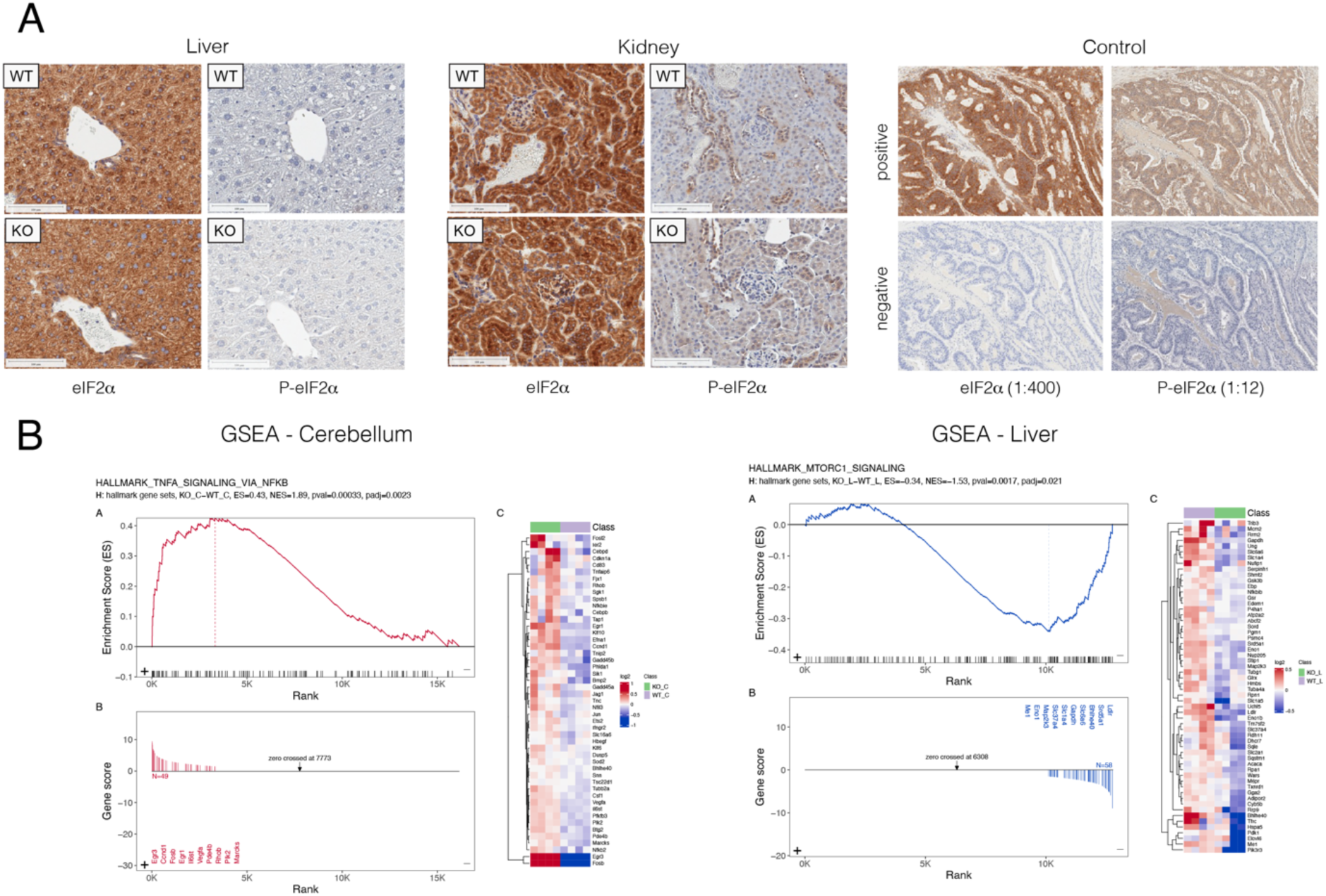
(a) Immunohistochemical (IHC) staining of total eIF2α (left) and (Ser^50^) P-eIF2α (right) in liver tissue and kidney tissue isolated from age-matched WT and *Thumpd1^-/-^* KO mice. Results are representative of n=4 biological replicates. (b) Gene Set Enrichment Analysis indicating activation of inflammatory gene expression in mouse cerebellum (TNFA_SIGNALING_VIA_NFKB, left) and downregulation of transcripts associated with mTOR signaling (MTORC1_SIGNALING, right) in mouse liver. Pathway analyses were generated from RNA-Seq data (n=4 biological replicates). Additional pathway analyses are provided in Supplementary Tables 8-9.

**Figure S6.**
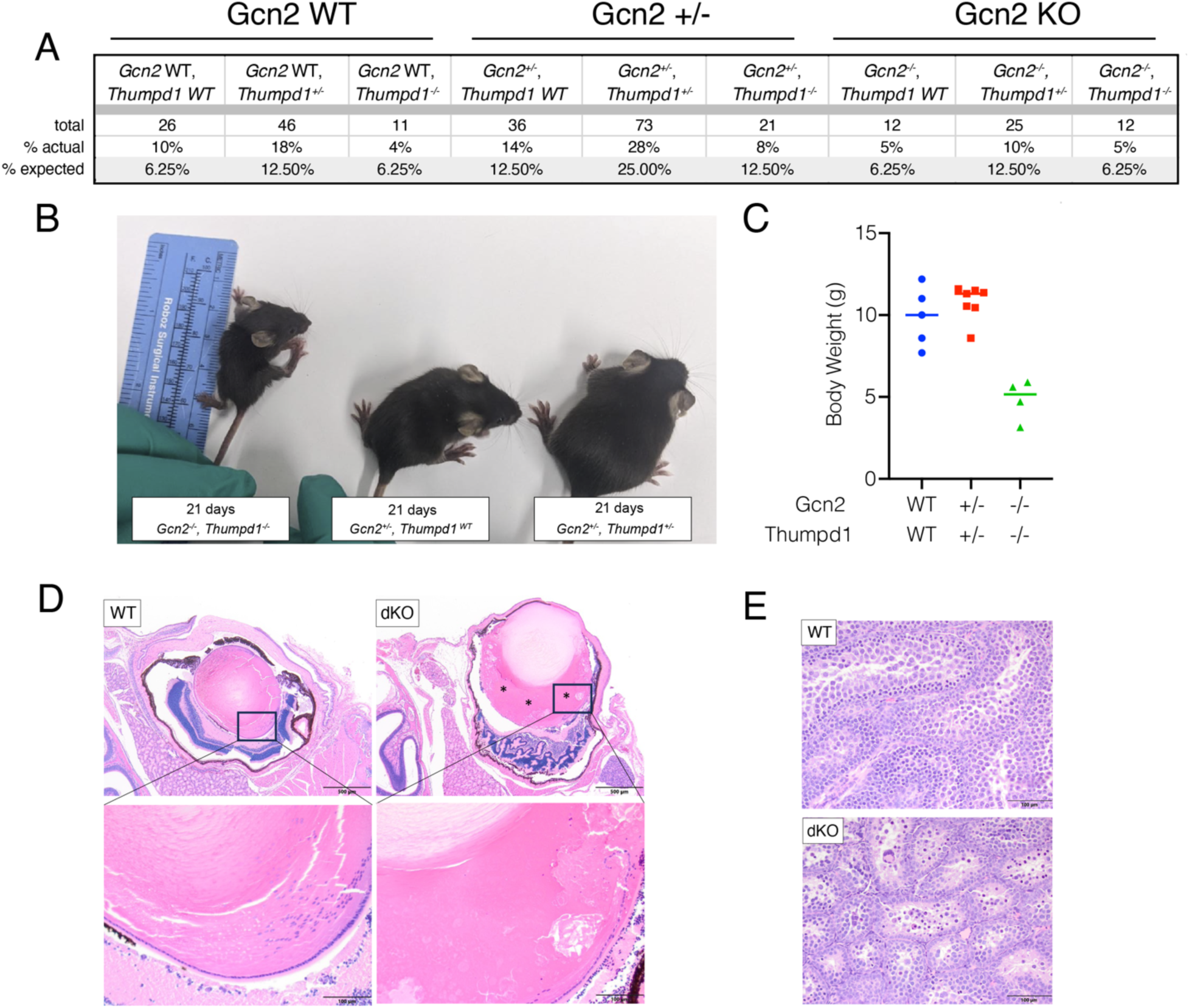
(a) Offspring annotated by genotype produced by *Thumpd1^+/-^*,*Gcn2^+/-^* dihybrid cross. (b-c) *Thumpd1*/*Gcn2* double KO mice are runted. (d) A *Thumpd1^-/-^/Gcn2^-/-^* double knockout (DKO) animal exhibits cataractous change where the lens is expanded by a liquefaction of lens fibers (*), which lack organization and are swollen and fragmented, often forming globules of degenerate lens proteins (Morgagnian globules). Hyperplasia of the lens epithelium is also observed. The animal also exhibits retinal dysplasia, where the retina is disorganized, poorly developed, and thrown into folds. There is retinal detachment with hypertrophy (tomb-stoning) of the retinal pigmented epithelium. (e) *Thumpd1^-/-^*, *Gcn2^-/-^* DKO mice exhibit multifocal seminiferous tubule degeneration with syncytial cell formation. Due to the challenge of isolating *Thumpd1^-/-^*, *Gcn2^-/-^* DKO mice prior to lethality and autoloysis onset, data for d-e are representative of n=1individual mouse.

**Figure S7.**
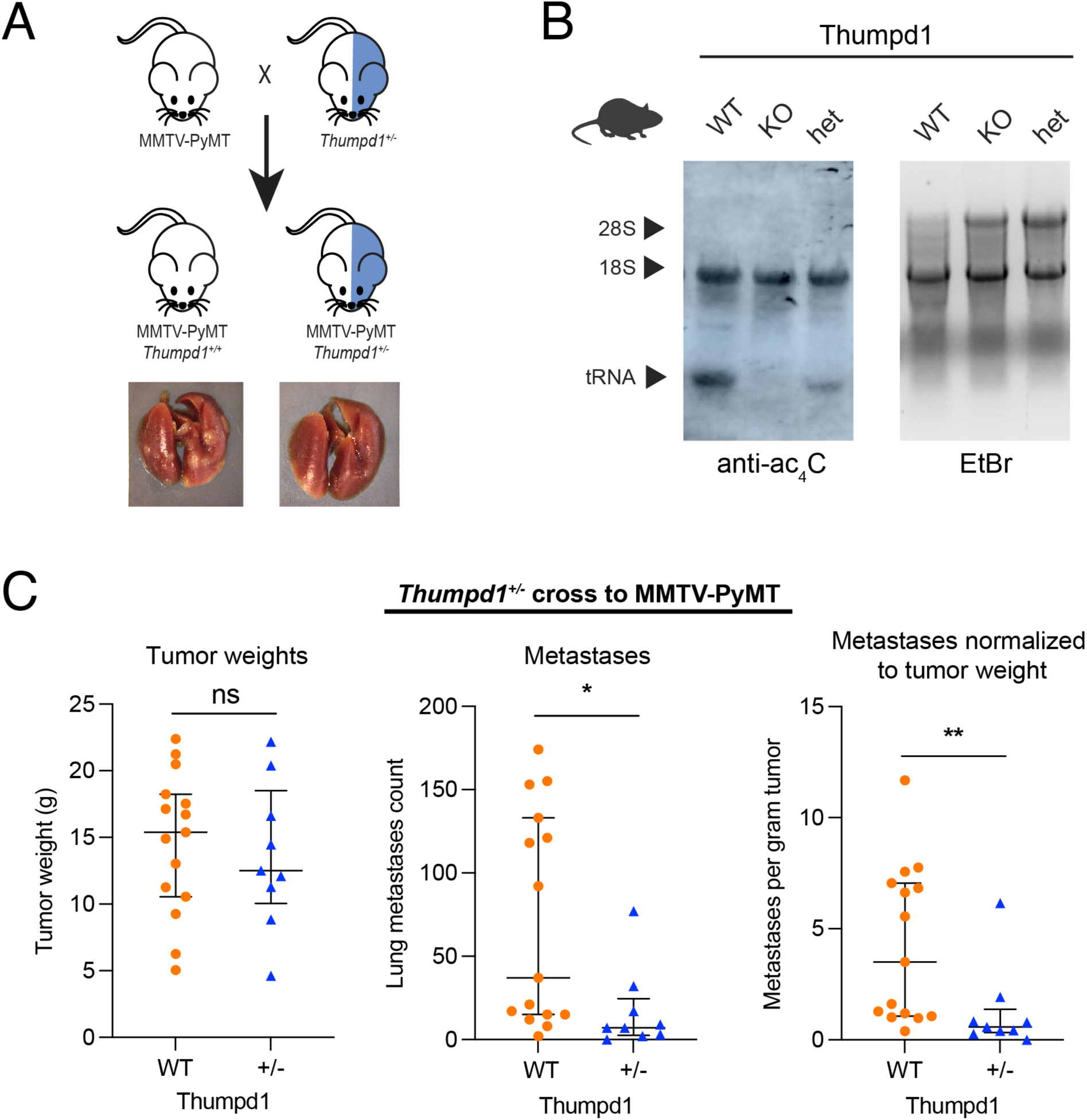
(a) Schematic for PyMT tumor metastasis model. (b) *Thumpd1^+/-^* heterozygotes exhibit qualitatively decreased tRNA acetylation as assessed by immuno-Northern blot. (c) *Thumpd1^+/-^* heterozygosity suppresses breast cancer metastasis but not primary tumor growth in PyMT model.

## SUPPLEMENTARY TABLES

**Table S1.** ac4C-seq analysis of tRNA acetylation sites across eukaryotic models

**Table S2**. mim-tRNA-Seq quantification of tRNA levels in THUMPD1 WT and KO HEK-293T cells

**Table S3.** Ribosome profiling analysis of codon occupancy in THUMPD1 WT and KO HEK-293T cells

**Table S4.** Proteomic analysis of THUMPD1 WT and KO HEK-293T cells

**Table S5.** Codon usage analysis of transcripts with altered translational efficiency in THUMPD1 KO cells

**Table S6.** Disome profiling analysis of ribosome collisions in THUMPD1 WT and KO HEK-293T cells

**Table S7.** mim-tRNA-Seq quantification of tRNA levels in cerebellum and liver of WT and Thumpd1 KO mice

**Table S8.** Differential gene expression analysis of cerebella from WT and Thumpd1 KO mice

**Table S9.** Differential gene expression analysis of liver from WT and Thumpd1 KO mice

## Notes

### Competing Interest Statement

The authors have declared no competing interest.

### Summary of Updates

This version of the manuscript has been revised to update the author list.

