## Supplementary Information for "Transfer RNA acetylation regulates in vivo mammalian stress signaling"

1. Chemical Biology Laboratory, Center for Cancer Research, National Cancer Institute, National Institutes of Health, Frederick, MD, USA. 2. RNA Biology Laboratory, Center for Cancer Research, National Cancer Institute, National Institutes of Health, Frederick, MD, USA. 3. Department of Molecular Genetics, Weizmann Institute of Science Rehovot 76100, Israel. 4. Department of Biochemistry and Biophysics, Center for RNA Biology, School of Medicine and Dentistry, University of Rochester, Rochester, NY, USA. 5. Animal Research Technical Support, Laboratory Animal Sciences Program, Frederick National Laboratory for Cancer Research, Frederick, MD, USA 6. Genome Modification Core, Laboratory Animal Sciences Program, Frederick National Laboratory for Cancer Research (FNLCR), Frederick, MD, USA. 7. Protein Mass Spectrometry Group, Center for Cancer Research, National Cancer Institute, National Institutes of Health, Frederick, MD, USA. 8. CCR Collaborative Bioinformatics Resource (CCBR), Frederick National Laboratory for Cancer Research, Leidos Biomedical Research, Inc, Frederick, MD, USA. 9. Laboratory of Cancer Biology and Genetics, National Cancer Institute, National Institutes of Health, Bethesda, Maryland, USA. 10. Molecular Histopathology Laboratory, Laboratory Animal Sciences Program, Frederick National Laboratory for Cancer Research, Frederick, MD, USA.

**Table of Contents for Supplementary Information**

| | Page |
|---|---|
| Supplementary figures | S2 |
| Key resources table | S9 |
| Experimental protocols | S11 |
| Full gel images | S35 |
| Oligonucleotide sequences table | S41 |
| References | S44 |

### Supplementary Figures

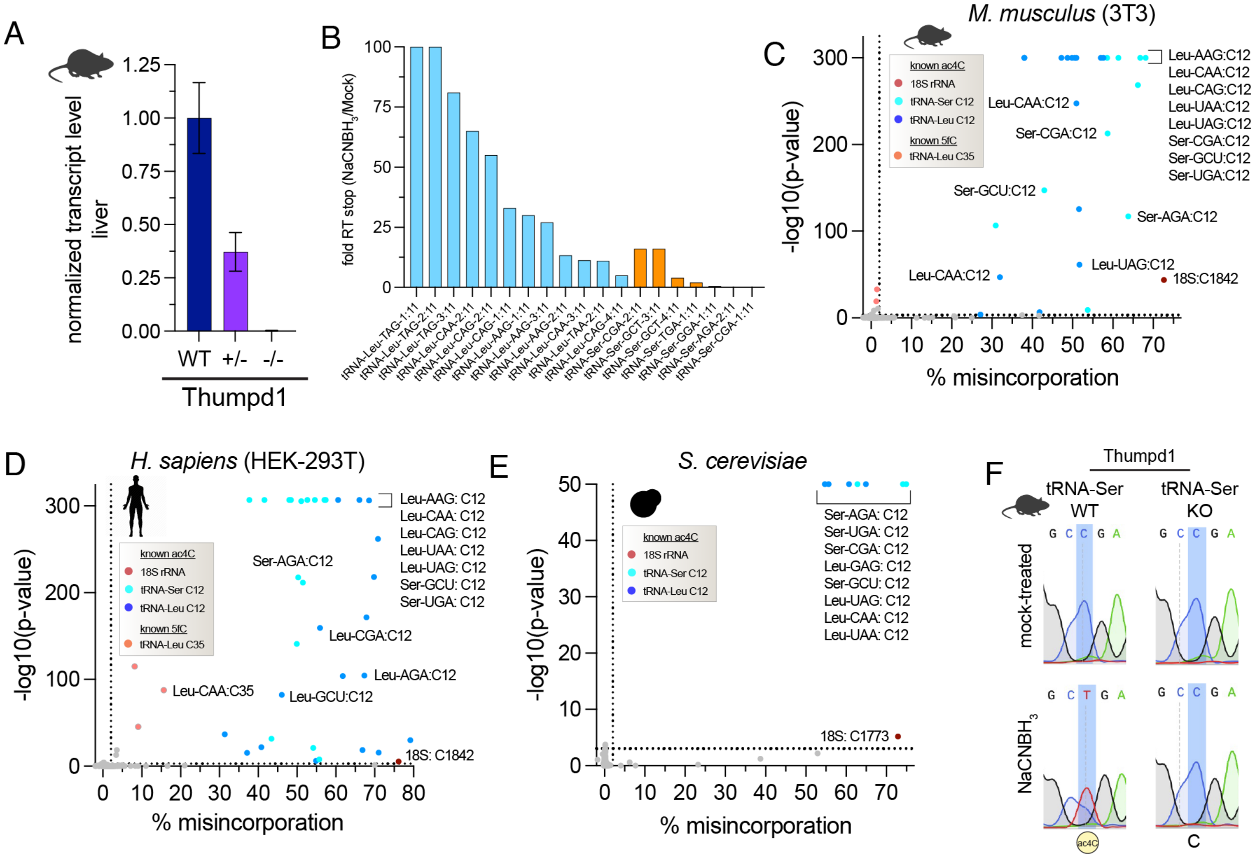


**Figure S1**. (a) Quantitative real-time PCR-based analysis of *Thumpd1* gene expression in RNA isolated from liver of *Thumpd1$^{WT}$*, *Thumpd$^{+/-}$*, or *Thumpd1$^{-/-}$* KO lines. Data represent the average of n=3 biological replicates. (b) Analysis of RT stops in tRNA$^{Leu}$ (blue) and tRNA$^{Ser}$ (orange). Fold RT stop was estimated by comparing the number of reads starting at C11 in $NaCNBH_3$-treated (‘starts.sample’) versus control (‘starts.control’). At positions where no stops were observed in the control the ratio was arbitrarily set to 100. (c) Distribution of ac$^4$C in murine (3T3) small RNA fraction. (d) Distribution of ac$^4$C in *H. sapiens* (HEK-293T) small RNA fraction. (e) Distribution of ac$^4$C in *S. cerevisiae* small RNA fraction. Values for c-e were calculated from the ‘C2T.MRD’ and ‘pval.CT2’ columns which correspond to the C -> T misincorporation rate and C->T p-value, respectively in Table S1. Nucleotide with a ‘pval.CT2’ of 0 were graphed on the y-axis at the value corresponding to the lowest calculatable p-value. (f) Sanger sequencing based ac$^4$C sequencing confirms loss of ac$^4$C in murine tRNA upon Thumpd1 knockout.

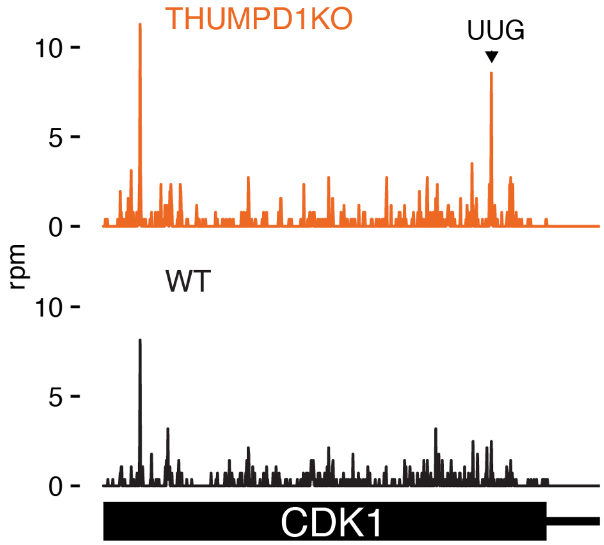


**Figure S2.** Analysis of codon occupancy across the *CDK1* transcript, with a stalling site at a Leu UUG codon highlighted.

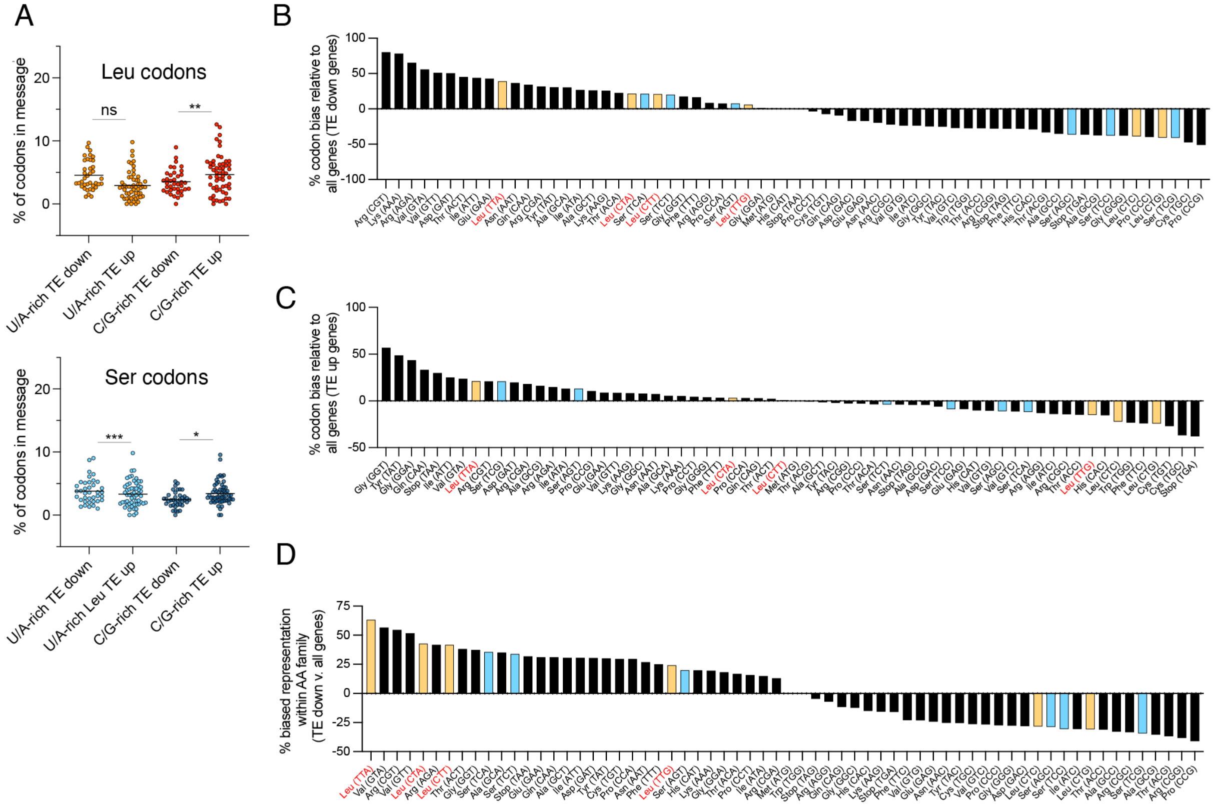


**Figure S3.** (a) Top: Comparative analysis of U/A-rich Leu codons (UUA, CUA, CUU, UUG) and C/G-rich Leu codons (CUG, CUC) in genes with increased (TE up) or decreased (TE down) translational efficiency in THUMPD1 KO HEK-293T cells. Bottom: Comparative analysis of U/A-rich Ser codons (UCA, UCU, AGU) and C/G-rich Ser codons (AGC, UCC, UCG) in genes with increased (TE up) or decreased (TE down) translational efficiency in THUMPD1 KO HEK-293T cells. Each point represents the combined codon content of a single TE up or TE down transcript. Significance was analyzed by two-tailed Student's *t* test (ns = not significant, * = $P < 0.05$, ** = $P < 0.01$, and *** = $P < 0.001$). (b) Analysis of codon bias in TE down transcripts relative to all CCDS-defined consensus coding sequences. A positive value indicates an increased proportion of the codon is present in TE down sequences. Leu codons = orange, Ser codons = blue, U/A-rich Leu codons are labeled on the x-axis in red. (c) Analysis of codon bias in TE up transcripts relative to all CCDS-defined consensus coding sequences. A positive value indicates an increased proportion of the codon is present in TE up sequences. Leu codons = orange, Ser codons = blue, U/A-rich Leu codons are labeled on the x-axis in red. (d) Analysis of amino acid family-specific codon bias in TE down transcripts relative to all CCDS-defined consensus coding sequences. For this analysis, the representation of each individual codon (e.g. Leu-UUA) relative to its amino acid family (e.g. all Leu) was calculated, and then the average values for TE down sequences were compared to the average values for all CCDS-defined consensus coding sequences. Leu codons = orange, Ser codons = blue, U/A-rich Leu codons are labeled on the x-axis in red.

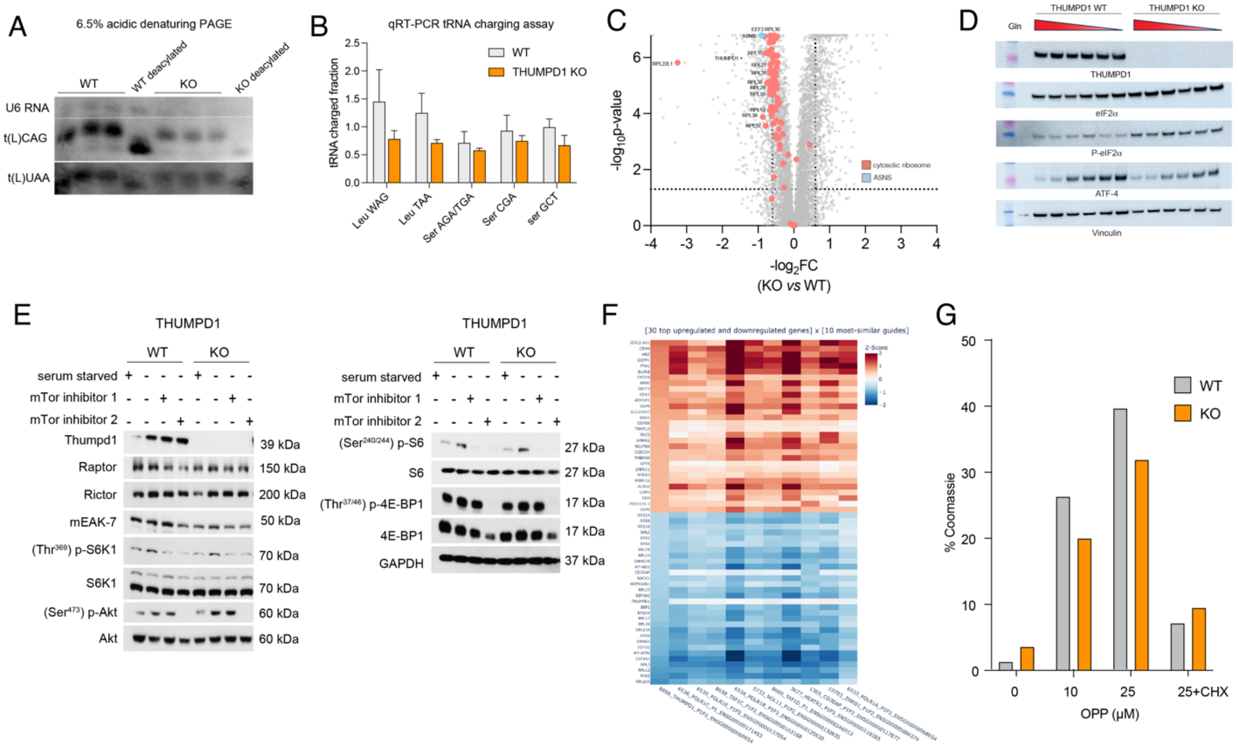


**Figure S4.** (a) Acidic denaturing PAGE analysis of tRNA charging in WT or THUMPD1 KO HEK-293T cells. Data are representative of n=2 biological replicates. (b) qRT-PCR analysis of tRNA charging in WT or THUMPD1 KO HEK-293T cells. Values for WT and KO tRNA paires were not significant as analyzed by two-tailed Student's *t* test ($P > 0.05$). Data are representative of n=2 biological replicates. (c) Ribosomal proteins (red) are downregulated in THUMPD1 KO HEK-293T cells. The ATF-4 target ASNS (blue) is also downregulated, suggesting the ISR is not activated by THUMPD1 KO. Values are derived from n=3 biological replicates. (d) THUMPD1 KO does not alter the threshold for glutamine-dependent activation of ATF-4. Glutamine concentrations (left to right): 2 mM, 0.2 mM, 0.02 mM, 0.002 mM, and no glutamine. (e) THUMPD1 KO cells do not show altered ability to activate mTOR signaling. Treatment conditions as follows: serum starved = removal of all amino acids and serum from medium (1 h), mTor inhibitor 1 = 1 µM AZD2014 (1 h), mTor inhibitor 2 = 10 nM rapamycin (1 h). Data are representative of n=2 biological replicates. (f) Single-cell RNA-Seq signature of THUMPD1 KO cells observed by Perturb-seq analysis of Replogle *et al.*[31] (g) Gel densitometry analysis of fluorescence signal from treatment of THUMPD1 WT and KO HEK-293T with *O*-propargyl puromycin (OPP) followed by click chemistry to a fluorescent azide. The percent of the fluorescent signal relative to the Coomassie signal was calculated, and used to produce the relative values given in Fig. 4g. Data are representative of n=2 biological replicates.

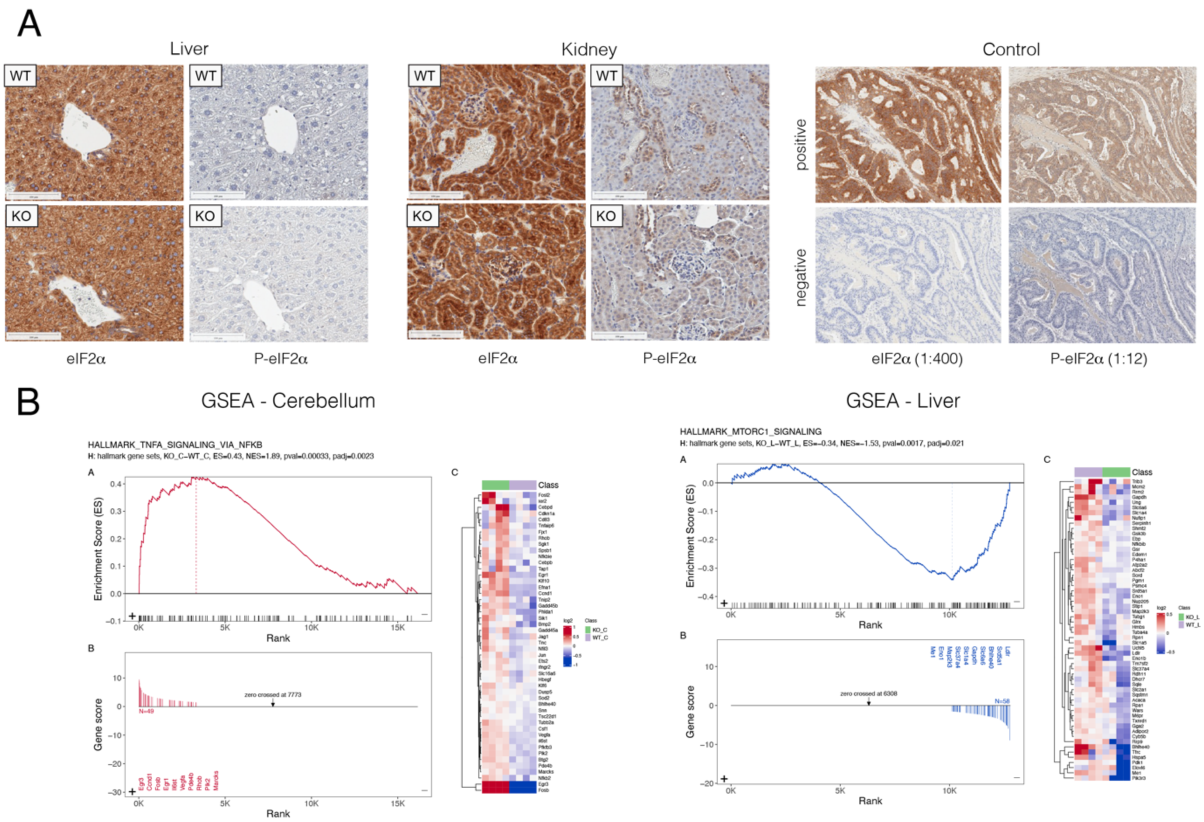


**Figure S5.** (a) Immunohistochemical (IHC) staining of total eIF2α (left) and (Ser$^{50}$) P-eIF2α (right) in liver tissue and kidney tissue isolated from age-matched WT and *Thumpd1*$^{-/-}$ KO mice. Results are representative of n=4 biological replicates. (b) Gene Set Enrichment Analysis indicating activation of inflammatory gene expression in mouse cerebellum (TNFA_SIGNALING_VIA_NFKB, left) and downregulation of transcripts associated with mTOR signaling (MTORC1_SIGNALING, right) in mouse liver. Pathway analyses were generated from RNA-Seq data (n=4 biological replicates). Additional pathway analyses are provided in Supplementary Tables 8-9.

A

| | Gcn2 WT | | | Gcn2 +/- | | | Gcn2 KO | | |
|---|---|---|---|---|---|---|---|---|---|
| | *Gcn2* WT, *Thumpd1 WT* | *Gcn2* WT, *Thumpd1*$^{+/-}$ | *Gcn2* WT, *Thumpd1*$^{-/-}$ | *Gcn2*$^{+/-}$, *Thumpd1 WT* | *Gcn2*$^{+/-}$, *Thumpd1*$^{+/-}$ | *Gcn2*$^{+/-}$, *Thumpd1*$^{-/-}$ | *Gcn2*$^{-/-}$, *Thumpd1 WT* | *Gcn2*$^{-/-}$, *Thumpd1*$^{+/-}$ | *Gcn2*$^{-/-}$, *Thumpd1*$^{-/-}$ |
| total | 26 | 46 | 11 | 36 | 73 | 21 | 12 | 25 | 12 |
| % actual | 10% | 18% | 4% | 14% | 28% | 8% | 5% | 10% | 5% |
| % expected | 6.25% | 12.50% | 6.25% | 12.50% | 25.00% | 12.50% | 6.25% | 12.50% | 6.25% |

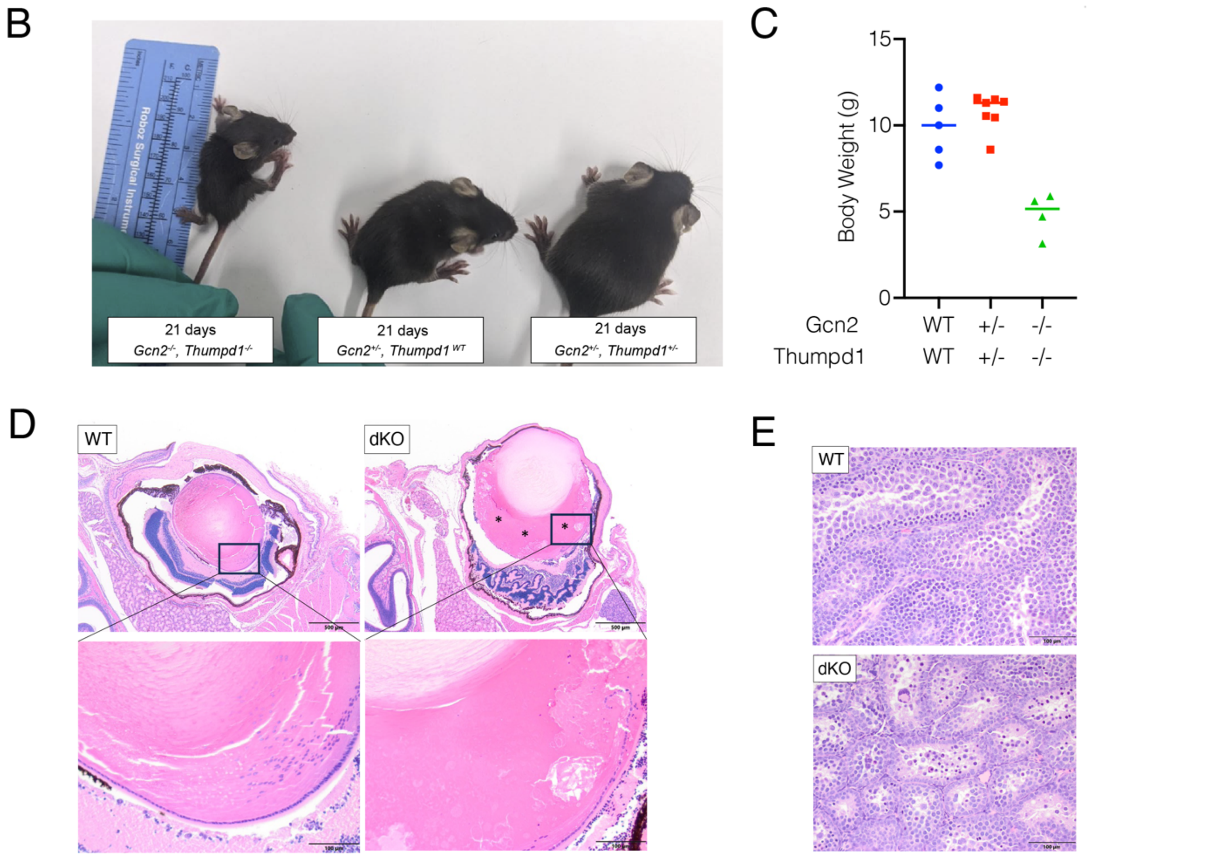


**Figure S6.** (a) Offspring annotated by genotype produced by *Thumpd1*$^{+/-}$,*Gcn2*$^{+/-}$ dihybrid cross. (b-c) *Thumpd1*/*Gcn2* double KO mice are runted. (d) A *Thumpd1*$^{-/-}$/*Gcn2*$^{-/-}$ double knockout (DKO) animal exhibits cataractous change where the lens is expanded by a liquefaction of lens fibers (*), which lack organization and are swollen and fragmented, often forming globules of degenerate lens proteins (Morgagnian globules). Hyperplasia of the lens epithelium is also observed. The animal also exhibits retinal dysplasia, where the retina is disorganized, poorly developed, and thrown into folds. There is retinal detachment with hypertrophy (tomb-stoning) of the retinal pigmented epithelium. (e) *Thumpd1*$^{-/-}$, *Gcn2*$^{-/-}$ DKO mice exhibit multifocal seminiferous tubule degeneration with syncytial cell formation. Due to the challenge of isolating *Thumpd1*$^{-/-}$, *Gcn2*$^{-/-}$ DKO mice prior to lethality and autoloysis onset, data for d-e are representative of n=1individual mouse.

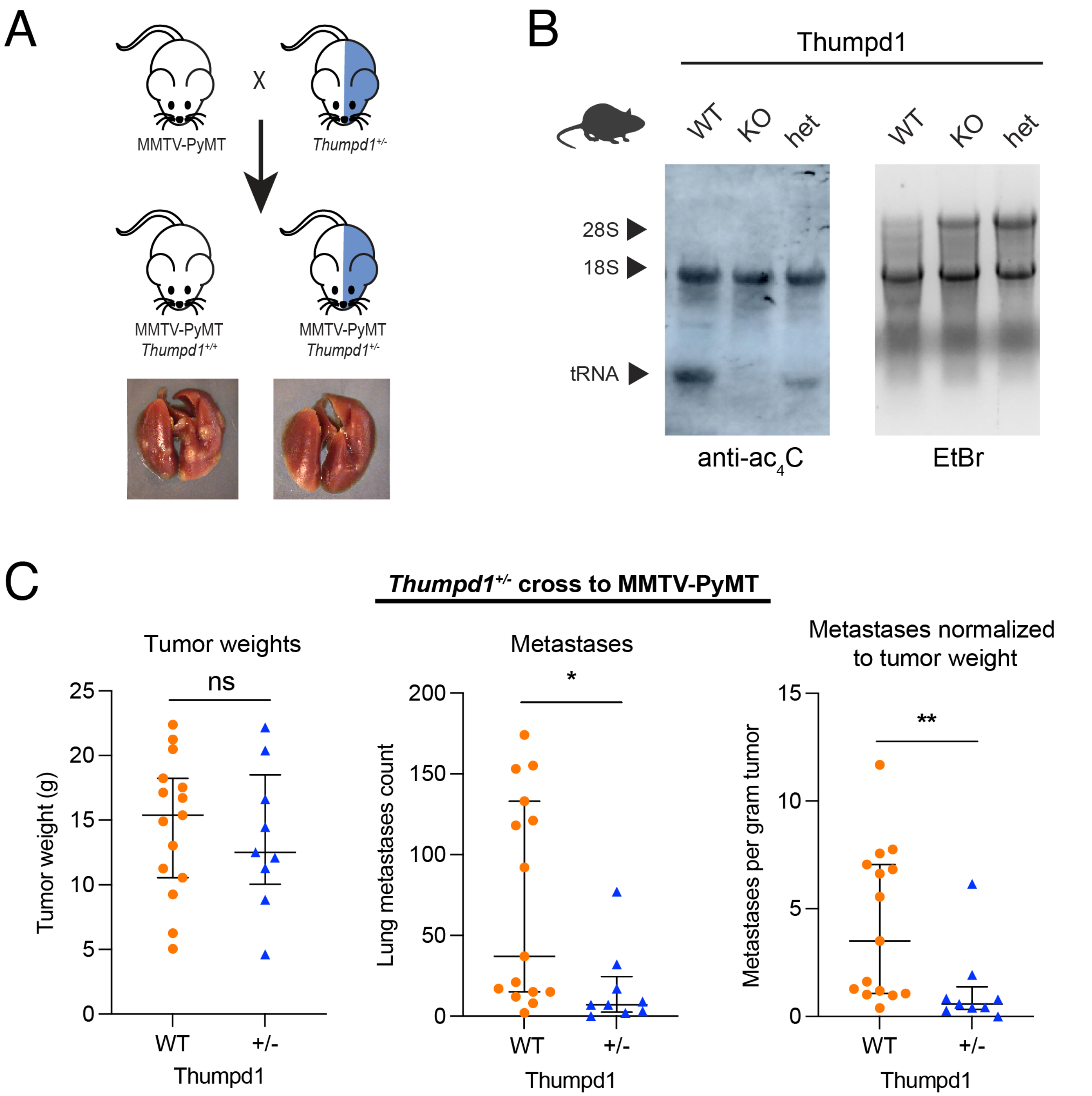


**Figure S7**. (a) Schematic for PyMT tumor metastasis model. (b) *Thumpd1*$^{+/-}$ heterozygotes exhibit qualitatively decreased tRNA acetylation as assessed by immuno-Northern blot. (c) *Thumpd1*$^{+/-}$ heterozygosity suppresses breast cancer metastasis but not primary tumor growth in PyMT model.

**KEY RESOURCES TABLE**

| REAGENT/RESOURCE | SOURCE | IDENTIFIER |
|---|---|---|
| **Antibodies** | | |
| Anti-ac$^4$C antibody | Abcam | Cat#: ab252215; RRID: AB_2827750 |
| Thumpd1 antibody | Bethyl | Cat#: A304-385A-M; RRID: AB_2620838 |
| eIF2α antibody | Santa Cruz | Cat#: sc-133227; RRID: AB_2096505 |
| Total eIF2a | Abcam | Cat#: ab169528; RRID: AB_2819002 |
| P-eIF2α antibody | Abcam | Cat#: ab32157; RRID: AB_732117 |
| Phospho-eIF2a | CST | Cat#: 3398; RRID: AB_2096481 |
| ATF-4 antibody | Cell Signaling | Cat#:11815S; RRID: AB_2616025 |
| Anti-rabbit IgG HRP-linked antibody | Cell Signaling | Cat#: 7074S; RRID: AB_2099233 |
| Anti-mouse IgG HRP-linked antibody | Cell Signaling | Cat#: 7076S; RRID: AB_330924 |
| α-mouse IgG HRP conjugate | Promega | Cat#: W4021; RRID: AB_430834 |
| Mouse monoclonal custom antibodies targeting endogenous mEAK-7 | Genscript (unpublished) | |
| Vinculin | Bethyl | Cat#: A302-535A-T; RRID: AB_2728768 |
| Raptor | Bethyl | Cat#: A300-553A; RRID: AB_2130793 |
| Rictor | Bethyl | Cat#: A300-459A; [RRID: AB_2179967 |
| S6K1 | CST | Cat#: 2708S; RRID: AB_390722 |
| (Ser$^{473}$) p-AKT | CST | Cat#: 4058S; RRID: AB_331168 |
| Total AKT | CST | Cat#: 4685S; RRID: AB_10698888 |
| S6 ribosomal protein | CST | Cat#: 2217S; RRID AB_331355 |
| (Ser$^{235/236}$) p-S6 ribosomal protein | CST | Cat#: 4858S; RRID: AB_916156 |
| (Ser$^{240/244}$) p-S6 ribosomal protein | CST | Cat#: 2215S; RRID: AB_331682 |
| (Ser$^{65}$) p–4E-BP1 | CST | Cat#: 9451S; RRID: AB_330947 |
| (Thr$^{37/46}$) p–4E-BP1 | CST | Cat#: 9459S; RRID: AB_330985 |
| (Thr$^{389}$) p-p70 S6 kinase | CST | Cat#: 9234S; RRID: AB_2269803 |

| | | |
|---|---|---|
| 4E-BP1 | CST | Cat#:9452S; RRID: AB_331692 |
| GAPDH | CST | Cat#:2118S; RRID: AB_561053 |
| **Software and algorithms** | | |
| Cutadapt v4.2 | Martin 2011[1] | RRID:SCR_011841 |
| BBMap v38.90 | (Bushnell et al., 2017)[2] | RRID:SCR_016965 |
| bowtie2 v2.3.5.1 | (Langmead & Salzberg, 2012)[3] | RRID:SCR_016368 |
| samtools v1.20 | (Danecek et al. 2021)[4] | RRID:SCR_002105 |
| R v4.2.0 | (R Core Team, 2023)[5] | RRID:SCR_001905 |
| ImageJ software (version 1.53t August 2022) | Peterson, T. (2010)[6] | |

### Experimental procedures

#### Animal models

*Thumpd1*$^{+/-}$ mice were generated by introduction of Cas9 protein and two synthetically modified guide RNAs (Synthego) into C57BL/6N fertilized eggs by microinjection. The synthetic guide RNAs were designed using sgRNA Scorer 2.0[7] to target 2 cuts, one in Exon 1 and one in exon 4. *Gcn2*$^{+/-}$ mouse was previously described[8] and obtained from Jackson labs (#008240). All mice were maintained and backcrossed on C57/BL6 background. Mice were housed at 25 °C in a 12 h light and 12 h dark cycle. Mice were 6-12 weeks of age and of either sex, with age-matched littermates used as controls unless noted otherwise. Following euthanasia, tissues were isolated, flash-frozen in RNAlater using liquid nitrogen, and stored at −80 °C until experiments were conducted. Frederick National Laboratory for Cancer Research is accredited by AAALAC International (Association for Assessment and Accreditation of Laboratory Animal Care) and follows the NIH Public Health Service Policy for the Care and Use of Laboratory Animals. All animal exp eriments were approved by the NCI-Frederick Institutional Animal Care and Use Committee (Approval ID: 24-437).

#### Genotyping

Tail DNA from Thumpd1 mice were isolated using NaOH extraction from 10-12 day old mice and subjected to polymerase chain reaction (PCR). PCR primer pairs used for genotyping of were D9 (GCTCGTTCATGTTGCAGGTG) and E9 (CAAACTGTCGCGTACGTGTG) which amplify a short segment spanning the start of exon 1, D7 (TGAGACCACCTTCCCACAGA) and E7 (GAGTGCAAAAGACTCGCAGC) which amplify a short segment spanning the end of exon 4, and D7/E9 which amplify a product only when the segment between the guides is deleted. Genotypes were analyzed by agarose gel electrophoresis for routine genotyping, while MiSeq analysis was used to sequence and identify breakpoints.

#### Construction of HEK-293T *Thumpd1* knockout cells

*Thumpd1* knockout cells were generated using the CRISPR/Cas9 system. Briefly, single guide RNAs (sgRNAs) targeting protein-coding sequence of human *Thumpd1* were designed using sgRNA Scorer 2.0.[7] Oligonucleotides containing the 20-nucleotide spacer sequence, along with appropriate 5′ overhangs, were annealed by mixing equal quantities (~50 pmol) of the forward and reverse oligonucleotides, heating for 2 min at 95 °C, and cooling in steps of 5 °C for 2 min duration until a final temperature of 25 °C. Each pair of annealed oligonucleotides was ligated into the BbsI site of the pX458 plasmid.[9] SpCas9(BB)-2A-GFP (pX458) was a gift from Feng Zhang (48138 Addgene plasmid; http://n2t.net/addgene:48138). Ligated plasmids then were transformed into the Stbl3 *Escherichia coli* strain, and colonies were grown out for large-scale plasmid preparation. Purified plasmids expressing *Thumpd1* guide 1a and 1b were cotransfected (∼1 μg) into HEK-293T (WT) cells using Lipofectamine LTX (Thermo Fisher Scientific, 15338030), and bulk green fluorescent protein-positive cells were then sorted using a flow cytometer and grown for approximately 5 days. After assaying for protein expression by immunoblotting, candidate clones were further subjected to single-cell sorting. Confirmation of gene editing for *Thumpd1* was determined by Sanger sequencing of DNA, RNA-seq, anti-ac$^4$C immuno-Northern blotting, and protein expression in whole-cell extract.

#### *Thumpd1*$^{+/-}$ cross to MMTV-PyMT metastasis model

Male FVB/N-Tg(MMTV-PyVT)634Mul/J (RRID:MGI:3032640)] mice were bred to 6-8 week old Thumpd1 +/- heterozygous female animals to generate F1 offspring. Genotyping of female animals was performed by PCR,[10] using DNA isolated from tail biopsies taken at weaning. MMTV-PyMT positive female animals were aged until experimental endpoint at 120 days of age. Lungs and tumors were harvested, and tumor weight and pulmonary surface metastases determined for each animal after euthanasia by a single investigator blinded to Thumpd1 status. Statistical significance between the two Thumpd1 genotypes was calculated using Mann Whitney U tests in GraphPad Prism. All animal experiments were approved by the NCI-Bethesda Institutional Animal Care and Use Committee (Approval ID: LPG-002).

#### Histopathology and bloodwork

A full necropsy was performed to determine the spectrum of pathology in Thumpd1 KO mice. Organ weights were recorded prior to fixation fore heart, kidney, liver, brain, lung, and spleen. All tissues were fixed in 10% NBF for 72 h and routinely processed for H&E staining and microscopic examination by a board-certified pathologist. At necropsy, blood was taken via cardiac puncture for complete blood counts (CBC), blood smear preparation, and clinical chemistry using Genesis hematology and Abaxis VetScan VS2 (Zoetis) analyzers. Full histopathology was performed on male and female WT (n = 6), ThumpD1 KO (n = 6), and ThumpD1/Gcn2 double KO (n = 4) mice.

**Growth conditions of HEK-293T THUMPD1 WT and KO cells**

HEK-293T cells for THUMPD1 WT, KO or rescue were grown in DMEM (Quality Biological, 68101520) supplemented with 10% FBS (Avantor Seradigm, 97068-085), 2 mM L-glutamine (Thermo Fisher, 25030081), and 1% penicillin-streptomycin (Thermo Fisher, 15140122). Cells were grown 37 °C under 5% $CO_2$ and passaged at 80-90% confluence. Cells tested negative for mycoplasma with the LookOut MycoPlasma PCR Detection Kit following manufacturer's instructions (Sigma-Aldrich, MP00035).

**Extraction of total RNA from mammalian cells**

HEK-293T Cells were grown until ∼80–90% confluency before harvesting. Cells were harvested by either scraping or by the addition of trypsin–EDTA, and centrifuging at 500 rcf, 2 min at room temperature. Pelleted cells were washed twice with cold PBS, and pellets were stored frozen at −80 °C. Total RNA from human cells was extracted using TRIzol according to the manufacturer's protocol. One milliliter TRIzol was used per $1 \times 10^7$ cells. The RNA pellet was resuspended in water and stored at −80 °C. Isolated total RNA was incubated with Turbo DNase (Invitrogen, AM2238) for 30 min at 37 °C to remove any DNA contamination. Typical extractions were carried out with $1 \times 10^7$ cells and yielded 400 µg of total RNA. Quality of the RNA was assessed by using the Agilent 2100 Bioanalyzer with RNA 600 nano kit (Agilent, 5067-1511) and the concentration was determined by the Nanodrop or Qubit fluorimeter.

**Extraction of total RNA from Thumpd1 mouse tissues**

The respective mouse organs were extracted from wild-type (WT), heterozygous (*Thumpd1*$^{+/-}$), and knockout (KO, *Thumpd1*$^{-/-}$) mice and were stored in RNAlater™ Stabilization Solution (Thermo Fisher Scientific, AM7020) at -80 °C. Based on the tissue type, portions of 50-100 mg were cut from the whole organ for RNA isolation. The tissues were minced into small pieces while soaked in RNA stabilizing agent, as this improves penetration of RNA stabilizing agent and retains integrity of RNA in the tissues. The minced tissues were then transferred to a 2.0 mL prefilled (1.5 mm) Zirconium homogenizer beads tube (Stellar Scientific BS-BEBU-215) as quickly as possible while maintaining cold conditions. The bead tubes were prefilled with 1 mL of chilled TRI reagent from the Direct-zol RNA Miniprep kit (Zymo Research, R2051). Samples were homogenized by beat beating with BeadBug Microtube Homogenizer at 400 rpm. The samples in the bead tubes were given pulses 4 times for 30 s, incubating on ice for 30 s between each beating. To remove particulate debris from homogenized samples, the samples were left on ice for 5 min and centrifuged at 21000 rcf for 3 min. The supernatants were transferred to nuclease-free 1.7 mL Eppendorf tubes. Total RNA was purified using Direct-zol RNA Miniprep kit following the manufacturer's protocol (Zymo Research, R2051) and total RNA was eluted in 50 uL of nuclease-free water. Total RNA was quantified using a nanodrop and the quality was checked using Agilent bioanalyzer (Agilent RNA 6000 Nano Kit, 5067-1511). The total RNA was stored at −80 °C until use.

**Extraction of proteins from Thumpd1 mouse tissues**

The respective mouse organs were extracted from wild-type (WT), heterozygous (*Thumpd1*$^{+/-}$), and knockout (KO, *Thumpd1*$^{-/-}$) mice and were stored in RNAlater™ Stabilization Solution (Thermo Fisher Scientific AM7020) at -80 °C. Portions of 50 mg of each tissue were cut from the whole organ for protein extraction. The tissues were minced into small pieces as quickly as possible while maintaining cold conditions. The minced tissues were then transferred to a 2.0 mL prefilled (1.5 mm) Zirconium homogenizer beads tube (Stellar Scientific BS-BEBU-215). The bead tube was prefilled with 500 uL of chilled TPER tissue protein extraction reagent (Thermo Scientific, 78510) with freshly added x1 Protease Inhibitor Cocktail (Cell Signaling, 5871). Samples were homogenized by beat beating with BeadBug Microtube Homogenizer at 400 rpm. The samples in the bead tubes were given pulses 4 times for 30 s, incubating on ice for 30 s between each beating. To remove particulate debris from homogenized samples, the samples were left on ice for 5 min and centrifuged at 10,000 rcf for 5 min. The supernatants were transferred to clean tubes and were sonicated on ice using 700W QSonica Q125 sonicator with the 1/16" microtip (2 × 5 sec pulse, 20% amplitude, 10 sec resting on ice between pulses). Samples were centrifuged at 21000 rcf for 30 min at 4 °C. The supernatant containing the proteins was collected leaving behind cell debris. Protein quantification was done using a Pierce™ BCA Protein Assay Kit (Thermo Scientific, 23225) following the manufacturer's protocol. The proteins were stored at −80 °C until use.

**Immuno-Northern blotting of ac$^4$C**

Total RNA was isolated from cells or mouse tissues as described above and quantified using the Qubit RNA BR assay kit (Thermo Fisher Scientific). Immuno-northern blots were performed using Invitrogen Northern-Max reagents (Thermo Fisher Scientific). Same amount of RNA (15 µg) from each condition were aliquoted and mixed with 1 vol of NorthernMax-Gly Sample Loading Dye (Thermo Fisher Scientific, AM8551). These were then incubated at 65 °C for 30 min and separated on a 1% agarose-1X Glyoxal Gel prepared using 10X NorthernMax-Gly Gel Prep/Running Buffer (Thermo Fisher Scientific, AM8678). Gels were run at 80 V for approximately 70 min, or until the dye front had migrated about 3 inches. Loading controls were analyzed by imaging of ethidium bromide before transfer. RNA was transferred onto Amersham Hybond-N+ membranes (Cytiva, RPN119B) using a downward capillary method as described previously.[11] After transfer, membranes were crosslinked three times at 150 mJ/cm$^2$ in a $UV_{254nm}$ Stratalinker 2400 (Stratagene). Membranes were then blocked with 5% non-fat milk in 0.1% TBST for 30 min at room temperature and washed 3 times at 5 min each in 0.1% TBST. Membranes were then incubated overnight at 4 °C with the anti-ac$^4$C antibody (Abcam, ab252215 [RRID: AB_2827750]; 1:2000 dilution) in blocking buffer (5% non-fat milk in 0.1% TBST). Membranes were washed 3 times at 5 min in 0.1% TBST and then incubated with HRP-conjugated secondary anti-rabbit IgG (Cell Signaling Technology, 7074 [RRID: 2099233]; 1:10000 dilution) in 5% non-fat milk for 1 h at room temperature. Membranes were washed 3 times at 10 min each in 0.1% TBST. SuperSignal ELISA Femto Maximum Sensitivity Substrate reagent (Thermo Fisher Scientific, 37075) was added directly to the membrane and signal was detected via chemiluminescent imaging using an Amersham ImageQuant 800 (Cytiva, 29399482).

**Mim-tRNAseq**

Small RNA enrichment total RNA

Total RNA was extracted from HEK-293T cells or mouse tissues as mentioned above. Small RNA enrichment was done according to the manufacturer's protocol using Quick-RNA Microprep Kit (Zymo Research, R1050) using 10-30 ug of purified total RNA as the starting material. Small RNA was eluted in 15-30 uL of nuclease-free water and was quantified using a nanodrop. The small RNA was stored at −20 °C until further use.

Small RNA demethylation treatment using ALKBH1

For efficient and quantitative tRNA sequencing, demethylation of small RNA was carried out to remove base methylations such as N1-methyladenosine (m1A), N3-methylcytosine (m3C) and N1-methylguanosine (m1G) present in tRNAs. The demethylation assay was done using recombinant D135S AlkB mutant enzyme expressed in house following the method used by Zheng *et al.*[12] Small RNA substrate (40 pmol) was reacted with 80 pmol of ALKBH1 enzyme in a 100 μL reaction containing 300 mM KCl, 2 mM $MgCl_2$, 2 mM ascorbic acid, 300 μM alpha-ketoglutaric acid, and 50 μM ammonium iron (II) sulfate. The mixture was incubated at room temperature for 2 h and immediately quenched by adding EDTA to final concentration of 5 mM. RNA was purified by the addition of 300 mM Sodium Acetate, 1 mM EDTA, 0.1 U/μL Superase-In (Invitrogen, AM2694) followed by isopropanol precipitation on dry ice for 30 minutes and spinning at 21000 r.c.f, 4°C for 30 min.

Mim-tRNAseq library preparation

Demethylated small RNA (<200 nt) samples from cells or mouse tissues were used for library preparation. RNA dephosphorylation was performed with 5U of T4 PNK (NEB, M0201L) at 37 °C for 1 hour in the presence of T4 PNK buffer without ATP (NEB, B0201S). For tRNA sequencing libraries a previously described 5'-phosphorylated, pre-adenylated adapter (oBZ407) with six randomized nucleotides at the 5′-end and a 3′ blocking group (/5Phos/AppNNNNNNCACTCGGGCACCAAGGA/3ddC)[13] was used. oBZ407 adapter was ligated to the RNA template using a truncated KQ T4 RNA ligase 2 (NEB, M0242L) for 1 h at 37 °C in the presence of 50% PEG 8000 (NEB, B0216). Ligated RNA was gel-purified and size-fractionated on a Criterion Precast TBE–urea 10% denaturing polyacrylamide gel (BioRad, 3450088) to enrich tRNA molecules using two markers of 40 and 80 nt and the RiboRuler low-range ssRNA ladder (ThermoScientific, SM1831). Gels were stained using 1× SYBR Gold nucleic acid stain (10,000×; Invitrogen, S11494) in 1× TBE for 3 min, and 3′-adapter-ligated RNA in size range of 60-100 nt was excised. Small RNA was recovered from excised polyacrylamide gel pieces by crush and soak method followed by isopropanol precipitation with 300 mM Sodium acetate, 1 mM EDTA, 0.1 U/μL Superase-In (Invitrogen, AM2694) overnight at 4°C and 1.5X volume of 100% isopropanol. Next, reverse transcription was carried out using MarathonRT reverse

transcriptase (Kerafast, EYU007) with RT primer oBZ408 (/5Phos/ RNNNAGATCGGAAGAGCGTCGTGTAGGGAAAGAGTGTAGATCTCGGTGGTC GC/iSP18/TTCAGACGTGTGCTCTTCCGATCTGTCCTTGGTGCCCGAGTG). Template RNA was subsequently hydrolyzed by the addition of 1 µl of 5M NaOH and incubation at 95 °C for 3 min. Reaction products were separated from unextended primer on Criterion Precast TBE–urea 10% denaturing polyacrylamide gel (BioRad, 3450088). Gels were stained with SYBR Gold, the region between 60 and 100 nt was excised. Circularization of purified cDNA with Circ Ligase (Biosearch Technologies, CL9021K) was carried out in 1X reaction buffer supplemented with 1 mM ATP, 50 mM $MgCl_2$ for 2 h at 60 °C, followed by enzyme inactivation for 10 min at 80 °C. Then the DNA was PCR amplified using Phusion high-fidelity PCR master mix (Fisher Scientific, F531L) for 10-12 cycles of 10 sec at 98 °C, 10 sec at 65 °C, and 5 sec at 72 °C with 5'-AATGATACGGCGACCACCGAGATCTACAC-3'
and 5'-CAAGCAGAAGACGGCATACGAGAT
[8nt barcode] GTGACTGGAGTTCAGACGTGTGCTCTTCCG-3' primers. PCR products were equimolarly pooled for cluster generation with additional size selection between 190-250 nt. The quality of the sequence libraries, size, purity, and concentration was validated using Agilent high-sensitivity DNA 1000 kit (Agilent Technologies, 5067-4626). Sequencing was performed as single-end reads for 100 cycles on a NextSeq machine (Ilumina).

**Analysis of tRNA-Seq libraries and misincorporation**

tRNA abundance was quantified using modification-induced misincorporation tRNA sequencing (mim-tRNA Seq) package (https://github.com/nedialkova-lab/mim-tRNAseq). Scripts are available at https://github.com/CCBR/TRANQUIL.

**Ribosome and disome profiling**

HEK293 WT and Thumpd1 KO cells were grown in 10-cm plates. Cells were washed with PBS once and collected with 1 ml footprint lysis buffer [20mM Tris-Cl (pH8.0), 150mM KCL, 5 mM MgCl2, 1mM DTT, 1% (v/v) Tritonx100, 0.1 mg/ml Cycloheximide] with vigorous scrapping. Lysates were digested with DNase I (2 units per mL) (Thermo Fisher Scientific, AM2222) for 15 min on ice and clarified by centrifugation at 15,000 rpm for 15 min at 4 °C. Lysates containing 20 µg of total RNA were digested with 750 units of RNase I (Thermo Fisher Scientific, AM2295) at

25 °C for 1 h with gentle shaking at 500 rpm and quenched by adding 200 units of SUPERaseIn (Thermo Fisher Scientific AM2694, 20U/µL). Nuclease-treated lysates were layered on 0.9 ml sucrose cushion [20 mM Tris-Cl (pH8), 150 mM KCl, 5 mM MgCl2, 1 mM DTT, 1 M sucrose]. Ribosomes were pelleted by centrifugation in a TLA100.3 rotor at 100,000 rpm for 1 hr at 4 °C and RNA was extracted by miRNeasy mini kit (QIAGEN). Footprints between 25-40 nt and 40-80 nt were size-selected separately for monosome and disome library preparation, respectively. Procedures for library construction were as described previously.[13]

Analysis of ribosome and disome profiling data

hg19 reference genome assembly from UCSC was used for human genome alignment. A human transcriptome file was generated to include canonical transcripts of known genes from UCSC genome browser. Libraries were trimmed to remove 3' adapter (NNNNNNCACTCGGGCACCAAGGA), and 4 random nucleotides included in RT primer (RNNNAGATCGGAAGAGCGTCGTGTAGGGAAAGAGTGTAGATCTCGGTGGTCGC/iSP18/TTCAGACGTGTGCTCTTCCGATCTGTCCTTGGTGCCCGAGTG) were removed from the 5' end of reads. Trimmed reads were aligned to human ribosomal and non-coding RNA sequences using STAR[14] with '-outFilterMismatchNoverLmax 0.3'. Unmapped reads were mapped to the human transcriptome file with '–outFilterIntronMotifs RemoveNoncanonicalUnannotated –outFilterMultimapNmax 1 –outFilterMismatchNoverLmax 0.1' as previously described[15]. All other analyses were performed using software custom written in Python 3.10 and R 4.3. Scripts are available at— https://github.com/NCI-RBL/Dockers/tree/main/workflows/RiboFootPrint.

**Leucine and serine repeat GFP reporter assays**

Construction of Leucine and Serine Repeat Reporter Plasmids

Site-directed mutagenesis was used to insert Leu or Ser codon repeats (see Table S1 for primer list and insertions) into the *N*-terminus of a pmaxGFP™ (Lonza Bioscience) reporter construct.

To generate each insertional GFP mutant construct, PCR reactions were carried out with WT pmaxGFP™ vector as a template, the pair of corresponding primers, and Q5 DNA polymerase (New England Biolabs, M0493S). Reactions were carried out with hot start step at 92 °C for 2 min then for 18 cycles with denaturing step at 95 °C for 30 sec, annealing step at 54 °C for 30 sec and polymerization step at 72 °C for 1 min 45 sec followed by reactions incubation at 72 °C for 5 min before holding at 4 °C. Then DpnI (New England Biolabs, R0176S) was added to PCR reactions and incubated for 1 h at 37 °C to digest template DNA followed by enzyme deactivation at 75 °C for 10 min. DpnI-treated PCR reactions were used to transform chemically competent XL1-Blue cells (Agilent). The resulted colonies were selected on LB plates containing 50 µg/mL kanamycin and plasmid DNA was prepared LB liquid cultures containing 50 µg/mL kanamycin using I-Blue Mini Plasmid Kit (IBI Scientific, IB47170). Plasmids containing the correct insertions of Leu or Ser codons (as determined by sequencing using a CMV promoter primer, Table S1) were selected for GFP assays.

**GFP reporter assays.**

Wild-type pmaxGFP™ vector and pmaxGFP™ vectors containing upstream Leu or Ser codon repeats were transiently transfected into HEK-293T cell line and the corresponding *THUMPD1* knockdown cell lines using jetPRIME® transfection reagent (Polyplus, 101000046) as suggested by manufacturer, in 96-well dishes in triplicates. Specifically, 100 ng of plasmid DNA samples in 10 µL of jetPRIME® buffer was mixed with 0.2 µL of jetPRIME® reagent, incubated 10 min at RT and added to each well containing $1x10^4$ cells in standard DMEM medium (Corning). EGFP fluorescence in live cells was measured at 48 h post-transfection time in SPARK microplate reader (Tecan), excitation 482 nm and emission 512 nm. The GFP reporter expression was calculated as the ratio of EGFP fluorescence sample values (in triplicates) normalized to the corresponding values of the sample of WT pmaxGFP™ vector transfected into HEK-293T (set to the value of 1) with the standard deviation calculated for each sample.

**Western blotting for analysis of eIF2α phosphorylation**

HEK-293T THUMPD1 WT/KO cells were plated at 500,000 cells/well in 6-well plates in 2 mL complete media and allowed to adhere overnight. For ISRIB treatment and complete deprivation of glutamine, media was aspirated 24 hours after plating cells, 1500 µL complete media with glutamine was replaced, and cells were treated dropwise with 500 µL media containing ISRIB (Cayman Chemical, 16258) for a final concentration of 1.1 µM ISRIB with 0.01% DMSO. Control wells were treated with 0.01% DMSO vehicle. Cells were returned to the incubator for 30 minutes, and media was then aspirated. For wells indicated for glutamine starvation, cells were washed twice with media lacking glutamine. Glutamine-containing or lacking media was replaced, and cells re-dosed with ISRIB/DMSO as described above, and cells were returned to the incubator for 6 hours. After 6 hours, cells were washed in 1 mL of 1X cold PBS, scraped in 500 μL PBS, transferred to a pre-chilled Eppendorf tube, centrifuged (500 rcf x 5 min), and pellets were snap frozen and stored at -80°C. Cells were lysed by sonication in 1x PBS supplemented with 1x protease/phosphatase inhibitor (Cell Signaling, 5872). For concentration-dependent glutamine deprivation, media was aspirated 24 hours after plating, cells were washed 1x with PBS and media was replaced with 2 mL media containing 2000, 200, 20, 2, 0.2, or 0 μM of L-glutamine. Cells were incubated for 3 hours before harvesting and lysing cells as described above.

Total protein quantification was performed using Precision Red Protein Assay (Cytoskeleton, ADV02). SDS-PAGE was performed using 4-12% Bis-Tris NuPAGE gels (Invitrogen, NP0322 and NP0323), with XCell SureLock Mini-Cells (Invitrogen, EI0002) and MES running buffer (Invitrogen, NP0002) according to manufacturer's protocols. Total protein (10 μg) was loaded per well, and BenchMark Pre-stained Protein Ladder was loaded on all gels (Invitrogen, 10748010). Gels were transferred by iBlot dry transfer (Invitrogen, IB1001) using nitrocellulose transfer stacks (Invitrogen, IB301001) at 20 volts for 1 min, 23 volts for 4 min, and 25 volts for 2 min. Total protein content was visualized using Ponceau stain after washing with 5% acetic acid in water. Membranes were blocked in StartingBlock (PBS) Blocking Buffer (Thermo Fisher, 37538) for 20 min at room temperature. Membranes were then probed overnight at 4°C with antibodies for Thumpd1 (Bethyl, A304-385A-M [RRID: AB_2620838]; 1:5000 or 1:2,000 dilution), eIF2α (Santa Cruz, sc-133227 [RRID: AB_2096505]; 1:1,000 dilution), P-eIF2α (Abcam, ab32157 [RRID: AB_732117]; 1:10,000 dilution), ATF-4 (Cell Signaling, 11815S [RRID: AB_2616025]; 1:1,000 dilution), or Vinculin (Bethyl, A302-535A-T [RRID: AB_2728768]; 1:10,000 dilution), with all dilutions made in StartingBlock Blocking Buffer. Secondary antibodies were either anti-rabbit IgG HRP-linked antibody (Cell Signaling, 7074S [RRID: AB_2099233] or anti-mouse IgG HRP-linked antibody (Cell Signaling, 7076S [RRID: AB_330924]) and were both incubated at 1:1000 dilutions in 5% non-fat dry milk in 1x TBST for 1 h at room temperature. Separate gels were run for eIF2α and P-eIF2α antibodies, but some membranes were re-probed with antibodies at different molecular weights. Membranes were washed at least 3 times with 1x TBST between antibodies and before imaging. Imaging of colorimetric and chemiluminescent signals was performed using an Amersham ImageQuant 800 (Cytiva, 29399482), and for chemiluminescent signal using Lumiglo (Cell Signaling Technology, #7003) or SuperSignal ELISA Femto Substrate (Thermo Scientific, 37074) according to manufacturer's protocols.

**Immunohistochemistry of eIF2a and eIF2a-P**

Tissue sections were stained on Leica Biosystems' BondMax autostainer with the following conditions: heat-induced epitope retrieval with EDTA for 20 minutes following by phospho-eIF2a (Cell Signaling Technology, 3398 [RRID: AB_2096481], rabbit monoclonal, 1:12) or total eIF2a (Abcam, ab169528 [RRID: AB_2819002], rabbit monoclonal, 1:400). Positive controls included human colon carcinoma and human lung carcinoma. Isotype negative controls involved replacing primary antibody with non-clonal, isotype-matched antibody from the same species as the primary antibody. Sections of brain, liver, and kidney were evaluated from WT (n=3), Thumpd1 KO (n=3), and Thumpd1/Gcn2 DKO (n=3). Slides were digitalized at 20× objective (0.5 × 0.5µm per pixel) using Aperio AT2 scanner (Leica Biosystems) and analyzed using HALO (Indica Labs, v3.6). Appropriateness of staining and regions of interest annotation were completed by a board-certified pathologist to include cerebral cortex, hippocampus, liver, and renal cortex. The percentage of positive pixels is reported.

***In situ* OPP labeling, cell culture maintenance and proteome harvesting for gel-based fluorescence readout**

HEK-293T WT and HEK-293T THUMPD1 cell lines were cultured as described above. For gel-based fluorescent detection of protein translation, three 6 cm diameter dishes (USA Scientific, CC7682-3359) were plated for each cell line with a plating density of 5.0 x $10^5$ cells. Each plate contained 5 mL of complete growth medium and returned to the cell incubator for propagation. Growth media was replaced every two days. Upon reaching 80 % confluency, 12.5 µL of 0-, 4-, and 10-mM OPP stocks dissolved in DMSO (Millipore Sigma, 276855) was added in a dropwise manner to each plate and swirled in a clockwise motion to ensure even distribution of OPP. Final OPP concentrations were 0, 10, and 25 µM respectively. To demonstrate a change in translation, a control condition was set up for each cell line that received a 15-minute preincubation with 12.5 µL of 72 mM cycloheximide (EMD Millipore, 239764-10MG) stock dissolved in DMSO for a final concentration of 180 µM before the 1 h incubation with 25 µM OPP. Plates were subsequently returned to the incubator for a 1 h incubation. Dosing of each plate was staggered by 10 min increments to ensure that harvesting was started after the 1 h incubation was completed. Following incubation, growth media was removed by aspiration and replaced with 1 mL of PBS. Cells were harvested by lifting cells from the plate's surface area with a cell lifter (VWR International, 75799-938). Once cells were dislodged from the plate, the PBS solution was transferred to a 15 mL conical, and the cell lifting process was repeated for a total of 2 times. Cells were centrifuged for 5 min at 500 rcf at 4 °C to form cell pellets. Following centrifugation, supernatant was aspirated, and cells were resuspended in 1 mL of ice-cold PBS and transferred to a 1.5 mL microcentrifuge tube. Cells were then pelleted by centrifugation for 5 min at 700g at 4 °C. Supernatant was subsequently aspirated and cell pellets were resuspended in 100 µL of lysis buffer consisting of 1X PBS supplemented with 1X protease inhibitor cocktail (Cell Signaling, 5871). Cell pellets were lysed by sonication using a 700W QSonica Q700 sonicator (15 × 2 sec pulse, amplitude 1, 30 sec resting on ice between pulses). Following sonication, lysates were centrifuged for 30 min at 4 °C at 21000 rcf. After centrifugation, supernatants were recovered, and protein concentration was determined by Precision Red Advanced Protein Assay (Cytoskeleton, AVD02) using the manufactures 96-well plate format. Lysates were subsequently diluted to a 1.3 mg/mL protein concentration using lysis buffer and stored at -80 °C.

<u>Fluorogenic detection of OPP labeled wild type and THUMPD1 knockout proteomes</u>

To assess global changes in the nascent proteome, proteins (68 µL, 1.3 mg/mL) labeled by OPP were visualized by SDS–PAGE through Cu(i)-catalyzed [3 + 2] cycloaddition (CuAAC) with a fluorescent azide as previously reported.[16] Briefly, 7 µL of a click chemistry master mixture consisting of TAMRA-azide (100 µM; 5 mM stock solution in DMSO; Millipore Sigma, 760757), TCEP (1 mM; 100 mM stock in 200 mM NaOH; Millipore Sigma, C4706-2G), Tris-(benzyltriazolylmethyl)amine ligand (TBTA; 100 µM; 1.7 mM stock in DMSO:tertbutanol 1:4; Millipore Sigma, 678937-50MG), and $CuSO_4$ (1 mM; 50 mM stock in H2O; VWR International, BDH9312-500G) was added to each labeled proteome. Reactions were vortexed and incubated at room temperature for 1 h in the dark. Reactions were vortexed every 20 min and returned to dark storage. Upon completion of the 1 h incubation, the cycloaddition reaction was quenched by addition of 8.35 µL of 1000 mM DTT (Millipore Sigma, D0632). Samples were then subjected to an acetone protein precipitation to remove cycloaddition reagents. Briefly, 187 µL of acetone

(Fisher Scientific, A1320) was added to each sample and incubated for 2 min at room temperature. Samples were then pelleted by centrifugation at 20,000 rcf at room temperature. Following centrifugation, the supernatant was removed and allowed to air dry for 1 min. Protein pellets were resuspended in loading buffer consisting of 4X LDS-loading buffer (Invitrogen, NP0007) and 100 mM DTT. Subsequently, samples were incubated at 95 °C for 10 min to dissolve protein pellet. 15 µL of each sample was analyzed by gel electrophoresis using Bis–Tris NuPAGE gels (4–12%, Invitrogen, NP0322), and MES running buffer (Life technologies, NP0002) in Xcell SureLock MiniCells (Invitrogen) according to the manufacturer's instructions. The Xcell SureLock MiniCells electrophoresis apparatus was connected to a PowerEase Touch 120W Power Supply (Thermo Fisher Scientific, PS0120) at 200 volts for 35 min. Gels were fixed and destained in a solution of 50/40/10% MeOH/$H_2O$/AcOH overnight to remove excess probe fluorescence, rehydrated with water for 30 min, and visualized using an Amersham ImageQuant 800 imager with IQ800 Control Software version 1.2.0.

Densitometry analysis of OPP labeled proteomes

Densitometry analysis to normalize fluorescence signal to the protein loading control was done as previously reported using ImageJ software (version 1.53t August 2022).[6] Briefly, ".PNG" images of the OPP labeled proteomes and Coomassie loading controls were uploaded to ImageJ. For a given sample, a rectangle was created to encompass a whole lane, and this process was done for all lanes analyzed. Once all lanes were selected, the lanes were analyzed using the "plot lanes" function to create a histogram of every lane. Next, the "tool" function was selected to enclose each histogram prior to selecting the "magic wand" function to integrate the area under the curve and create a "OPP fluorescence" densitometry value. This process was repeated for the Coomassie image to create a corresponding whole lane "loading control" densitometry value. The fluorescence densitometry values were normalized using the following equation: $\% \, Coomassie = (loading \, control) \times (100)$. The normalized data was then plotted using GraphPad software (Version 10.2.3 (347), April 21, 2024).

**RT-qPCR based analysis of Thumpd1 mRNA**

Mouse liver tissue total RNA was isolated and quantified using Nanodrop as described above. RT-qPCR was performed using Luna Universal One-step RT-qPCR kit (New England Biolabs,

E3005S) using an LightCycler® 480 II PCR instrument (Roche). Real-time instrument was programmed with the following thermocycling protocol:

| Cycle step | Temperature | Time | Cycle |
|---|---|---|---|
| Reverse transcription | 55 °C | 10 min | 1 |
| Initial denaturation | 95 °C | 1 min | 1 |
| Denaturation<br>Extention | 95 °C<br>60 °C | 10s<br>30s | 45 |
| Melt curve | 60-95 °C | various | 1 |

The cycle threshold values (Ct) were obtained from the LightCycler 480 SW 1.5.1 software. Thumpd1 primers used for the RT-qPCR of the total RNA from Thumpd1 WT, KO, and het liver tissues are listed in Table S1.

Melt curves were also performed to confirm the presence of a single amplicon by RT-PCR and the absence of primer dimer. The heterogeneous and Thumpd1 KO samples were normalized to the wildtype mouse using $2^{-\Delta\Delta C_t}$ relative abundance formula. The data is represented in technical triplicates (n=3).

**Sanger sequencing analysis of ac$^4$C in tRNA$^{Ser}$**

<u>DNase treatment and demethylation</u>

Total RNA (5 ug) was treated with 1 uL of TURBO™ DNase (2 U/uL) (Invitrogen™ AM2238) in a 50 uL reaction volume at 37 °C for 30 min to remove any genomic DNA contaminations. Total RNA was then purified using the RNA Clean & Concentrator-5 kit (Zymo Research, R1013) following the manufacturer's protocol. The demethylation assay was performed as described above with modifications for total RNA. Briefly, in a reaction volume of 100 µL containing 5 µg of total RNA (~40 pmol of tRNA) treated with 1:5 molar ratio of D135S mutant AlkB (200 pmol). The reaction buffer contained 300 mM KCl, 2 mM $MgCl_2$, 50 µM of $(NH4)_2Fe(SO4)_2 \cdot 6H_2O$, 300 µM 2-ketoglutarate (2-KG), 2 mM L-ascorbic acid, 50 µg/mL BSA, and 50 mM MES buffer (pH 5.0). The reaction was incubated for 2 h at room temperature and quenched by the addition of 5 mM EDTA. Total RNA was purified using the RNA Clean & Concentrator-5 kit (Zymo Research, R1013) following the manufacturer's protocol.

NaCNBH$_3$ reduction of RNA

Total RNA samples (2-3 ug) were next treated with the reducing agent sodium cyanoborohydride (100 mM NaCNBH$_3$ in $H_2O$) or water (as a control) in a final reaction volume of 100 μL. Reactions were initiated by the addition of 1 M HCl to a final concentration of 100 mM and incubated for 20 min at room temperature. Reactions were stopped by neutralizing the pH by the addition of 30 μL 1 M tris−HCl pH 8.0. The quenched reactions were adjusted to 200 μL with H2O, purified via ethanol. precipitation, and washed via 70% ethanol. The pelleted RNA was dried using a Speedvac, resuspended in $H_2O$, and quantified using a Nanodrop 2000 spectrophotometer. Samples were stored at −20 °C until reverse transcription was performed.

RT and PCR of tRNA$^{Ser}$

An amount of ~200−500 ng of purified RNA from individual reactions (NaCNBH$_4$ treated or water control) were mixed with 4.0 pmol of the reverse (RT) primer for tRNA$^{Ser}$ AGA/TGA (Table S1) in 1x First Strand reaction buffer (SuperScript™ III Reverse Transcriptase kit from Invitrogen™ 18080093) to a final volume of 20 μL. The annealing reaction was done by heating to 95 °C for 1 min followed by 65 °C for 5 min and then transferring to ice for 1 min. After annealing, reverse transcription was performed with SuperScript™ III (Invitrogen, 18080093) enzyme by adding 5 mM DTT, 25 units of RNasin, 100 units of SuperScript™ III enzyme, and 500 μM dNTPs (5 mM GTP, 10 mM CTP, 10 mM ATP, and 10 mM TTP). The reactions were incubated at 55 °C for 60 min. Samples were quenched by increasing the temperature to 70 °C for 15 min.

The cDNA products from the reactions and controls were then PCR amplified. PCR reactions were set up with 2 μL of the RT reaction in a 50 μL volume with 1 unit of Phusion® High-Fidelity DNA Polymerase (New England Biolabs M0530), 1x HF buffer, 2.5 pmol each forward and reverse PCR primers for tRNA$^{Ser}$ AGA/TGA (Table S1), 200 μM each dNTP. Thermocycling conditions were as below.

| | | |
|---|---|---|
| Initial denaturation | 95 °C | 1 min |

| | | |
|---|---|---|
| 33 Cycles | 95 °C<br>Annealing temperature 67 °C<br>72 °C | 0.15 min<br>0.30 min<br>0.30 min |
| Final extension | 72 °C | 7 min |
| Hold | 4 °C | Infinity |

PCR products were run on a 2% agarose gel, stained with SYBR safe (Thermo Fisher, S33102), and visualized on a UV transilluminator at 302 nm. Bands of the desired size were excised from the gel. DNA was extracted using NucleoSpin Gel and PCR Clean-up kit (Macherey Nagel, 740609.50) and submitted for Sanger sequencing (Genewiz) with the forward sequencing primer for $tRNA^{Ser}$ AGA/TGA (Table S1). Processed sequencing traces were viewed using the SnapGene software. The peak height for each base was measured, and the percent misincorporation was determined using the equation: "Percent Misincorporation = (peak intensity of T)/ (sum of C and T base peaks) x 100%". Final misincorporation values were determined by subtracting the background water control misincorporation levels from that of the corresponding reactions.

**$Ac^4C$-Seq**

HEK-293T (ATCC, CRL-3216) and 3T3 cells (ATCC, CRL-1658) were cultured in 10 cm plates in DMEM with 10% FBS, penicillin (100 U/ml) and streptomycin (100 g/ml) in 37 °C as described above. S. cerevisiae strain BY4741 was harvested in mid-log phase after growth at 30 °C in standard YPD medium (1% yeast extract, 2% Bacto Peptone, 2% dextrose). Total RNA from HEK293T and 3T3 cells was isolated using TRIzol as described above. Total RNA was isolated from yeast using the MasterPure Complete RNA Purification Kit (VWR, MC85200) according to the manufacturer's protocol. RNA concentrations and purity were measured by nanodrop and RNA stored in pure water at -80C prior to library construction.

Library construction

Uncharging of tRNAs was performed as previously described in QuantM-tRNA-seq.[17] Total RNA (1 µg) for each sample was uncharged by incubating in 20 mM Tris–HCl (pH 9.0) at 37 °C for 45 min and subsequently neutralized by the addition of an equal volume of 20 mM sodium acetate/acetic acid (pH 4.8) with 20 mM NaCl. The uncharged total RNA samples were next size-selected on a column according to the manufacturer's protocol to maintain RNAs <200nt (RNA Clean and Concentrator-5 kit, Zymo Research, R1013). The size-selected RNA was next dephosphorylated and ligated with a 3' RNA oligo containing an internal barcode, a 3nt unique molecular index (UMI), and the Truseq R2 Illumina adapter. The barcoded samples could be pooled as described in RNAtag-Seq protocol[18] with a 7nt barcode and 3nt UMI. Samples were next treated with either $NaCNBH_3$ or a mock-treatment of water as described in the $ac^4C$-seq protocol[19]. Reverse transcription was performed using TGIRT at 42 °C for 16 h[20]. RNA was hydrolyzed and a second ligation was performed by adding a 3' DNA oligo (5' to the RNA strand) containing the Truseq R1 Illumina adapter and a 6bp UMI for BY4741/HEK-293T/3T3 or without a UMI for mouse liver samples. Barcoded primers were then used to amplify the sequencing library via PCR. Libraries were subsequently sequenced on Illumina NovaSeq 6000 platform with an SP100 kit with read-lengths split evenly between R1 and R2.

Processing of reads, alignment, and analysis

Cutadapt (v4.2)[1] command [cutadapt -a AGATCGGAAGAGCACACGTCTGAAC -A AGATCGGAAGAGCGTCGTGTAGGGA -m 20] was applied to paired-end parental fastq files prior to merging of reads with BBMap (v38.90)[2] command [bbmerge.sh] using default settings. Merged reads were deduplicated at the sequence level based on uniqueness of the sequence present in the merged fastq files including a UMI using BBMap command [dedupe.sh] and default settings[21]. Alignments of the merged & deduplicated reads were performed according to the bowtie2 (v2.3.5.1) parameters simplified to [bowtie2-align-s -k 100 --very-sensitive --ignore-quals --np 5 --reorder] and passed to the pipeline described in[22] using reference genomes sacCer3, mm10, and hg38 with annotations from the Genomic tRNA Database.[23] Consolidated pileup tables of the aligned bam files were created by piping the output of 'samtools mpileup' into cpup (https://github.com/y9c/cpup) and analysis performed to determine the cytidine to thymine misincorporation rates using custom R scripts.

**RNA-seq**

Total RNA from Thumpd1 WT and KO HEK-293T cells and mouse tissues (liver and cerebellum) were extracted as described above. All the samples (n=4 for mouse tissues and n=3 for HEK-293T) were pooled and sequenced on NovaSeq 6000 S1 using Illumina® Stranded Total RNA Prep,

Ligation with Ribo-Zero Plus and paired-end sequencing. Briefly, RNA-Seq FASTQ files were aligned to the reference genome using STAR[14] and raw counts data produced using RSEM.[24] HEK-293T samples were aligned to GRCh38 using the GENCODE_46 GTF annotation, while mouse tissue samples were aligned to mm10 using the GENCODE_M21 GTF annotation. Downstream analysis and visualization were performed within the NIH Integrated Analysis Platform (NIDAP) using R programs developed by a team of NCI bioinformaticians on the Foundry platform (Palantir Technologies). The RSEM counts matrices were filtered for low counts (<1 cpm) and normalized by quantile normalization using the limma package.[25] Differentially expressed genes were calculated using limma-Voom.[26] GSEA was performed using fgsea package.[27]

**Proteomic analysis**

Lysis, Digestion, and TMTpro labeling

Each cell pellet was lysed in 500µL EasyPep Lysis buffer (Thermo Fisher, PN A45735) and treated with 2 µL universal nuclease (Thermo, PN 88700). Protein concentration was determined by the BCA method and 100 µg was taken from each condition for digestion. Samples were adjusted to 100 µL total with lysis buffer and treated with 50 µL each of reducing solution and alkylating solution provided with the EasyPep kit (Thermo, A40006). Incubated for 1 h in the dark at 25 °C then made 4 aliquots of 40 µL (20 µg) for each condition and added 60 µL of 13 ng/µL trypsin/LysC (provided with EasyPep kit) and incubated at 37 °C overnight for a total of 19 h at which point 20 µL of 5 µg/µL TMTpro 18-plex label (Thermo, PN A52045) was added to samples and incubated for 1 h at 25 °C. Excess TMTpro was quenched with 20 µL of 5% hydroxylamine, 20% Formic acid for 10 min and samples were then combined. Samples were cleaned using EasyPep mini columns provided with the EasyPep kit as described in the manual. Eluted peptides were dried in speed-vac.

Off-line fractionation and LC/MS analysis of peptides

TMTpro labeled peptides were fractionated by High-pH reverse phase liquid chromatography using a Waters Acquity UPLC system with a fluorescence detector (Waters, Milford, MA) using a 150mm x 3.0mm Xbridge Peptide BEMTM 2.5 µm C18 column (Waters, MA) operating at 0.35 mL/min. The dried peptides were reconstituted in 50 µL of mobile phase A (10 mM ammonium formate, pH 9.3) and eluted from the column in mobile phase B (10 mM Ammonium Formate, 90% ACN, Thermo Fisher Scientific). The peptides were eluted using gradient elution of 10 – 50% phase B (1.5 – 60 min) followed by 50 – 90% phase B (60 – 65 min). Sixty-five fractions were collected, and the fractions were then consolidated into 12 pools based on the chromatogram intensity and vacuum centrifuged to dryness. Each fraction was resuspended in 50 µL of 0.1% FA and 10 µL was analyzed using a Dionex U3000 RSLC in front of a Orbitrap Eclipse (Thermo) equipped with an EasySpray ion source. Solvent A consisted of 0.1%FA in water and Solvent B consisted of 0.1%FA in 80%ACN. Loading pump consisted of Solvent A and was operated at 7 µL/min for the first 6 min of the run then dropped to 2 µL/min when the valve was switched to bring the trap column (Acclaim™ PepMap™ 100 C18 HPLC Column, 3 µm, 75 µm I.D., 2 cm, PN 164535) in-line with the analytical column EasySpray C18 HPLC Column, 2 µm, 75 µm I.D., 25 cm, PN ES902). The gradient pump was operated at a flow rate of 300nL/min. Each run used a linear LC gradient of 5-7%B for 1 min, 7-30%B for 83 min, 30-50%B for 25 min, 50-95%B for 4 min, holding at 95%B for 7 min, then re-equilibration of analytical column at 5%B for 17 min. MS acquisition employed the TopSpeed method with a 3 sec cycle time and the following parameters: Spray voltage was 1800V and ion transfer temperature was 275 ºC. MS1 scans were acquired in the Orbitrap with resolution of 120,000, AGC of 4e5 ions, and max injection time of 50 ms, mass range of 400-1600 m/z; MS2 scans were acquired in the Orbitrap using method with resolution of 50,000, AGC of 1.25e5, max injection time of 86 ms, HCD energy of 38%, isolation width of 0.4Da, intensity threshold of 2.5e4 and charges 2-5 for MS2 selection. Advanced Peak Determination, Monoisotopic Precursor selection (MIPS), and EASY-IC for internal calibration were enabled and dynamic exclusion was set to a count of 1 for 15sec.

Database search and post-processing analysis

MS files were searched together with Proteome Discoverer 2.4 using the Sequest node. Data was searched against the Uniprot Human database from Feb 2020 using a full tryptic digest, 2 max missed cleavages, minimum peptide length of 6 amino acids and maximum peptide length of 40 amino acids, an MS1 mass tolerance of 10 ppm, MS2 mass tolerance of 0.02 Da, fixed modifications for TMTpro (+304.207) on lysine and peptide N-terminus and carbamidomethyl (+57.021) on cysteine and variable oxidation on methionine (+15.995 Da). Percolator was used for FDR analysis and TMTpro reporter ions were quantified using the Reporter Ion Quantifier node and normalized using the total peptide intensities of each channel. The Log2FC (Median of groups) and pvalues (ANOVA) were calculated within the PD2.4 software FDR was set to < 1%. Proteins with p-value <0.05 and Log2FC cutoffs of >0.6 and <-0.6 were considered differentially expressed. TMTpro channel assignment for conditions were as follows: WT replicates □ 126, 127N, 127C, 128N; KO replicates □ 128C, 129N, 129C, 130N; Rescue replicates □ 130C, 134N, 134C, 135N.

**Northern blotting-based analysis of tRNA charging**

Total RNA isolation under acidic conditions

Total RNA for Northern immunoblots was prepared under acidic conditions to preserve aminoacylation essentially as described by Varshney.[28] and applications of this approach by Chernyakov *et al.*[29] WT or *THUMPD1* knockdown HEK-293T cells grown to about 70-75% confluency on 100 mm dishes were quickly washed with cold 0.3 M sodium acetate, 10 mM EDTA, pH 4.5, then 0.25 mL of wash buffer was added to the cells followed by addition of 0.75 mL Trizol reagent (Life Technologies) saturated with wash buffer and thorough pipetting of cell suspension to promote cell lysis. After 5 min, the cell lysate was transferred to a microfuge tube and 0.2 mL of chloroform was added, suspension was vortexed, incubated for 2min to allow for phase separation and centrifuged at 12,000 rcf for 15 min at 4 °C. The aqueous phase was transferred to a new tube and RNA was precipitated by addition of two volumes of cold 100% ethanol, incubating on dry ice for 30 min and centrifugation at max speed for 15 min at 4 °C. RNA pellets were washed with cold 70% ethanol, re-centrifuged, air-dried for 10 min and resuspended in 10 mM sodium acetate, 1 mM EDTA, pH 4.5. RNA concentrations were measured using NanoDrop Microvolume Spectrophotometer (Thermo Fisher Scientific). All RNA samples were prepared in triplicates.

Northern immunoblots analysis of tRNA aminoacylation

RNA samples (10 μg total RNA) prepared under acidic conditions were resolved on 1 mm thick 6.5% polyacrylamide gel (acrylamide to bis-acrylamide ratio 19:1) containing 8 M urea and 0.1 M sodium acetate pH 4.5. Each sample was mixed with an equal volume of acidic RNA loading dye (0.1 M sodium acetate pH 4.5, 8 M urea, 0.05% bromophenol blue, 0.005% xylene cyanol) and ran at 450 V for about 20 h in a cold room. The samples were loaded in triplicates. Deacylated

RNA controls were prepared by incubating the samples in 0.1 M Tris-HCl pH 9.0 and 1 mM EDTA for 30 min at 37 °C followed by ethanol precipitation and dissolving the RNA pellets in 10 mM sodium acetate, 1 mM EDTA, pH 4.5. After separation on the gel, the portion of the gel containing tRNA was cut and RNA was transferred to a Hybond $N^+$ nylon membrane (Cytiva) using BioRad Protean II electrophoresis cell in 1×TAE buffer at 15V for 1 h in a cold room. The membrane was rinsed in fresh 1×TAE buffer, air-dried for about 30 min and RNA was UV crosslinked at 120,000 microjoules/$cm^2$ in Stratalinker UV crosslinker (Stratagene). After RNA crosslinking the Nylon membrane was soaked in 2×SSC for 1 min then pre-hybridized in ULTRAhyb™ Ultrasensitive Hybridization Buffer (Ambion) for 1 h at 42 °C using a Hybaid H-9360 hybridization oven followed by hybridization with 50-100 pmol of biotinylated oligonucleotide probe specific to one of the tRNAs of interest (Table S1) for 12 h at 42 °C. The membrane then was washed twice with 2 ×SSC, 0.1% SDS and once with 2 ×SSC for 30 min at 42 °C each time. After that the membrane was soaked for 1 min in 1×PBS containing 0.05% Tween-20 followed by blocking in 5% BSA dissolved in the same buffer for 30 min at RT. Then membrane was incubated with HRP-conjugated streptavidin (BioLegend, 1:1000 dilution in 1×PBS, 0.05% Tween-20, 405210) for 2 h at RT, washed twice with 1×PBS, 0.05% Tween-20 and once with 1×PBS 20 min each time. The membrane was treated for 5 min with Thermo Scientific SuperSignal West Pico PLUS Chemiluminescent Substrate (0.1 mL/$cm^2$) and tRNA bands on membrane were visualized using BioRad ChemiDoc™ XRS+ imager and Image Lab 4.1 software.

**qPCR-based tRNA charging assay**

The tRNA charging assay was performed as previously described by Pavlova *et al*[30]. Briefly, HEK-293T THUMPD1 WT and KO cells were grown in DMEM media as described above and washed with cold PBS and lysed by adding 1 mL TRIzol (Thermo Fisher, 15596026) to the plate and incubating for 5 min on ice. Lysates were then collected and mixed with 200 μL chloroform in an Eppendorf tube by shaking vigorously for 20 sec. After centrifuging for 15 min at 18,600 rcf at 4 °C, top fraction was collected, and the total RNA was precipitated with 2.7x volumes of cold ethanol in the presence of 2 μL of GlycoBlue Coprecipitant (ThermoFisher, AM9515) overnight at 4 °C. Pellet was resuspended in 300 μL of tRNA precipitation buffer (0.3 M acetate buffer, pH = 4.5, 10 mM EDTA) and pelleted again by adding 2.7x volumes of cold ethanol, incubating at -20 °C overnight, and centrifuging for 30 min at 18,600 rcf at 4 °C. Pellet was washed with 80% ethanol and resuspended in 32 uL tRNA resuspension buffer (10 mM acetate buffer pH = 4.5, 1 mM EDTA) before determining the concentration via Nanodrop. For the oxidation treatment, 2 µg of RNA from each sample was used. Oxidization reactions were done by treating the RNA with 0.2 M $NaIO_4$ in sodium acetate buffer, pH 4.5 for 20 min at room temperature in the dark. For the controls, 0.2 M NaCl was used. Reactions were quenched by adding 2.2 µL of 2.5 M glucose and incubating for 15 min at room temperature in the dark. Into each glucose-quenched reactions, 1 µL of yeast Phe tRNA (Sigma, R4018) solution (7 ng/µL stock in water) was added to serve as a spike-in control and the RNA were precipitated with ethanol as previously described. Next, to facilitate the deacylation, pelleted RNA was resuspended in 100 µL of 50 mM Tris, pH 9 and incubated at 37 °C for 45 min. The reactions were quenched with 100 µL of tRNA quench buffer (50 mM Na Acetate buffer, pH = 4.5, 100 mM NaCl) and RNA was pelleted by ethanol precipitation. The pellet was resuspended in 10 µl of water and the concentration was measured using the nanodrop. Concentrations in all the tubes were adjusted to the same level by adding water. Next, the adapter ligation step was carried out with a 5′-adenylated DNA adaptor (5′-/5rApp/TGGAATTCTCGGGTGCCAAGG/3ddC /- 3′), ~380 ng of RNA, using T4 RNA ligase 2, truncated KQ (NEB, M0373) at 18 °C, overnight. Then the CSQ_RT primer (5'GCTGCCTTGGCACCCGAGAATTCCA3') was annealed (30 sec at 90 °C, 5 min at 65 °C, immediately put on ice for 1 min) to the adapter region of tRNA before carrying out the reverse transcription reaction with SuperScript RT IV (Thermo, 18090050) according to the manufacturer's protocol. The synthesized cDNA was diluted (1:10) and 2.5 uL of each reaction was used to set up each qPCR with tRNA isodecoder-specific primers using 2x SYBR Green mix (Life Technologies, 4368702). The forward primer matched the 5′ end of the tRNA, and the reverse primer covered the junction between the 3′ end of the tRNA and the ligated adaptor. The primer pairs used are listed in Table S1.

LightCycler® 480 II PCR instrument (Roche) instrument was used for the qPCR step with the following thermocycling protocol:

| Cycle step | Temperature | Time | Cycle |
|---|---|---|---|
| Reverse transcription | 55 °C | 10 min | 1 |
| Initial denaturation | 95 °C | 1 min | 1 |

| | | | |
|---|---|---|---|
| Denaturation<br>Extention | 95 °C<br>60 °C | 10s<br>30s | 45 |
| Melt curve | 60-95 °C | various | 1 |

The cycle threshold values (Ct) were obtained from the LightCycler 480 SW 1.5.1 software. Average Ct value for each reaction and control was calculated from two technical qPCR replicates for each of the three biological replicates. Finally, the charged tRNA fraction was calculated by normalizing average ${}^{tRNA}$Ct to the yeast-Phe tRNA using the following formula.

$$\Delta\Delta Ct = (\Delta Ct\ Leu - \Delta Ct\ yphe)rxn - (\Delta Ct\ Leu - \Delta Ct\ yphe)con$$

$$charged\ fraction = 2^{-(\Delta\Delta Ct)}$$

**Analysis of mTOR activation in THUMPD1 KO cells**

HEK-293T THUMPD1 WT/KO cells were plated ($3x10^6$ cells) into 100 mm tissue culture plates with 10 mL of complete medium (10% FBS, DMEM, 1% P/S) and allowed to adhere for 48 hours. Next, cells were starved with DMEM lacking amino acids (Thermo Fisher Scientific catalog #ME120086L1) for 1 hour and/or re-fed with complete medium for 1 hour. For inhibitor treatments, medium was aspirated out and 10 mL of DMEM complete medium with DMSO, DMSO + Rapamycin (10 nM), or DMSO + AZD2014 (1µM) was replaced for 1 hour. After incubation cells were collected with a cell scraper and lysed in cold NP-40 lysis buffer (50 mM tris, 150 mM NaCl, and 1.0% NP-40 at pH 8.0). Protein lysate (25 µg) was separated in Novex WedgeWell 10-20% tris-glycine gels (Thermo Fisher Scientific, XP10202BOX) for proteins smaller than 100 kDa or 3-8% tris-acetate gels (Thermo Fisher Scientific, EA03785BOX) for proteins larger than 100 kDa. Proteins were transferred to polyvinylidene difluoride membranes (Bio-rad, 1620177), incubated with primary antibodies overnight at 4 °C, and then incubated with secondary antibodies at room temperature for 1 h in 1x TBST + 5% blocking (Bio-rad, 1706404). Membranes were incubated with SuperSignal West Pico Plus Chemiluminescent Substrate (Thermo Fisher Scientific, 34578). Primary antibodies were as follows: mouse monoclonal custom antibodies targeting endogenous mEAK-7 were generated by Genscript (unpublished). All antibodies from Bethyl Laboratories were as follows: Raptor (A300-553A [RRID: AB_2130793]), Rictor (A300-459A [RRID: AB_2179967]). All antibodies from CST were as follows: ($Thr^{389}$) p-p70 S6 kinase (9234S [RRID: AB_2269803]), S6K1 (2708S [RRID: AB_390722]), ($Ser^{473}$) p-AKT (4058S [RRID: AB_331168]), total AKT (4685S [RRID: AB_10698888]), ($Ser^{235/236}$) p-S6 ribosomal protein (4858S [RRID: AB_916156]), ($Ser^{240/244}$) p-S6 ribosomal protein (2215S [RRID: AB_331682]), S6 ribosomal protein (2217S [RRID: AB_331355]), ($Thr^{37/46}$) p–4E-BP1 (9459S [RRID: AB_330985]), ($Ser^{65}$) p–4E-BP1 (9451S [RRID: AB_330947]), 4E-BP1 (9452S [RRID: AB_331692]), GAPDH (2118S [RRID: AB_561053]). Antibodies p-S6, S6, and 4E-BP1 were used at 1:3000 dilution and the remainder at 1:1000 dilution in 5% bovine serum albumin (BSA) in 1× tris-buffered saline with Tween-20 (TBST) buffer with 0.04% sodium azide. Secondary antibodies for immunoblot analysis: 1:10,000 dilution for an α-mouse IgG horseradish peroxidase (HRP) conjugate (Promega, W4021 [RRID: AB_430834]) and 1:20,000 dilution for an α-rabbit IgG HRP conjugate (Promega, W4011 [RRID: AB_430833]).

**Data and software availability**

Generated high-throughput sequencing datasets are publicly available in the NCBI's Gene Expression Omnibus (GEO) under accession numbers GSE272399 and GSE272400. Proteomic data is available via MassIVE under accession number MSV000094896 using the reviewer login MSV000094896_reviewer and password THUMPD1.

**Full gel images**

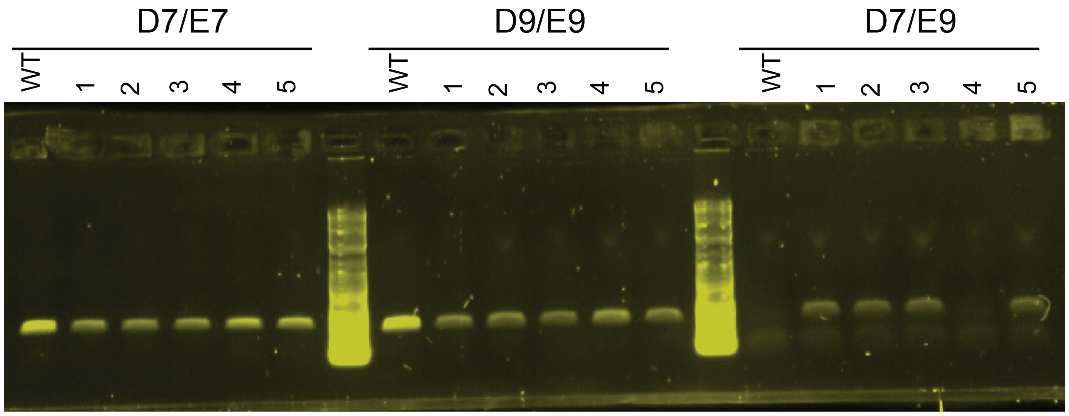


Full agarose gel image for PCR-based genotyping gel pertaining to Fig 1c. Lanes labeled as WT, 1, and 2 from each primer pair is shown in Fig 1c.

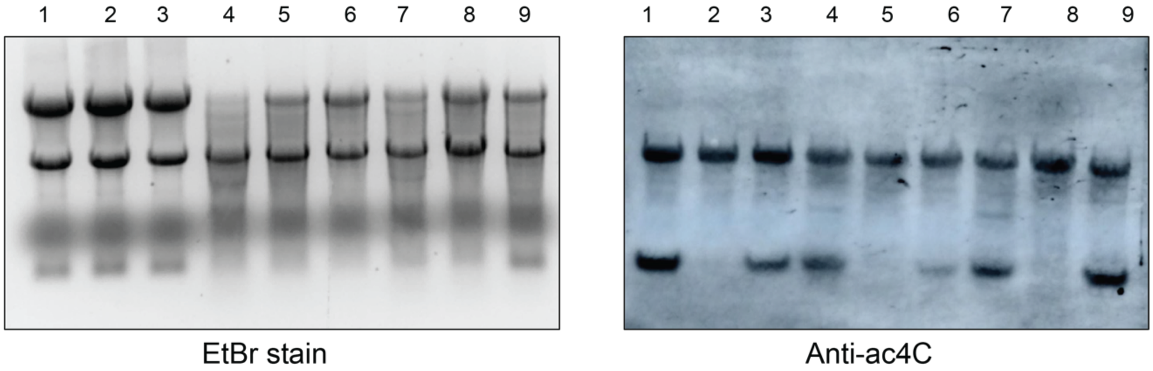


Full immuno-Northen blot and the Ethidium bromide-stained image pertaining to Fig 1d and Fig 3b. Lanes 4 and 5 correspond to data from mouse Thumpd1 WT and KO RNA shown in Fig 1d. Lanes 1 and 2 correspond to data from HEK-293T THUMPD1 WT and KO RNA shown in Fig 3b.

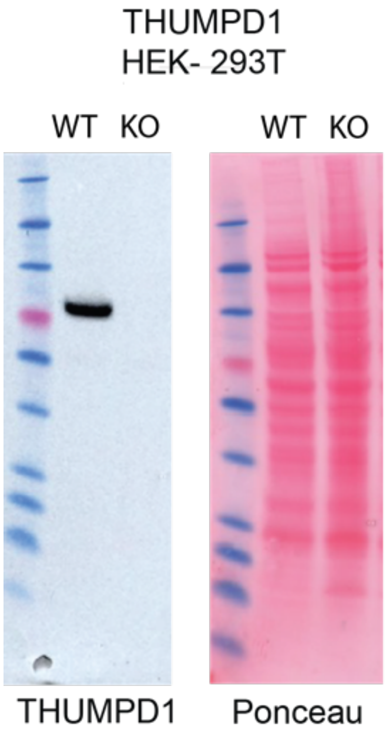


Full western blots pertaining to Fig 3a.

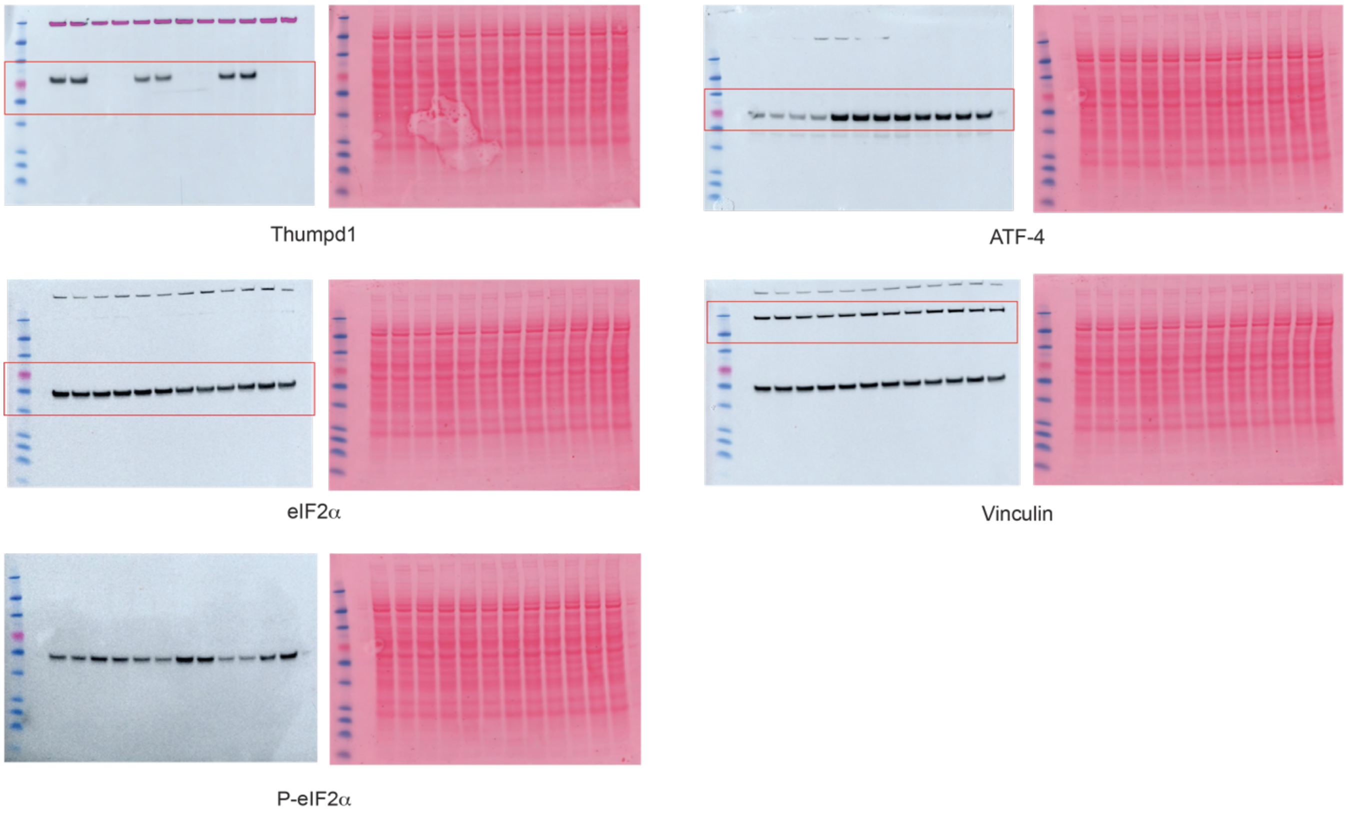


Full Western blots pertaining to Fig 4e.

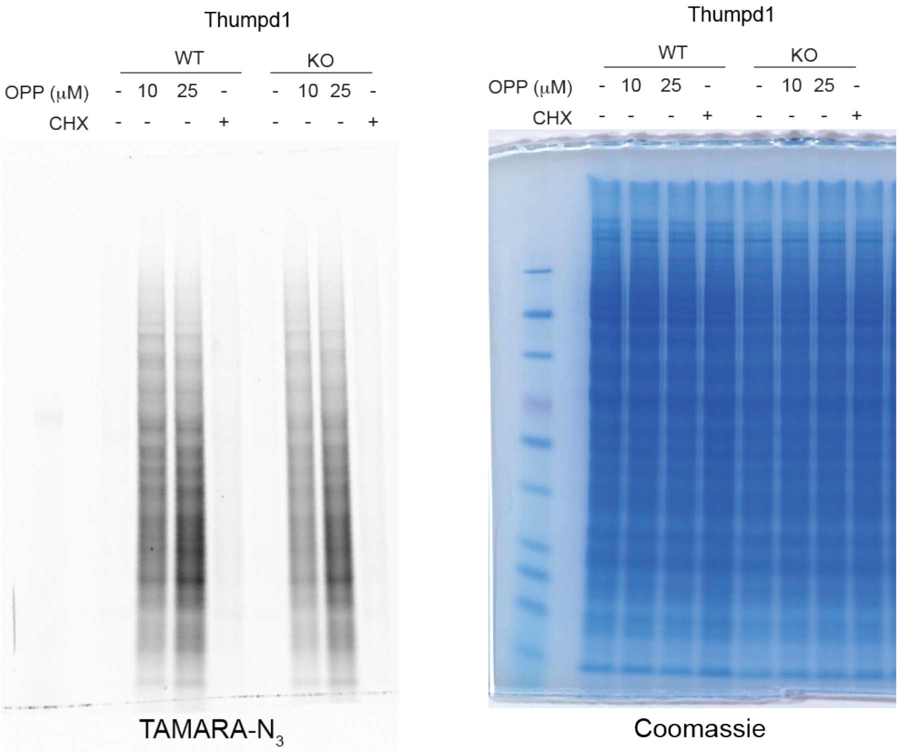


Full SDS-PAGE gel pertaining to Fig 4f.

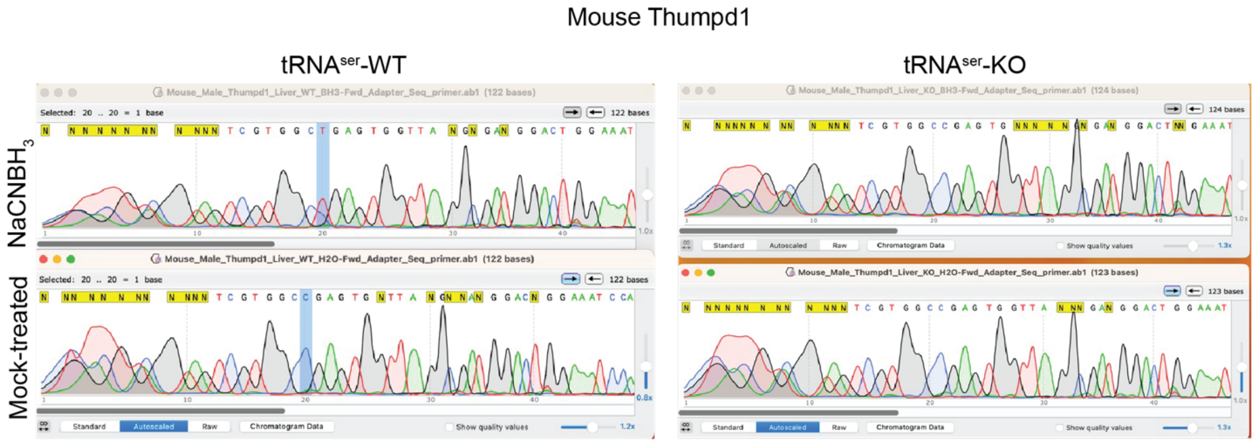


Full Sanger traces pertaining to Fig S1f.

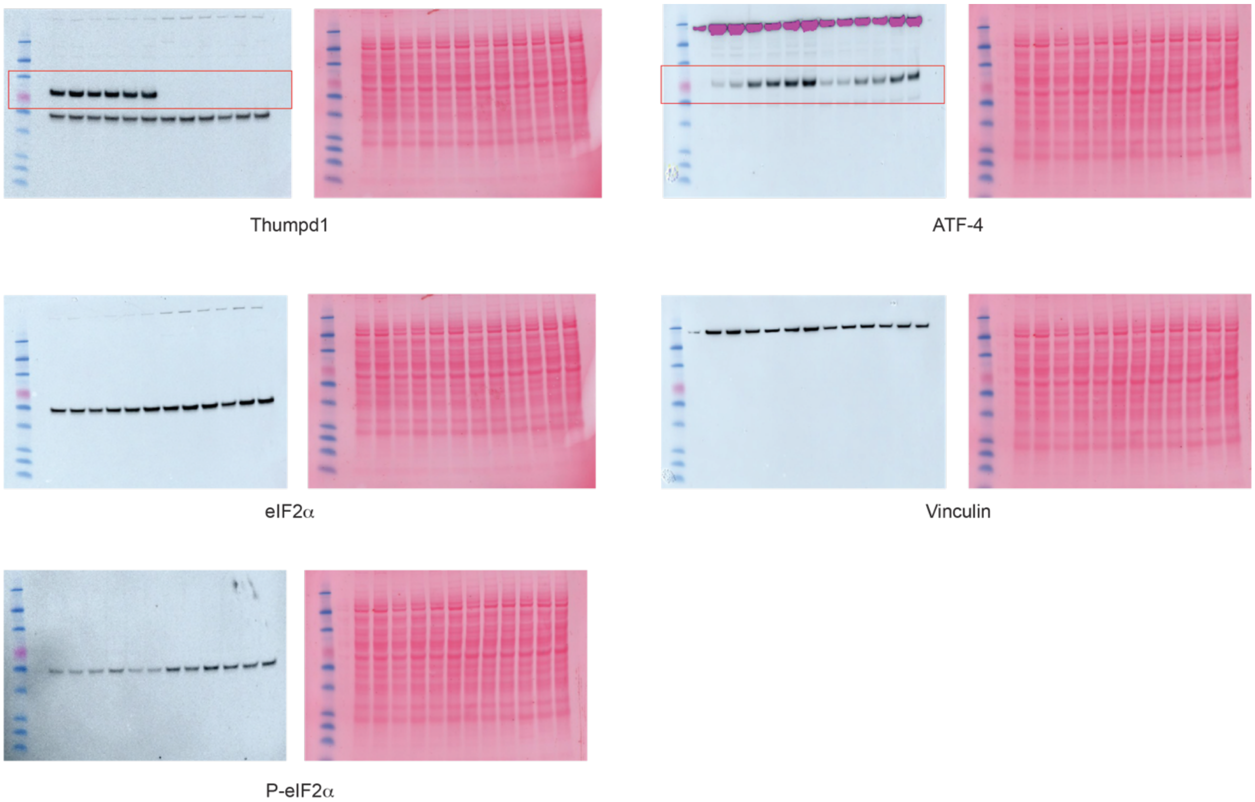


Full Western blots pertaining to Fig S4c.

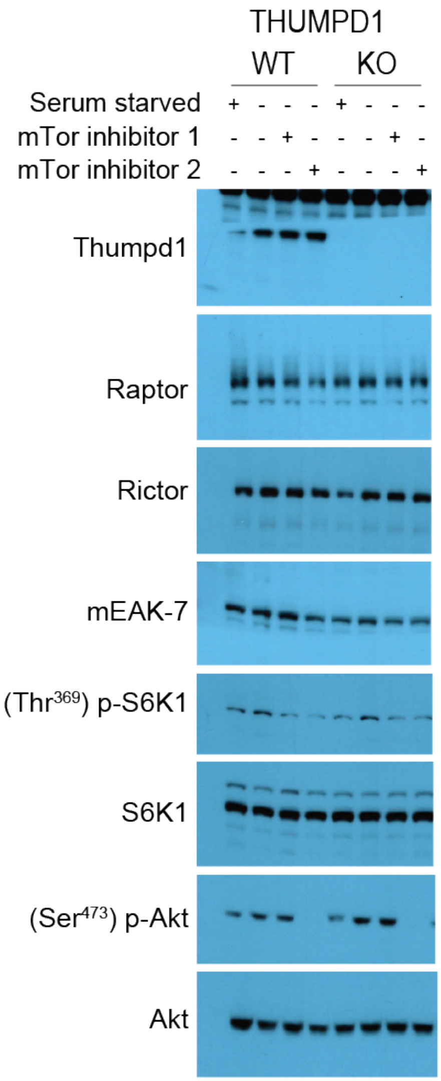


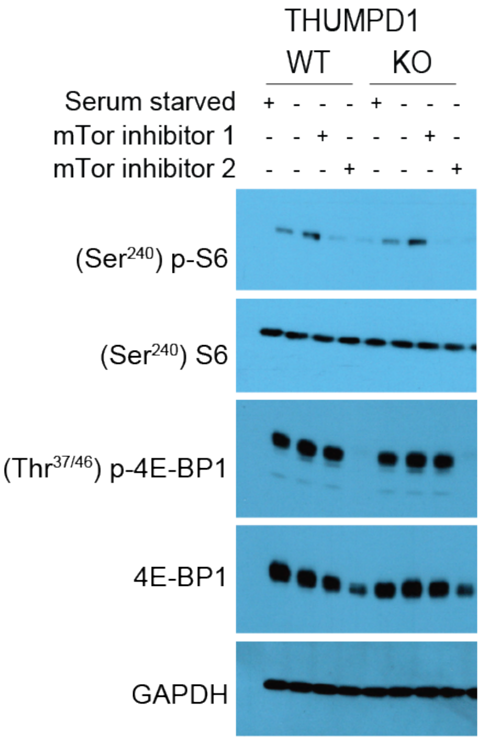


Autoradiographs pertaining to Fig S4e.

**Table of oligonucleotides used in this study, related to the Methods section**

| **NAME** | **SEQUENCE (5` -> 3`)** |
|---|---|
| **Guide RNAs for generation of *Thumpd1*$^{+/-}$ mice** | |
| Guide 1 | CAGAGAGUAUCGAGGAGCUC |
| Guide 2 | CCUGUUGCCGGAAAGCGCAA |
| **Oligonucleotide PCR primers for mouse genotyping** | |
| D9 PCR primer | GCTCGTTCATGTTGCAGGTG |
| E9 PCR primer | CAAACTGTCGCGTACGTGTG |
| D7 PCR primer | TGAGACCACCTTCCCACAGA |
| E7 PCR primer | GAGTGCAAAAGACTCGCAGC |
| **Oligonucleotides for mim-tRNAseq library preparation** | |
| oBZ407 | /5Phos/AppNNNNNNCACTCGGGCACCAAGGA/3ddC |
| RT primer oBZ408 | /5Phos/ RNNNAGATCGGAAGAGCGTCGTGTAGGGAAAGAGTGTAGATCTCGGTGGTCGC/iSP18/TTCAGACGTGTGCTCTTCCGATCTGTCCTTGGTGCCCGAGTG |
| PCR primer I | AATGATACGGCGACCACCGAGATCTACAC |
| PCR primer II | CAAGCAGAAGACGGCATACGAGAT[8nt_barcode]GTGACTGGAGTTCAGACGTGTGCTCTTCCG |
| **Oligonucleotides for ribosome and disome profiling** | |
| 3` adapter | NNNNNNCACTCGGGCACCAAGGA |
| RT primer | RNNNAGATCGGAAGAGCGTCGTGTAGGGAAAGAGTGTAGATCTCGGTGGTCGC/iSP18/TTCAGACGTGTGCTCTTCCGATCTGTCCTTGGTGCCCGAGTG |
| **Oligonucleotides used for the Leucine and serine repeat GFP reporter assays** | |
| Leu 3xCTG/C codon | CTG CTC CTG ATCGAGTGCCGCATCAC |
| Leu 3xCTG/C codon | CAG GAG CAG CTTCATGGCGGGCATGG |

| | |
|---|---|
| Leu 3xCTA codons | CTA CTA CTA ATCGAGTGCCGCATCAC |
| Leu 3xCTA codons | TAG TAG TAG CTTCATGGCGGGCATGG |
| Leu 3xTTA codon | TTA TTA TTA ATCGAGTGCCGCATCAC |
| Leu 3xTTA codon | TAA TAA TAA CTTCATGGCGGGCATGG |
| Ser 3xTCG-codon | TCG TCG TCG ATCGAGTGCCGCATCAC |
| Ser 3xTCG-codon | CGA CGA CGA CTTCATGGCGGGCATGG |
| Sequencing oligo for insertional mutant screening (CMV promoter) | CGCAAATGGGCGGTAGGCGTG |
| **Oligonucleotides for RT-qPCR based analysis of Thumpd1 mRNA** | |
| Thumpd1_fwd | GAGAAGAGGCTGAGAAGATTCC |
| Thumpd1_rev | ATCCTGGAGAATATGATGCACTAA |
| **Oligonucleotides for sanger sequencing analysis of $ac^4C$ in $tRNA^{Ser}$** | |
| RT primer $tRNA^{Ser}$ AGA/TGA | TGGCGTAGTCGGCAGGATTC |
| Forward primer $tRNA^{Ser}$ AGA/TGA | GACTGGAGTTCAGACGTGTGCTCTTCCGAT CTGTACGGTAGTCGTGG |
| Reverse primer $tRNA^{Ser}$ AGA/TGA | TGGCGTAGTCGGCAGGATTC |
| Forward sequencing primer | GACTGGAGTTCAGACGTG |
| **Northern immunoblots analysis of tRNA aminoacylation** | |
| tRNA-Leu-UAA-1-1 biotinylated probe | /5Biosg/TGGTACCAGGAGTGGGGTTCGAA |
| tRNA-Leu-CGA-1 biotinylated probe | /5Biosg/TCGTGTCAGGAGTGGGAT |
| **qPCR-based tRNA charging assay** | |
| 5′-adenylated DNA adaptor | /5rApp/TGGAATTCTCGGGTGCCAAGG/3ddC / |
| CSQ_RT primer | GCTGCCTTGGCACCCGAGAATTCCA |
| Leu WAG (AAG/TAG) Fwd | GGTAGYGTGGCCGAGCG |
| Leu WAG (AAG/TAG) Rev | GAGAATTCCATGGCAGYGGTGGG |
| Leu CAA/CAG Fwd | CAGGATGGCCGAGYGGTCTAAGGC |
| Leu CAA/CAG Rev | GAGAATTCCATGGTGTCAGRAGTGGG |
| Ser AGA/TGA Fwd | GTAGTCGTGGCCGAGTGGTTAAG |
| Ser AGA/TGA Rev | GAGAATTCCATGGCGTAGYCGGC |

| | |
|---|---|
| Yeast phe (spike-in) Fwd | GCGGAYTTAGCTCAGTTGGGAGAG |
| Yeast phe (spike-in) Rev | GAGAATTCCATGGTGCGAAYTCTGTGG |
